# Exploring Disordered Regions of Human Spliceosome Proteins

**DOI:** 10.64898/2026.01.07.698225

**Authors:** Bruno de Paula Oliveira Santos, Krishnendu Bera, Luca Grisanti, Isabella Caterina Felli, Roberta Pierattelli, Alessandra Magistrato

## Abstract

Introns are removed from messenger RNAs by the spliceosome, a protein-RNA machinery enriched with intrinsically disordered regions (IDRs). Lacking stable 3D structures, IDRs can adopt diverse conformations interlacing proteins and RNAs components of the spliceosome and regulating splicing. Here we performed a comprehensive bioinformatics analysis of the human spliceosome proteome revealing that many proteins contain over 40% of disordered residues. Spliceosome IDRs are mainly driven by compositional bias due to an excess of charged and RS-like sequences, with both the nature and extent of this disorder being broadly conserved evolutionarily. Additionally, these IDRs are frequent targets of post-translational modifications, especially phosphorylation, and are hotspots for cancer-associated mutations, implicated in different cancer types. Our results collectively underscore central role of IDRs in splicing regulation and disease.

**TOC Graphic:** 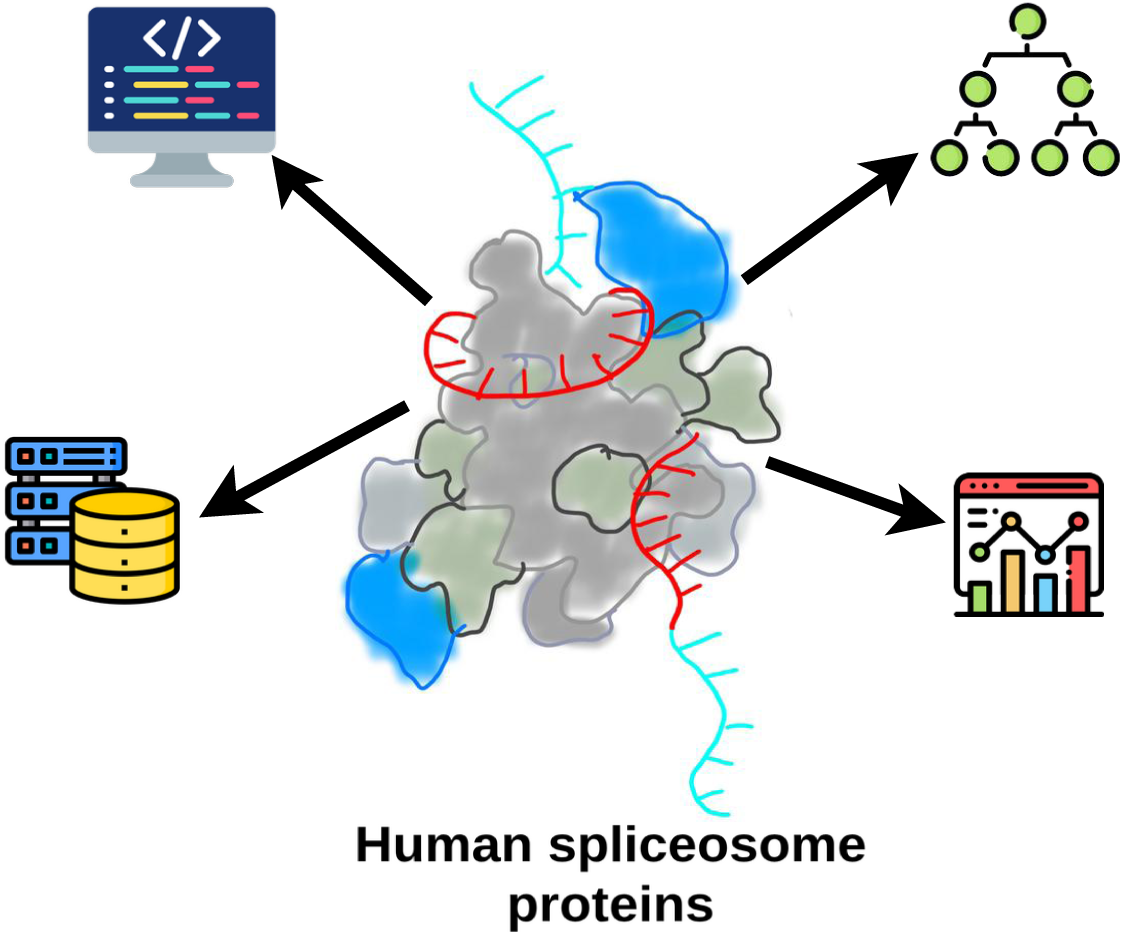

## 1 Introduction

Intrinsically disordered proteins (IDPs) or proteins with intrinsically disordered regions (IDRs) lack regular three-dimensional structure and are characterized by a large set of dynamic and interconverting conformations. IDRs are widespread in the proteome and are present across all domains of life (1). They play critical roles in cellular functions, such as transcriptional regulation, signal transduction, and subcellular organization. The plasticity and structural heterogeneity of IDRs indeed expand their repertoire of macromolecular interactions and allow them to be finely modulated by their structural and chemical environment (2–4). IDRs are also abundant in ribonucleoprotein complexes involved in gene expression, regulation and synthesis (5). Among them, the spliceosome machinery, which promotes premature messenger RNA (pre-mRNA) splicing via dynamic protein/RNA binding and dissociation events, contains a large fraction of IDRs in its components (6).

In eukaryotic cells, the spliceosome removes the noncoding regions (introns) from a pre-mRNA transcript and connects the coding sequences (exons) (7). The most prevalent spliceosome form, the major spliceosome, consists of five small nuclear RNA (snRNA) strands (U1, U2, U4, U5 and U6) and 100-200 associated proteins, which assemble into small nuclear ribonucleoprotein particles (snRNPs) (8,9). The ordered and regulated assembly of these snRNPs and auxiliary proteins, form the spliceosome complex that undergoes extensive rearrangements during the splicing process. The resulting major spliceosome, which splices most introns (U2-dependent introns) uses the U1 and U2 snRNPs to scan pre-mRNA and identify specific intron sequences (splice sites). Namely, U1 and U2 snRNPs bind sequentially to the pre-mRNA—initially forming the E complex, and later A complexes, where the initial recognition and assembly steps occur. In the subsequent steps of the cycle, the spliceosome remodels to perform catalysis, assembling into the B, B^act^, B* complexes (the latter being where the branching reaction occurs) (10). This is followed by the formation of the C and C* complex (where the exon-ligation step occurs) (11), the P complex, where the products are formed, and, finally, the ILS complex, where the intron and the exon are released (12) and the spliceosome is dismantled to undergo a new splicing cycle (4,13,14). (15). Besides those contained in snRNPs, additional proteins are involved in splicing. They can function individually or assemble into multiprotein auxiliary complexes, such as the Nineteen Complex (NTC) complex (16), the exon-junction complex (EJC) (17), the cap-binding complex (CBC) (18), the retention-and-splicing complex (RES) (19), and the transcription-export complex (TREX) (20).

The second form of spliceosome, the minor complex, splices a small class (<1%) of introns (U12-dependent introns), which have stronger splice site consensus sequences as compared to U2-dependent introns. This spliceosome variant consists of four unique snRNAs (U11, U12, U4atac and U6atac) and 14 unique protein factors (21). They work together with the core protein components shared by both spliceosomes, likely through conserved mechanisms.

Irrespectively of the spliceosome type, mutual recognition of spliceosome subunits, correct assembly and components remodeling are paramount for splicing fidelity. In this context, IDRs play a key role in spliceosome function by forming weak interactions that interlace many protein/RNA components and modulate binding dynamics. Moreover, IDRs can be finely regulated by post-translational modifications (PTMs), which readily alter their set of interactions.

Spliceosome IDRs can be divided into four categories: (i) regions containing predicted secondary structure (SS) elements (referred to as SS-IDR); (ii) long (≥25 residues) compositionally-biased IDRs (referred to as CB-IDRs), which includes RS-like, poly-P/Q, G-rich regions, charged disorder and non-charged disorder (5); (iii) IDRs object of PTMs (PTM-IDR) and (iv) of cancer associated mutations (CA-IDR).

To explore their abundance, types, functional and regulatory roles, we performed an integrative sequence-based analysis of spliceosome proteins, focusing on their IDR content, complemented by statistical correlation analysis of PTM and cancer-associated sites, and phylogenetic inferences. Overall, this study deepens our understanding of IDRs in splicing, highlighting their implications in splicing regulation and disease.

## 2 Results

To characterize the prevalence of intrinsic disorder in spliceosome proteins, we performed a bioinformatic analysis across different splicing steps, protein classes and sequence contexts (Figures 1-3, Supplementary Method Section 1.1-1.4 of the Supporting Information for details). To this end, we retrieved the spliceosome sequences from UniProt database (www.uniprot.org), predicted their disorder content and calculated the disorder percentage in different groups.

**Figure 1.**
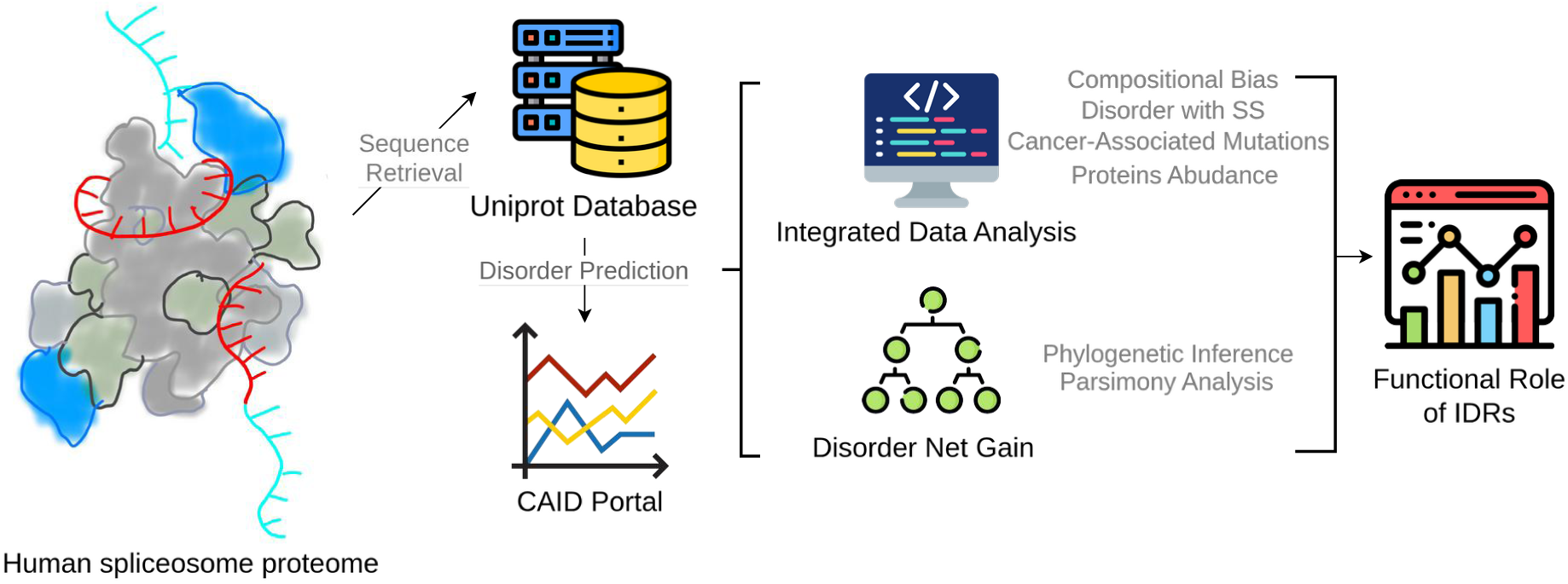
Pipeline of bioinformatic analysis of spliceosome IDRs. Bioinformatic analysis began with sequences retrieval from UniProt database, followed by intrinsic disorder prediction. Sequence-based and statistical analysis were then performed to investigate the type of disorder (compositional bias, disorder with secondary structure), and the associated features, including post-translational modifications and cancer-associated mutations; and finally with their evolutionary history.

**Figure 2.**
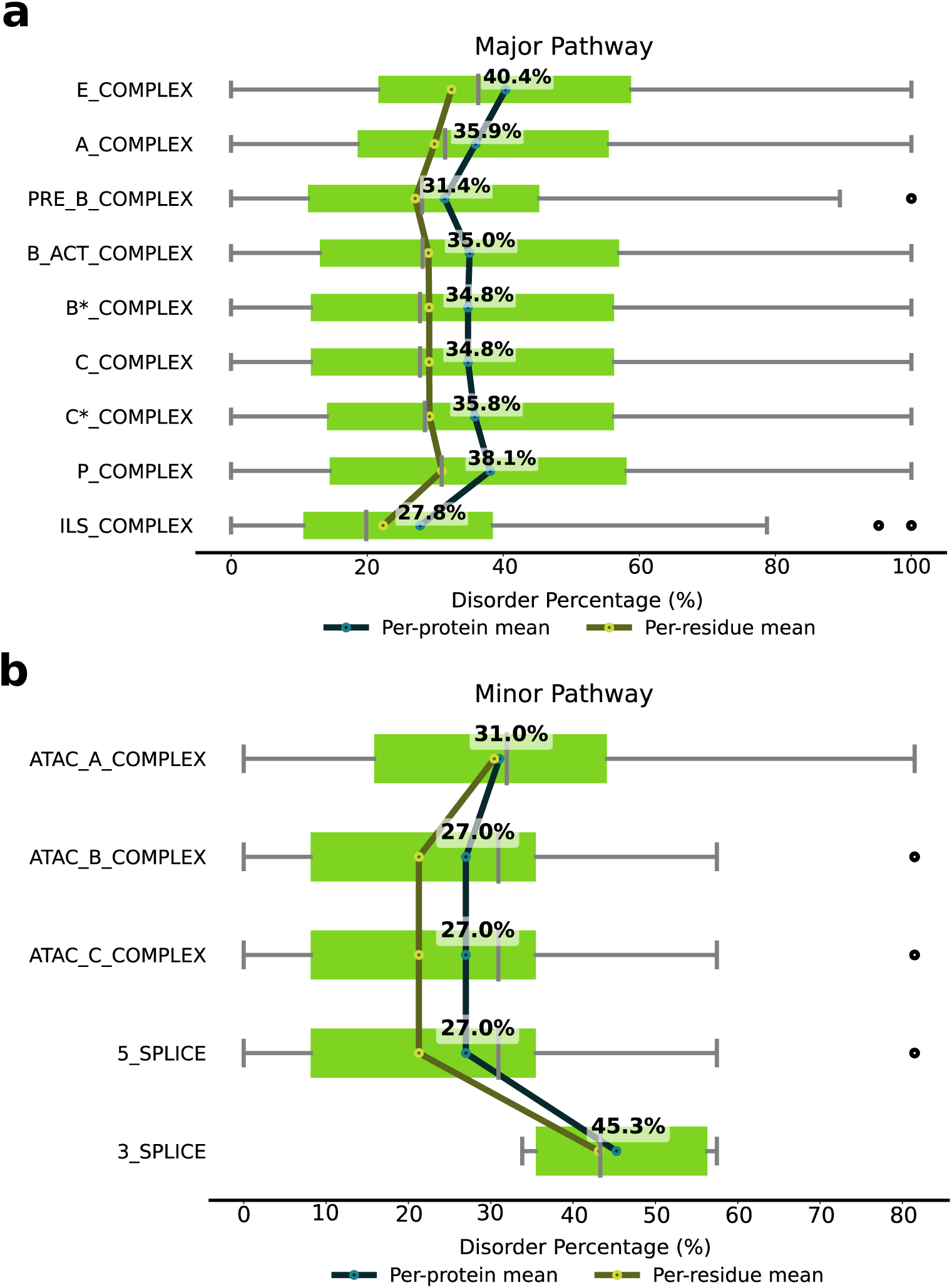
Disorder percentage in spliceosome complexes. Disorder content in complexes belonging to (A) major and (B) minor spliceosome pathways. Per-protein and per-residue mean are reported in blue and yellow dots, respectively. The green bars represent boxplots, showing the distribution of disorder percentages with five key statistical measures: minimum value, first quartile (25 ^th^ percentile), median, third quartile (75^th^ percentile), and maximum value. The reported % refers to per-protein means, since they reflect the average disorder level of a typical protein within each class.

**Figure 3.**
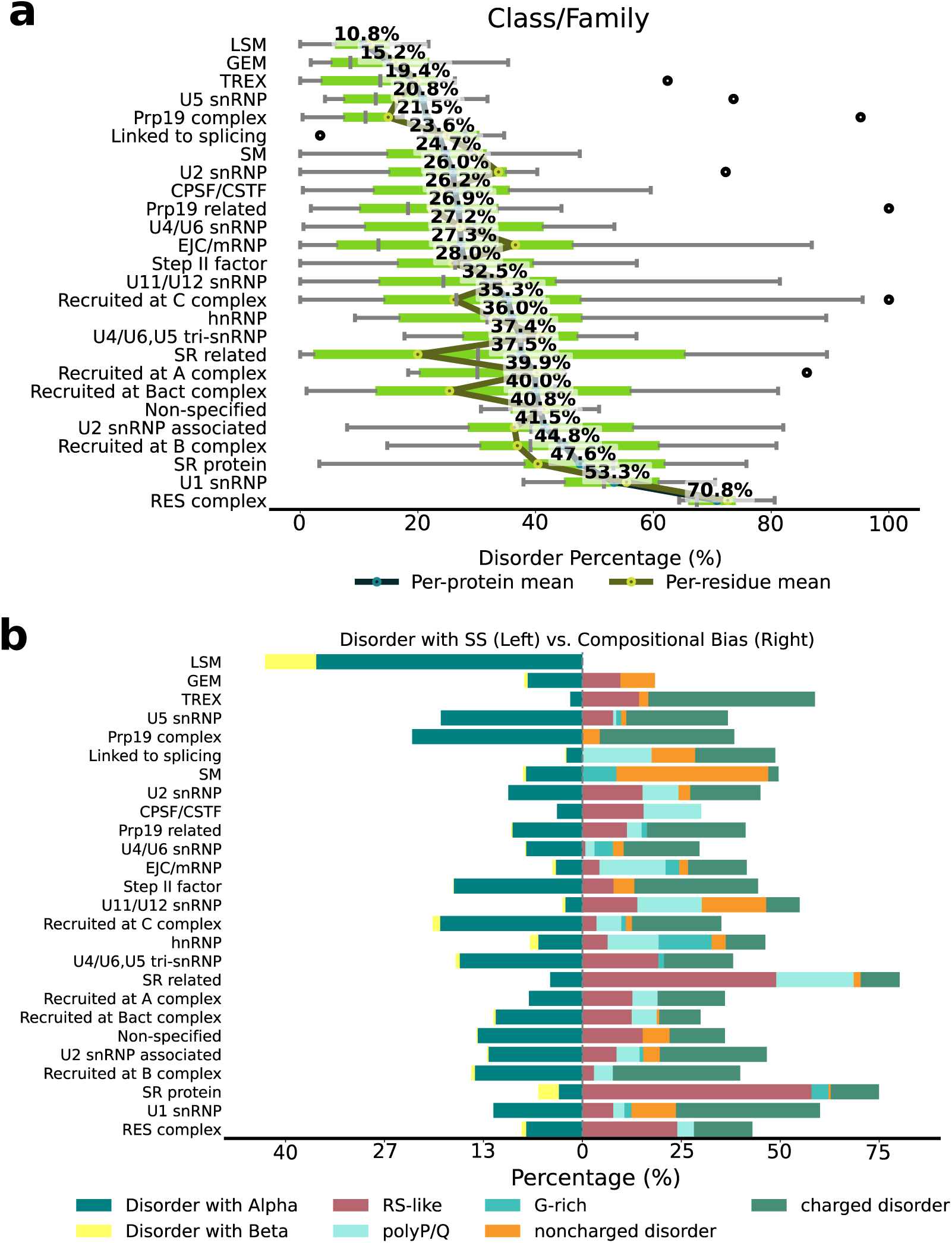
Disorder percentage and type in spliceosome. A) Class/Family of spliceosome proteins. Per-protein and per-residue mean are reported in blue and yellow dots, respectively. The green bars represent boxplots, showing the distribution of disorder percentages with five key statistical measures: minimum value, first quartile (25^th^ percentile), median, third quartile (75^th^ percentile), and maximum value. The reported % refers to per-protein means, reflecting the average disorder level of a typical protein within each class. B) Type of intrinsically disordered content in different protein classes. Disorder content is divided in disorder with secondary structure (SS, left) vs disorder with compositional bias (CB, right). In the disorder with SS the percentage of residues containing α-helix and β-sheet are depicted in yellow and blue, respectively. In the disorder with CB: RS-like, poly P/Q, G-rich, non-charged, charged are depicted in red, light-blue, cyan, orange, dark green, respectively.

We first evaluated the degree of disorder in spliceosome complexes. We observed that in the major spliceosome, the initial E complex exhibits the highest disorder content. The percentage decreases in the A complex until the pre-B complex is reached, slightly increasing in the B^act^ and B* complexes, where the first splicing reaction occurs, to remain roughly constant until the P complex forms. Disorder content decreases then drastically at the ILS complex (Figure 2A). These data imply that in the central and final steps of the splicing cycles, where the spliceosome complex is fully assembled and engaged in catalysis, and in dismantling its components, the IDRs’ plasticity is less crucial.

Although data on the minor spliceosome are more limited, it overall appears to contain a lower proportion of disordered residues, with the highest content being in the minor spliceosome A (AT-AC) complex (31%). Conversely, the disorder content of the remaining minor-spliceosome specific complexes remains constant (27%) and increases during 3’-splice site cleavage (45.3%) (Figure 2B).

We next analyzed the disorder content by protein classes or families. We observed that only three families have less than 20% of disorder content (i.e. LSM (10.8%), GEM (15.2%) and TREX (19.4%)). Conversely, many families exhibited disorder content equal or above 40% (i.e. proteins recruited at B^act^ complex (40.0%), Non-specified, i.e., no classes or families labeled/annotated (40.8%), U2 snRNP associated (present at A, B, B^act^ and B* complexes) (41.5%), Recruited at B complex (44.8%), SR protein (47.6%), U1 snRNP (53.3%) (both present at E, A, B complexes) and RES complex (present at A, B, B^act^ complexes) (70.8%) (Figure 3A). These results confirm that the early spliceosome complexes contain proteins with the largest IDRs. However, the proportion of disordered/ordered sites is different in every complex (Figure 2A).

We also annotated the disorder types, classifying the IDRs as CB-IDR and SS-IDR. CB-IDRs are characterized by the presence of residues or motifs with a higher frequency than normally expected for in vertebrates’ proteins (Figure 3B). This analysis revealed that CB-IDR content varied between 0 and 75%, with the SR and SR-related proteins exhibiting the highest content.

The most common types of CB-IDR were associated with the presence of charged residues (18.57%), followed by RS-like motifs (12.82%) and poly P/Q sequences (6.0%). However, the relative abundance of these CB-IDR segments was different in distinct protein classes (Table S1). Besides the RES, SR and SR-like proteins, in which the RS-like content was the largest, in the other groups the CB-IDRs were owed to the presence of charged residues.

Conversely, SS-IDR (i.e., regions predicted to be disordered and to contain secondary structure content (i.e. α-helices or β-sheets)) varied from 2 to 43%, with LSM (43.7%) being the class with the largest SS-IDR content, followed by Prp19 complex (23.4%), proteins recruited at C complex (20.6%) and U5 snRNP proteins (19.5%). Within the SS-IDR, the α-helical type of secondary structure was predominant (Figures 3B and S1-S2).

Interestingly, the disorder content in LSM, Prp19 complex and U5 snRNP classes of proteins is low. The existence of these SS-IDRs in these regions suggest that they are transitional. In a few cases, IDRs were characterized by a proportional amount of CB-IDR and SS-IDR content (U4/U6.U5 snRNP, Step II factor, recruited at C complex, Prp19 complex), suggesting that these classes may have a promiscuous behavior during splicing.

Next, we inspected the presence and type of PTMs occurring within the IDRs of spliceosome proteins (Supplementary Methods 1.6). Firstly, we calculated the relative abundance of PTMs in ordered or disordered regions of the spliceosome proteins. An enrichment test, comparing the proportion of residues with PTMs in disordered regions vs ordered regions, showed that PTMs in IDRs are more abundant (Fisher test p-value = 1.101e^-115^). Namely, we identified 1,012 sites hosting PTMs over 32,068 sites without PTMs in IDRs as compared to 831 sites hosting PTMs over 78,059 sites without PTMs in ordered regions.

Then, we calculated the PTMs density per protein, revealing that PTMs per residue are significantly more abundant in IDRs (Wilcoxon test p-value=5.782e^-15^). Additionally, we evaluated the PTM distribution per class/family (Figure 4A), confirming the prevalence of PTMs in IDRs (> 50%). Finally, we inspected the relative abundance of each PTM type, defined as the proportion of PTMs located in IDRs relative to the total number of PTMs across the entire protein sequence (including both ordered and disordered regions). Interestingly, citrulline (0.66), phosphoserine (0.64), N-acetylalanine (0.6) and N-acetylserine (0.54) were more abundant in IDRs than in ordered regions (ratio > 0.5) (Figure 4B). Notably, these PTMs are commonly associated to diverse biological functions ranging from rapid signaling (phosphoserine), epigenetic regulation (citrulline) and fundamental protein life-cycle management (N-acetylation) (22–26), suggesting that they play similar regulatory roles in splicing.

**Figure 4.**
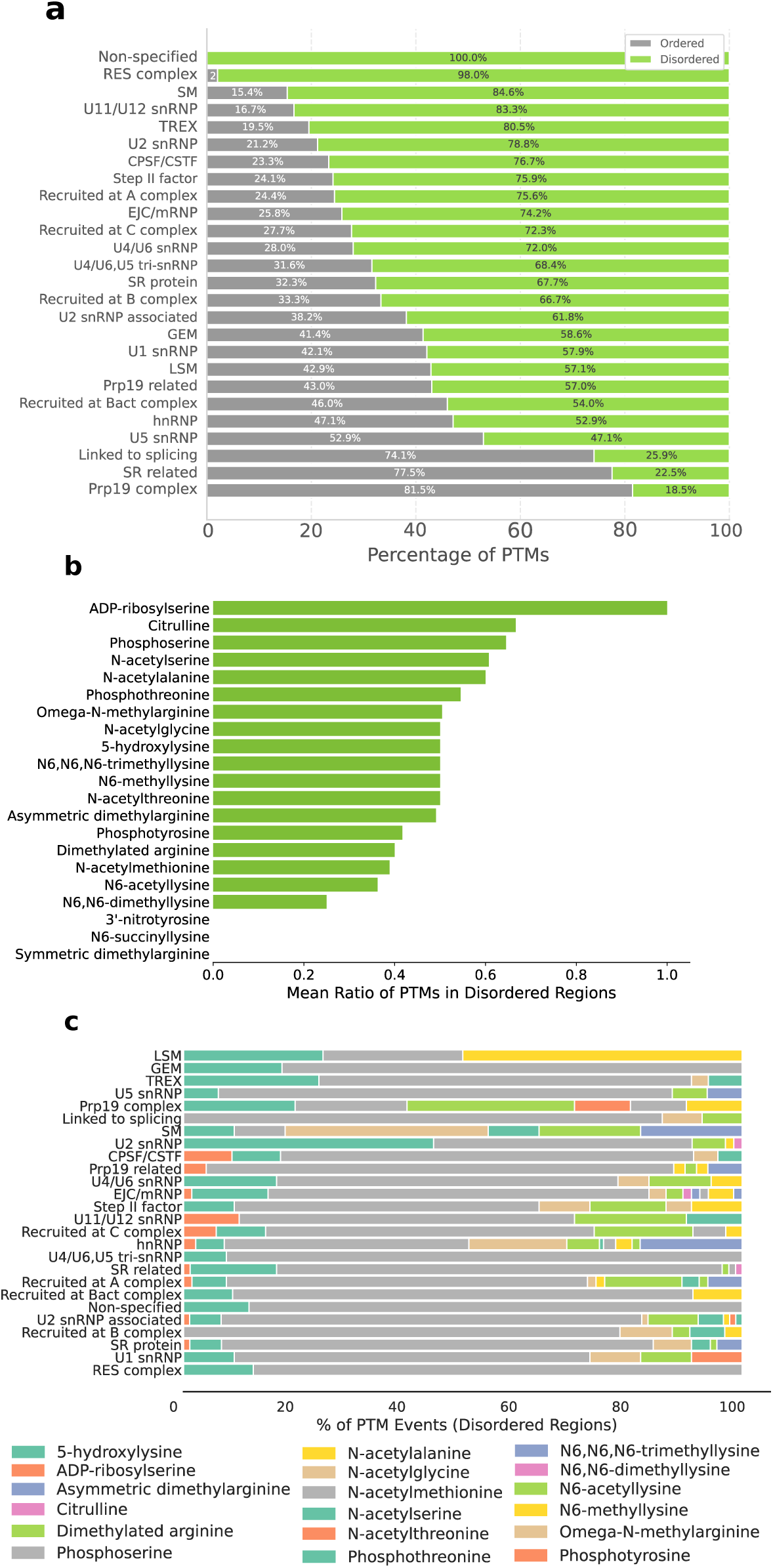
Distribution and characteristics of post-translational modifications (PTMs) in spliceosome proteins. A) PTM distribution in different class/family groups (ordered vs disordered regions). B) Comparison of the proportion of PTMs occurring within intrinsically disordered regions (IDRs) versus those found across the whole sequence of spliceosome proteins. C) Relative abundance of different PTM types across groups/families of spliceosome proteins. Groups are listed according to PTMs abundance from the top to the bottom. PTMs data, presented as percentages, provide insights into PTM preferences and potential functional specialization in subgroups.

**Figure 5.**
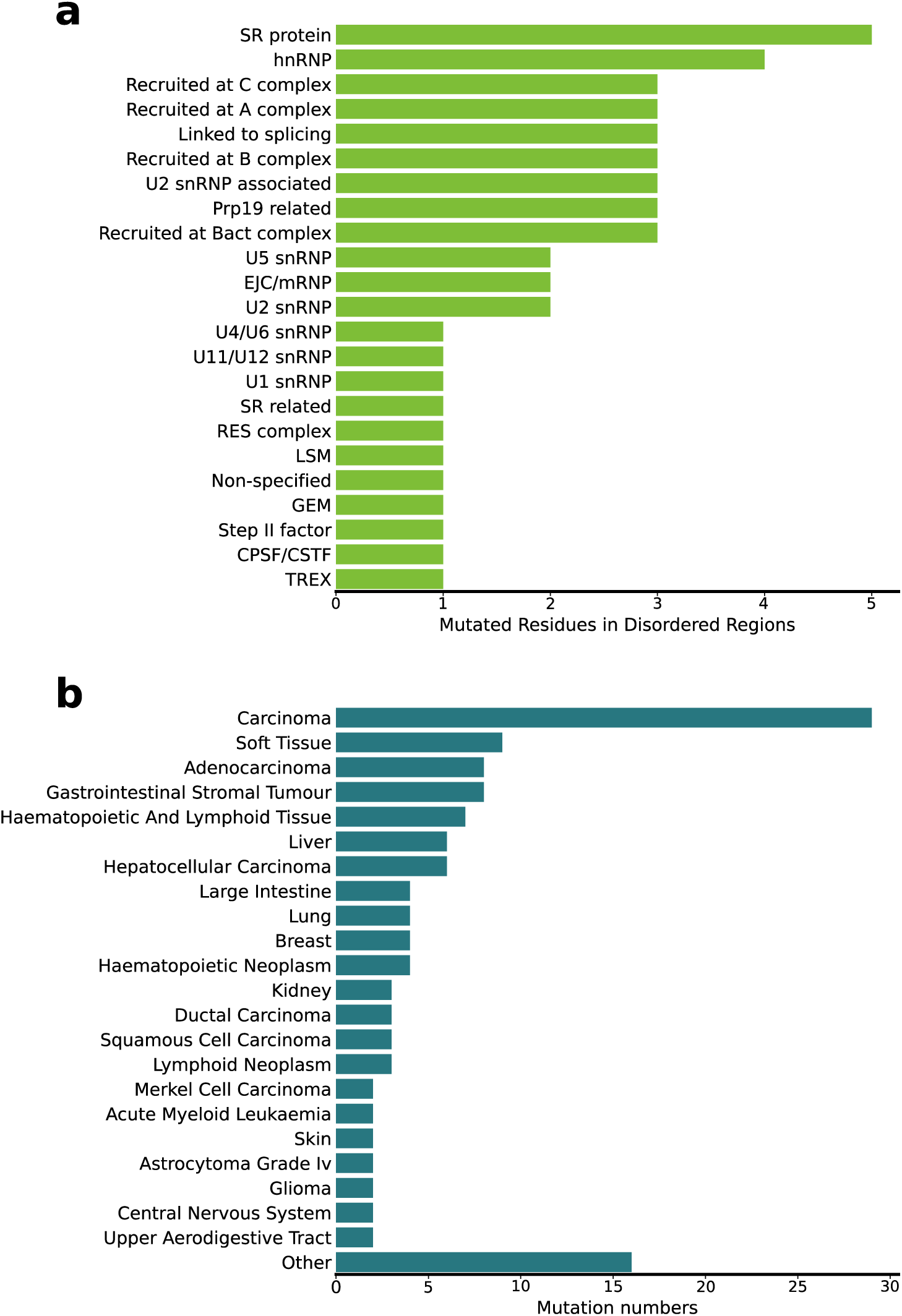
Cancer-associated mutations in intrinsically disordered regions (IDRs) of spliceosome proteins. A) Number of mutated residues within IDRs across different protein families. B) Number of mutations occurring in IDRs associated with distinct tumor types.

Focusing only in IDRs, we then inspected the abundance of PTM types in different protein class/families (Table S2), with phosphoserine (65,7%) and phosphothreonine (11.2%) being the predominant ones (PTM percentage was calculated as the amount of a specific PTM type divided by the sum of the overall number of all PTM types) (Figure 4C). These PTMs may be involved in modulating protein-protein interactions and subcellular compartment regulation. Other PTM types, such as N6-acetyllysine (6.1%), omega-N-methylarginine (4.4%), N-acetylalanine (3.9%) and asymmetric dimethylarginine (2.4%), instead, showed class/family-specific patterns.

Due to the centrality of the spliceosome for gene expression and regulation and its implication in cancer (27–29), we further investigated the frequency of cancer-causing mutations in IDRs (Supplementary Methods 1.6). This analysis revealed that among spliceosome protein families, the SR and hnRNP proteins are most-frequently objects of mutations (Figure 4A). Overall, 43 proteins contained IDRs that host cancer-associated variants. Among them are FUS, RMB10, SF3B1, SRSF2, U2AF1 and THOC2 (Table S3). The number of mutations in the IDRs of these proteins varied from 1 to 5 and were associated with different types of tumors, with carcinomas being the most abundant category (Figure 4B). Remarkably, their implication in diverse cancer types reflects the broad impact of splicing alterations in tumorigenesis. Complete data on mutations in ordered and disordered regions is provided in Tables S3 and S4.

To check for an association between PTMs and cancer-associated variants, we even examined whether cancer-causing mutations, flanking (i.e. located within five residues of (30) ) the PTM site, occurred frequently.

This enrichment analysis identified 17 proteins hosting cancer-causing mutations nearby the PTMs sites. In nine of these proteins (FUS, THOC2, SRSF2, WBP11, U2AF2, SF3B1, RBM10, DDX41, RBM8A) these cancer-associated variants are related to skin abnormality (Human Phenotype Ontology identifier HP:0000951), while in five of them (SRSF2, SF3B1, PRCC, DDX41, RBM8A) to neoplasms (HP:0002664). In both cases, the mutations are predominantly associated with phosphoserines and phosphothreonines (Table S5).

Next, we ranked the proteins by their disorder fraction and considered their cellular abundance. Among the proteins containing the largest IDRs, only a subset was highly abundant (Table 1). These proteins (SRRM1, FUS, YBOX1, and RU17) have disorder content ranging from 50 to 90% and are predominantly characterized by CB-type of disorder, even if the type of residues causing the CB varies (Supplementary Results 1). The observed differences in the type of sequences causing the CB-type of disorder may allow their multivalent interactions critical to form and remodel dynamic ribonucleoprotein complexes (36,37). Their high expression levels and disorder content suggest that these proteins may play critical roles in cellular homeostasis as detailed in Supplementary Results 1. The regulatory role of FUS, SRRM1, and YBX1 is further supported by their hosting of multiple PTMs (Table S2).

**Table 1.**
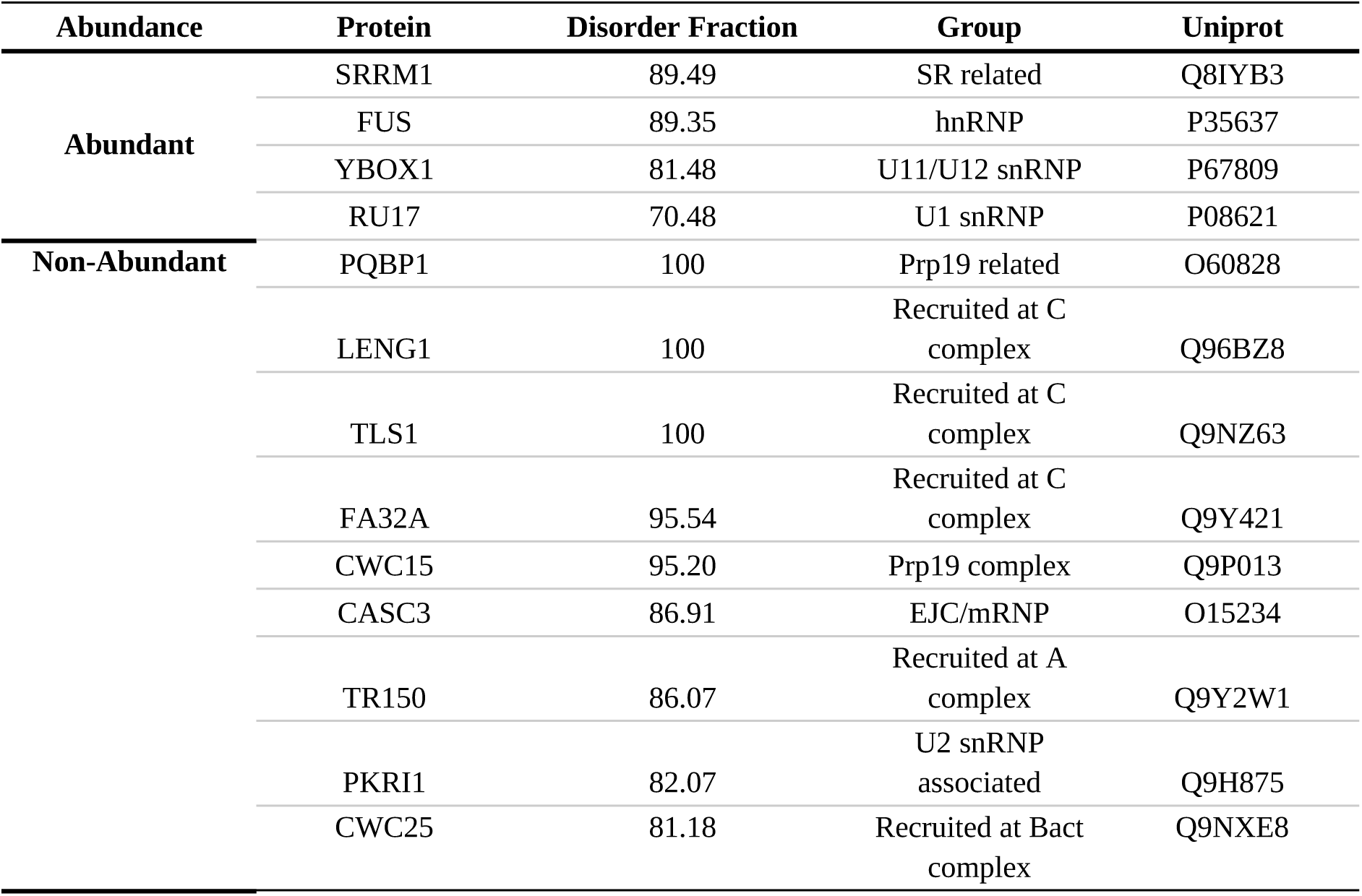
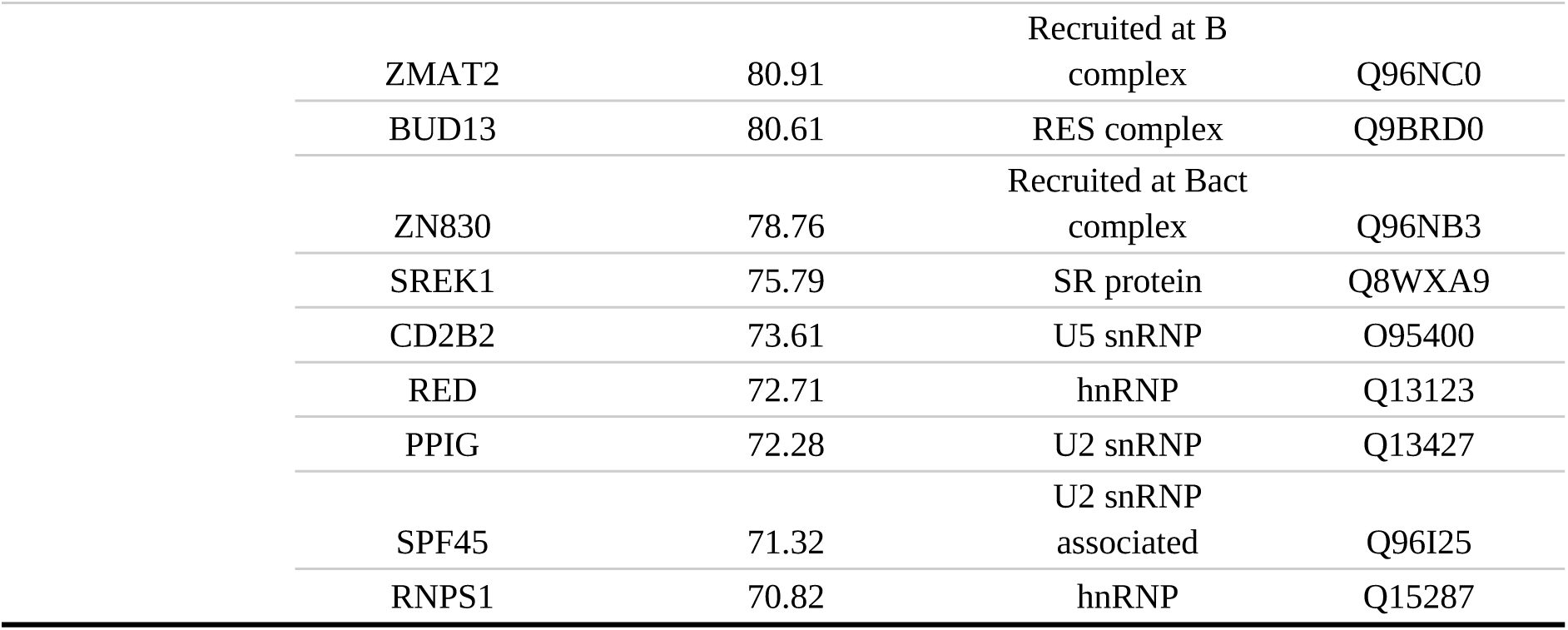
Spliceosome proteins exhibiting the highest disorder fraction (above 70%)

Additionally, we investigated the evolutionary dynamics of spliceosome proteins exhibiting the highest IDR content (> 70%) (Supplementary Methods 1.5, Supplementary Results S2, Figures S4-S25). Specifically, we inspected whether structural disorder played a role in the evolutionary history of these proteins and, if their orthologs retained similar disorder content. To this end, we constructed phylogenies for 19 proteins. Multiple sequence alignments revealed that, in most cases, the high disorder content was maintained across orthologs (Supplementary Results 2). Only, in a few cases, specific clades deviated markedly, displaying loss in disorder, thus reflecting different evolutionary trajectories (Supplementary Results S2).

Commonly, disordered sites of proteins display rapid evolutionary dynamics, while ordered sites tend to be more conserved. This metric is evaluated through disorder-to-order transitions (DOT), which refers to the change between ordered and disordered states. Interestingly, when analyzing this property in spliceosome IDRs, we observed that SS-IDRs, located adjacent to ordered regions in the same protein or at positions corresponding to ordered regions in ortholog proteins, exhibited significantly lower DOT rates as compared to the other disordered regions (Mann-Whitney U = 2.39 x 10^6^, p < 10^-9^). Due to their lower DOT values and preservation of amino acid sequence, SS-IDRs appear to be more evolutionarily conserved than the other random disordered sites, suggesting that their secondary structure imposes evolutionary constraints across orthologs.

Notably, when analyzing the increase/decrease of disorder content across evolution (net gain of disorder), clades exhibited mixed trends, indicating protein specialization and different functions for the IDR content in different organisms (Figures S24-S25). Collectively, this analysis revealed that the predominant evolutionary trend in IDRs was based on persistence of SS-IDRs. These regions are thus likely conserved to play a functional role in splicing regulation.

## Discussion

Half of the spliceosome proteins were predicted to contain extensive IDRs (48.5%), which are unevenly distributed in amount and type. Notably, a big proportion (35.7%) of these proteins had >40% of their sequence classified as disordered. Disorder content showed a strong correlation with RS-like and charged compositional bias and approximately 15% of disordered regions alternates with secondary structure formation. The highest percentage of disorder was observed at the E complex, suggesting that IDRs are key for early spliceosome assembly.

Notably, spliceosome IDRs are target for PTMs (31), with phosphorylation of serine and threonine residues being the most abundant type. The addition of a bulky and negatively charged phosphoryl group is expected to regulate splicing by modulating interaction affinity or complex assembly, particularly in SR proteins and other splicing regulators (32). This regulatory principle is exemplified in several core spliceosome components. As an example, phosphorylation of SAP155 (U2 snRNP) is tightly coordinated with catalytic steps (33,34), phosphorylation of PRP28 by SRPK2 is essential for stable tri-snRNP integration (35,36), and multi-site phosphorylation of SF3B1 N-terminal by CDK11 modulates RNA interaction within the B^act^ complex (37).

Interestingly, only a few among the most disordered proteins (e.g., SRRM1, FUS, YBOX-1, SNRNP70) are abundant in the human proteome. Their function may be that of interaction hubs within the spliceosome and in dynamic ribonucleoprotein aggregates. The remaining highly disordered proteins, characterized by lower cellular abundance, must be instead more specifically implicated in splicing regulation.

Across evolution, spliceosome proteins retain the SS-IDR content (Figure S24-S25), which is conserved in different clades. In contrast to other protein families whose IDRs are rapidly evolving (38), spliceosome IDRs display low rates of disorder-order transitions throughout evolution, as reflected by their low changes per node values (ranging from 0.01 to 0.08). Notably, higher changes per node are concentrated in SS-IDR segments, indicating that the limited DOT are focused on specific residue positions across lineages. This pattern suggests constrained flexibility, which is consistent with the function of spliceosome proteins as an essential cellular machinery evolving under purifying selection (39), and may be associated with the presence of phosphorylation sites (40).

Lastly, we identified multiple cancer-associated mutations in IDRs which cluster in splicing-related families, such as SR proteins, hnRNP, and mostly in proteins recruited at A, B, B^act^ and C complexes. Since spliceosome IDRs flexibly interlace proteins and RNA, even subtle perturbation in these interaction networks may alter their function ultimately triggering splicing defects. The tumor types associated with these variants were diverse, confirming the systemic impact of splicing dysregulation in cancer.

Overall, our study underscores the central role of IDRs in splicing regulation and disease, revealing that spliceosome IDRs are abundant, evolutionarily conserved, and functionally important regions that host regulatory and cancer-associated variants. These features align with the highly dynamic, transient protein–protein and protein–RNA interactions driving spliceosome function.

## ASSOCIATED CONTENT

### Supporting Information

Supplementary Information consists of Supplementary Methods, Supplementary Tables S1-S3, and Supplementary Figures S1-S25.

## Author Contributions

The manuscript was written through contributions of all authors. All authors have given approval to the final version of the manuscript.

## ACKNOWLEDGEMENT

We thank the MUR-PRIN2022 project CUP B53D23025350001 for financial support. A.M. acknowledges the CINECA award under the ISCRA initiative, for the availability of high performance computing resources and support. A.M. thanks the Italian Association for Cancer Research (project AIRC IG 24514).

## Data availability

Data, scripts, input will be available at https://github.com/bposantos/spl_idp_project.git. following the FAIR principles [https://doi.org/10.1038/s41592-025-02635-0].

