## Supplementary Material for "Exploring Disordered Regions of Human Spliceosome Proteins"

#### Table of Content

Supplementary Methods ……………………………………………….pages 2-4

Supplementary Results ………………………………………………..pages 5-9

Supplementary References ……………………………………………pages 10-12

Supplementary Figures S1-S25 ………………………………………pages 14-32

Supplementary Tables S1-S5 …………………………………………pages 33-67

### Supplementary Methods

#### 1.1 Sequences Retrieval

The pipeline of work is represented in Figure 1. Protein sequences were retrieved from the UniProt database using the following query: (go:0005681) AND (organism_id:9606), which selects proteins annotated with the Gene Ontology (GO) term for the spliceosome (GO:0005681) in *Homo sapiens* (organism ID: 9606). The search, performed on December 3rd, 2024, returned 383 protein entries. Sequences were subsequently curated by removing duplicates (equal sequences), fragments (sequences present in bigger sequences), and alternative isoforms, resulting in a final dataset of 205 unique full-length proteins. Sequences were manipulated in FASTA format and all data treatment was performed using *in house* Python scripts. Data and scripts will be available at https://github.com/bposantos/spl_idp_project.git.

#### 1.2 Disorder Prediction

Intrinsic disorder in protein sequences was predicted using tools available through the CAID (Critical Assessment of Intrinsic Disorder) portal. Disorder annotations were generated using a subset of predictors evaluated in the CAID2 challenge (1), which benchmarked disorder prediction methods across multiple datasets. We selected the top five performing predictors based on ROC curve performance under the Disorder-NOX benchmark (selected to avoid bias from missing experimental annotations): fIDPlr2, fIDPnn2, DisPredict3, IDP-Fusion, and DisoPred. Binary consensus disorder predictions were obtained via majority voting across the five selected CAID2 predictors, i.e., each predictor gives a score, converted to 0 (ordered) or 1 (disordered), and if residue i resulted in 1 per 3 or more predictors, the consensus sequence received i = 1, otherwise, i = 0 (2). The access date to perform the analysis ranged from February 05 to May 20 2025. Modified residues were discarded, since the portal did not support any “X” residue. The output files were stored as .tsv files, and the consensus sequences were transferred to a single .csv file for further manipulation.

#### 1.3 Computational Workflow and Python Scripts

All data processing and analysis steps were implemented in Python (v3.8) using standard scientific libraries such as pandas, Biopython, NumPy, and Matplotlib. Custom scripts were used to: (i) filter UniProt sequences (removal of fragments, duplicates, isoforms); (ii) parse disorder prediction outputs and create correspondence among sequence and disorder positions; (iii) detect and annotate compositional bias regions using fLPS algorithm; (iv) retrieve PTMs with UniProt API to disordered regions; (v) create correspondence among the spliceosome IDR database and the COSMIC database in order to extract the information about mutation in disordered regions, (vi) generate figures and compute statistics.

#### 1.4 Sequence Analysis

To extract functional and structural information from IDRs, we developed custom Python scripts to integrate multiple layers of annotation. Specifically, we retrieved and combined data on: (i) disorder and secondary structure predictions, (ii) disorder and compositional bias regions, and (iii) disorder with post-translational modifications (PTMs). Secondary structure information was obtained using the S4Pred predictor, which enables the analysis of structural propensities within disordered segments (3).

Consistently with the annotation of Korneta and Bujnicki (2012) (4) for the compositional bias (CB-IDRs), we searched for hnRNP-like G-rich segments, which are regions containing RGG and related repeats ([RSY]GG, R[AGT][AGTFIVR]). For the ‘non-charged’/‘charged’ disorder IDR, we searched for regions rich in non-charged/charged residues (PQMGVWA)/(RKDE). For RS-like segments, we searched for IDRs that are rich in arginine and serine residues; and for polyP/Q segments, IDRs that contain repeats of proline or glutamine residues

We classified a type of disorder as disorder with secondary structure (SS-IDR) whose sequences are predicted to be coiled-coil with secondary structure. CB-IDRs were identified using the fast Lowest Probability Subsequences (fLPS) algorithm, which detects low-probability amino acid subsequences within protein sequences (5). The method calculates the log-probability of sliding windows based on a background frequency model. In this case, we considered amino acid frequencies specific to vertebrates. Regions where specific amino acids are significantly overrepresented were assigned statistical scores. Using dynamic programming, fLPS efficiently merges overlapping or adjacent biased segments. The output includes the coordinates, type of compositional bias, and the statistical significance of each identified region.

##### 1.5 Post translational Modifications (PTMs) and Cancer associated mutations

Post translational modifications (PTMs) were retrieved from the UniProt API. To obtain information also on the presence of cancer associated mutations in these regions, the COSMIC database was used. Cosmic_CompleteTargetedScreensMutant_v102_GRCh38.tsv and Cosmic_Classification_v102_GRCh38.tsv were downloaded from their website to obtain: gene symbol, COSMIC gene ID, COSMIC phenotype ID, mutation amino acids, tumor primary site, and subtypes (6). Protein abundance was obtained from the human proteome from PaxDb (7). Spliceosome proteins were retrieved, and the top 25% most abundant proteins were classified as abundant, while the remaining ones as non-abundant.

##### 1.6 Phylogenetic analyses

Human reference protein orthologs were identified using the InParanoid database (8). Sequence and species duplications were removed. Uniprot API was used for sequence retrieval of the orthologs sequences, and they were employed for Intrinsically disordered proteins (IDP) prediction using CAID IDP portal. Multiple sequence alignment (MSA) was performed using MAFFT with the flags --localpair and maximum iteration of 1000 (9). To remove excessive gaps in the MSA and create accurate analysis in next steps, a trimming was executed using TrimAL with the flags -gt 0.3 and - cons 50 (10). Phylogenetic trees and ancestral sequence reconstruction were obtained using IQ-TREE (11) and plotted using ETE3 python library (12). Analyses were performed, initially, for the 22 SPL proteins with more than 70% content of disorder, but three were discarded for having less than 20 orthologs to build the trees: FUS, SREK1 and RNPS1. To root each tree, the most basal leaf was chosen based on the specie's age, using TimeTree webserver for the selection (13). For each protein phylogeny and its corresponding MSA site, state changes (order vs. disorder) were inferred using parsimony as implemented in GLOOME (14) and then normalized by the number of nodes in the phylogenetic tree. When a species contained a gap at a given position, it was excluded from the analysis of that site.

The statistical analyses were performed in several steps to rigorously compare the distributions of net gain values among clades. First, a global Kruskal-Wallis test was applied to assess whether there were significant differences among the clades (A, B, C and D, when present), as this non-parametric test does not assume normality. If the global test was significant, pairwise comparisons between clades were conducted using the Mann-Whitney U test, and the resulting p-values were adjusted for multiple testing using the Bonferroni correction to control the family-wise error rate. To evaluate the homogeneity of variances among clades, both Levene's test (robust to non-normality) and Bartlett's test (sensitive to deviations from normality) were performed. The coefficient of variation was also calculated for each clade to quantify relative variability. Additional analyses included comparing the distribution of the ordered clade to the combined distribution of the other clades using the Mann-Whitney U test, assessing differences in variance, measuring the mean absolute distance from zero (as a proxy for order/disorder), and calculating the proportion of values near zero within each clade. Lastly, the relationship between sequence substitution rate and disorder-to-order transitions was measured by Spearman coefficients. All analyses were performed using Python libraries such as SciPy and statsmodels.

#### Supplementary Results

**1. Abundance of highly disordered spliceosome proteins.**

Proteins characterized by a high cellular abundance and disorder content comprise SRRM1. This protein has an extremely high disorder content (89.5%), with a small fraction of SS content (5.6% of disorder with alpha-helix) and a very high CB type of disorder, which can be divided into: RS-like (37.2%), poly P/Q (22.0%), noncharged (2.0%) and charged (11.1 %).

SRRM1 acts as a key scaffold protein, aiding constitutive and exonic splicing enhancer (ESE)-dependent splicing by bridging sequence-specific SR proteins (e.g., SFRS4, SFRS5, TRA2B/SFRS10) and core snRNP components (e.g., SNRP70, SNRPA1). SRRM1 also stimulates mRNA 3′-end cleavage and binds both pre-mRNA and spliced mRNA. SRRM1 also interacts with RNA and DNA with low sequence specificity, showing similar affinity for single- and double-stranded substrates (15–17). Therefore, its high IDR content is consistent with its functional versatility and tendency to interact with multiple partners.

FUS also contains an extremely high (89.35%) disorder content, with a small fraction of secondary structure (α-helix 0.21% and β-sheet 3.83%), and a larger fraction CB where the G-rich regions are the most abundant (31.94%) followed by poly P/Q (23.38%) and RS-like (2.09%). FUS binds single-stranded RNA containing the consensus sequence 5′-AGGUAA-3′ (18) and associates with nascent pre-mRNAs to couple transcription and splicing by mediating interactions between RNA polymerase II and U1 snRNP (19). FUS also autoregulates its expression via binding to its own pre-mRNA and promoting nonsense-mediated decay (20).

YBOX1, containing 82.48% of disorder, has exclusively CB disorder. This contains non-charged residue type (25.62%) followed by RS-like (18.52%) and poly-P/Q (8.33%). YBOX1 primarily functions as an RNA-binding protein, recognizing the 5′-[CU]CUGCG-3′ motif and selectively binding mRNA transcripts modified with 5-methylcytosine ($m^5C$) (21). It stabilizes target mRNAs by recruiting ELAVL1 (HuR), a key regulator of mRNA stability, thereby preventing degradation (22,23).

RU17 (also known as U1 small nuclear ribonucleoprotein 70 kDa, or U1-70K) is a core component of U1 snRNP, essential for recognizing the 5′ splice site of pre-mRNA and to initiate spliceosome assembly (24,25). RU17 contains 70.48% of disorder, with a small fraction of SS content (5.84% a-helix) and larger fraction of CB disorder die to charged (28.6%), poly P/Q, (4.12%) and of G-rich (2.52%) sequences.

SRSF3 is a serine/arginine-rich splicing factor that binds the consensus motif 5′-C[ACU][AU]C[ACU][AC]C-3′ within pre-mRNA and promotes the inclusion of specific exons during alternative splicing (26,27). Its disorder content is 50%, contains 8.54% of a-helical and 40.24% of b-type disorder with SS, and but predominantly an 87.8% of RS-like CB disorder. As such, the high degree of intrinsic disorder in these proteins is predominantly of CB type, even if the types of abundant residues change in the different cases. The differences in the type of CB sequences enables flexible and multivalent, interactions critical for forming dynamic ribonucleoprotein complexes and for facilitating phase separation, particularly in the cases of FUS and SRRM1 (15,28).

**2. Disorder as an evolutionary strategy for conformational flexibility**

Considering the 22 SPL proteins with more than 70% of disorder content, we investigated whether structural disorder plays a role in the evolutionary history of these proteins (22 minus 3, which were not able to have their trees built) and, consecutively, if their orthologs retained similar levels of disorder (Figure S5-S23). The multiple sequence alignments reveal that, in most cases, proteins follow the query protein in the high disorder content. However, specific clades can deviate markedly, displaying lower or altered disorder and thus, reflect different evolutionary trajectories.

Leukocyte receptor cluster member 1 (Leng1) orthologs had a common ancestor with intermediate levels of disorder in the sequence region 257-281. Clade A (Phylum: Tracheophyta; Kingdom: Plantae) diverged and specialized, with a loss of disorder in this region, whereas the other clades continued the ancestral pattern. Clade B (Phylum: Ascomycota, Basidiomycota; Kingdom: Fungi) maintained similar levels of disorder to the ancestral, while clade C (Phylum: Chordata, Nematoda, Arthropoda; Kingdom: Animalia) slightly gained in levels of disorder (Figure S24A). Our results confirm that Animalia proteins contain much more disorder content than other kingdoms, consistent with a strategy of conformational plasticity. In contrast, Plantae emerge as the most ordered lineage, with over 95% of residues near zero and minimal variability, reflecting an emphasis on structural precision. Fungi occupy an intermediate position but align statistically with Animalia, forming a shared disordered strategy that suggests ecological convergence and possibly close evolutionary kinship. Statistical comparisons confirm that Plantae differ radically from both Fungi and Animalia ($p < 10^{-13}$), whereas Fungi and Animalia do not differ significantly from each other ($p = 0.12$).

Pqbp1, a polyglutamine-binding protein 1, orthologs belong to the Animalia kingdom. A loss in disorder in clade A occurred from residues 100-126 (Phylum: Arthropoda, Nematoda), with some branches with high levels of order. Clade B and C are both from the phylum Chordata, but clade B (Class: Actinoperi) has the same levels of disorder from the ancestral proteins, representing an equilibrium state, but with a gain in disorder in clade C (Classes: Mammalia, Aves, Squamata, Crocodylia, Testudines) (Figure S24B). Our findings reveal an evolutionary gradient in protein structural strategies within the animal kingdom. Invertebrates such as Arthropoda and Nematoda (A) exhibit a markedly ordered profile, reflecting a conservative strategy. Vertebrates, by contrast, diverge toward disorder: fishes (B) already show significant conformational plasticity, representing an intermediate stage that supports aquatic adaptability, while tetrapods (C) display only disorder in the region, with no residues near neutrality and the highest variability, consistent with a radical strategy of maximum flexibility. Statistical tests confirm that invertebrates differ significantly from vertebrates ($p < 10^{-13}$), whereas fishes and tetrapods cluster together without significant differences ($p = 0.12$).

Tr150, a Thyroid hormone receptor-associated protein 3, orthologs, from sites 491-565, have a singular evolutive story. All branches belong to the phylum Chordata. Clade A (Class: Mammalia, Amphibia, Squamata, Aves), B and C (Class: Actinopteri), with B belonging to non-percomorph and C to percomorphs (Figure S24C). Tetrapods developed a unique evolution strategy for this region (Tetrapods vs fishes: $p = 9.77e^{-09}$). In fishes, the percomorphs radiation seems to have changed the disorder strategy: percomorphs lost in net gain disorder ($p = 2.11e^{-14}$). Clade B is high in disorder while clade C is an ordered region, representing an innovative strategy for this portion of the protein. The results highlight a distinct evolutionary strategy in Clade C (derived percomorph fishes) compared to tetrapods (Clade A) and basal actinopterygians (Clade B). While A and B are statistically indistinguishable and share highly variable, broadly disordered profiles (CV up to 112%, low near-zero residue proportion), Clade C stands apart with a significantly different pattern (pearson correlation $< 10^{-13}$), showing reduced variance, a three-fold higher proportion of near-neutral residues (62.7%), and overall structural order. This suggests that percomorphs did not simply inherit the disorder tendencies of vertebrates but instead evolved a distinct regime of constraint, possibly linked to their extensive ecological radiation and functional diversification. In this view, the percomorph lineage represents not a reversion toward order but the adoption of a novel adaptive strategy, distinguishing it sharply from both tetrapods and more basal fish clades.

Evolutionary analysis of Bud13 homolog protein reveals distinct trajectories of structural disorder across major vertebrate lineages. In Actinopteri (Clade A, fishes), the protein shows a substantial gain in disorder (+0.355), suggesting an evolutionary shift toward increased flexibility. By contrast, tetrapods (Clade B: Mammalia, Amphibia, Squamata, Aves) remain close to neutral (−0.048), while the mammalian subclade (Clade C) exhibits a pronounced loss in disorder (−0.164). These patterns are statistically robust, with highly significant differences among groups ($p = 6.52 \times 10^{-32}$) and a clear divergence of mammals from other tetrapods ($p = 5.38 \times 10^{-4}$). Taken together, the results indicate a progressive reduction in disorder content along the mammalian lineage, consistent with an evolutionary trend toward increased protein folding and structural constraint. Moreover, the rate of change is accelerated in mammals, with a 3.84-fold greater shift relative to other tetrapods ($p = 1.19 \times 10^{-4}$), pointing to a lineage-specific tightening of protein organization that may reflect functional adaptation of Bud13 homolog in mammalian systems (Figure S24D).

Zn830, Zinc finger protein 830, clade A (Phylum: Ascomycota, Basidiomycota, Mucoromycota; Kingdom: Fungi); clade B (Phylum: Tracheophyta; Kingdom: Plantae); clade C (Phylum: Chordata; Kingdom: Animalia); clade D (Phylum: Arthropoda, Nematoda and Brachiopoda; Kingdom: Animalia). Data in Figure S24E shows Chordate went through a molecular revolution in direction to order, while other animals are similar to fungi. The difference between Chordate and fungi is great ($p = 6.15e^{-10}$), Chordate and plants ($p = 1.33e^{-03}$), and Chordate and other animals ($p = 2.64e^{-08}$). This way, fungi present themselves with a conservative evolutive strategy, with low variance (0.001) and has this region moderately disordered. Plants have a higher variability and multiple evolutive strategies. Chordates are radically different, with higher folding rates, and low variability among the components. Other animals, however, are impressively similar to fungi, not the Chordate.

CD2B2, a CD2 antigen cytoplasmic tail-binding protein 2, orthologs belong to Clade A (Chordata), B (Plantae), C and D (Fungi). Our analyses reveal a clear evolutionary gradient in protein structural strategies across major clades: Chordata (A) stand out as radically more disordered, with high variability and strong departure from neutrality, reflecting an adaptive investment in conformational plasticity. In contrast, Plantae (B) retain an intermediate strategy consistent with the presumed ancestral state of the Animal–Plant lineage, showing remarkable stability and low variance, likely constrained by photosynthetic efficiency, developmental rigidity, and environmental stability. Fungi (C, D) exhibit a progressive internal gradient toward increasing order, with C occupying an intermediate position and D significantly more structured, suggesting ecological specialization and structural constraints as drivers of this trend (Figure S24F). Statistically, Chordata form a distinct group compared to all others, while Plants cluster with both fungal clades, and a significant difference emerges between fungal subgroups C and D. These findings support a scenario in which the common ancestor of Animals and Plants had an intermediate strategy, Plants conserved this ancestral state, Animals diverged radically toward disorder, and Fungi followed a path of progressive ordering, creating a stepwise evolutionary gradient from disordered to ordered strategies.

Altogether, these examples highlight four proteins that display a progressive increase in disorder content culminating in mammals, contrasted with two proteins that follow the opposite evolutionary strategy, showing increased order and greater regional specialization (Figure S25). Considering the 19 proteins analyzed, the predominant evolutionary trend in intrinsically disordered regions is due the persistence of disorder with secondary structure, a feature consistently abundant across all proteins examined. Such disorder with secondary structure likely enhances protein flexibility and functional adaptability.

##### 3. Occurrence of Cancer-Associated Mutations

Cancer associated mutations were frequent in abundant proteins such as in SRSF3, which plays a complex and multifaceted role in cancer biology by functioning as both a proto-oncogene and, in specific contexts, as a tumor suppressor (29–32).

SRRM1 is a splicing factor that significantly contributes to tumorigenesis in both hepatocellular carcinoma (HCC) and prostate cancer. In HCC, SRRM1 promotes malignant behaviors such as increased cell proliferation, migration, invasion, and resistance to apoptosis by activating the JAK/STAT signaling pathway. This correlates with poor patient survival (33). In prostate cancer, SRRM1 is elevated in aggressive and therapy-resistant forms, particularly in castration-resistant prostate cancer. Silencing SRRM1 reduces tumor growth, suggesting its value as both a diagnostic/prognostic biomarker and a therapeutic target in advanced cancers (34).

The FUS protein plays a critical role in oncogenesis. In gastric cancer, FUS expression is significantly elevated and collaborates with the RNA methyltransferase METTL3 to facilitate RNA maturation, a process essential for rapid proliferation of cancer cells (35). This partnership influences critical steps in RNA processing, including the alternative splicing and m6A methylation of mRNAs encoding proliferation-associated genes such as PCNA, MCM2, and the anti-apoptotic factor BIRC5. Hence, also FUS appears as a promising therapeutic target (36).

The YBOX1 has again multifaceted roles in cancer (37) since, by acting as transcription factor and RNA-binding protein, it orchestrates the expression of genes involved in cell cycle progression. YBOX1 directly binds to and activates the promoters of critical regulators, like GSK3B, EGFR, and Cyclin D1, enhancing the expression of cyclins (D1, E1, A, B1) and facilitating the G1 to S phase transition in different cancers (pancreatic, Breast, and Lung adenocarcinomas). Furthermore, YBOX1 collaborates with several long noncoding RNAs (e.g., LINC00665, SBF2-AS1, DARS1-AS1) to activate pro-tumorigenic signaling pathways such as PI3K/AKT, Wnt/β-catenin, and mTOR. Beyond transcription, YBOX1 exerts extensive post-transcriptional control, influencing alternative splicing, mRNA stability, and translation (38). Its activity is entwined with post-translational modifications, particularly phosphorylation at Ser102, which drives its nuclear translocation and enhances its oncogenic functions. Overall, YBOX1 integrates signals from multiple pathways to drive uncontrolled growth, inhibit apoptosis, and promote aggressive cancer phenotypes, making it a promising therapeutic target.

### Supplementary Figures

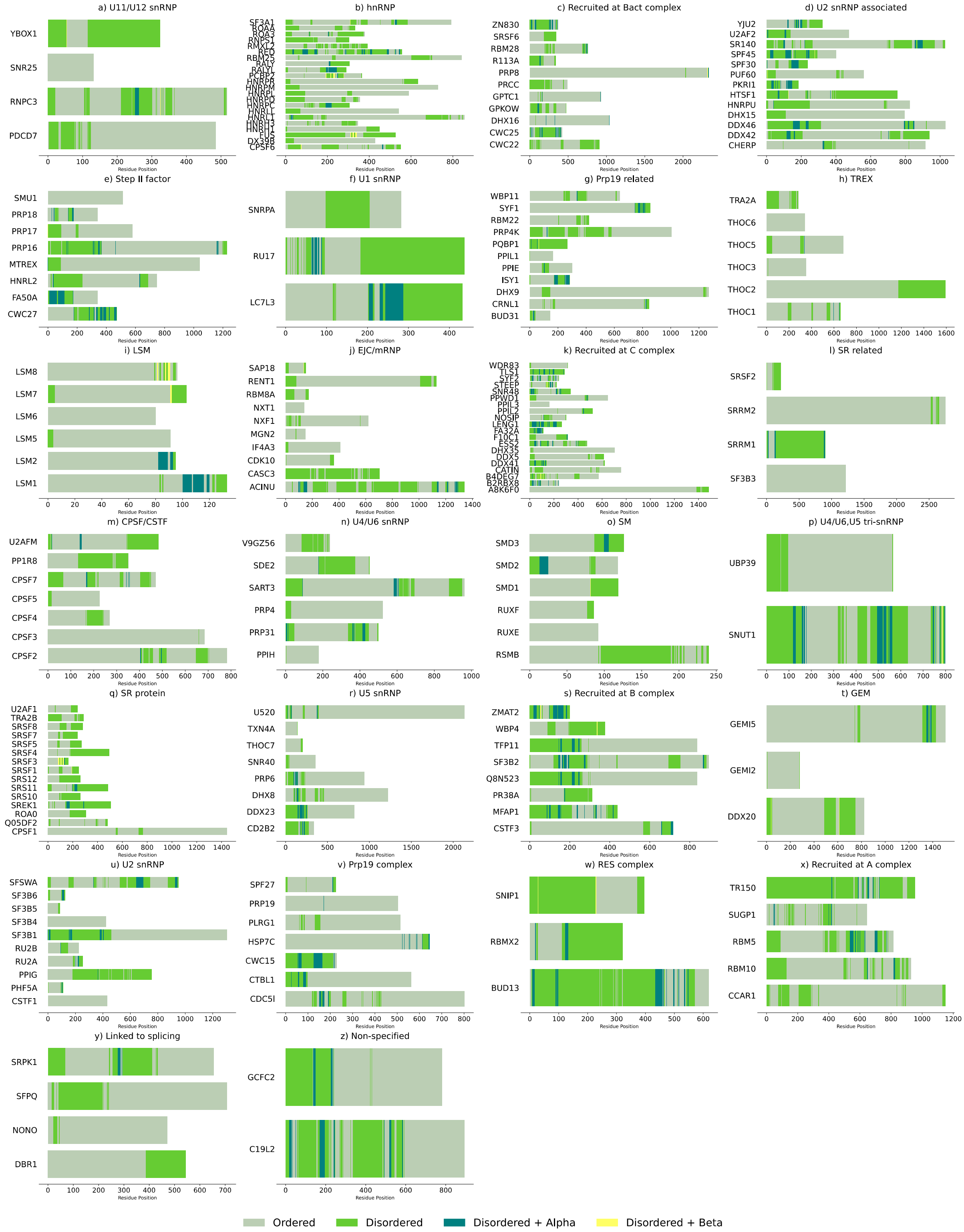


**Figure S1. Segmentation plot of disorder with secondary structure.** A-Z) Class/Family of spliceosome proteins. Each bar represents a different protein. Ordered and disordered residues are shown in grey and green, respectively. Within the green region green-blue and yellow segments refer to alpha and beta-type secondary structure.

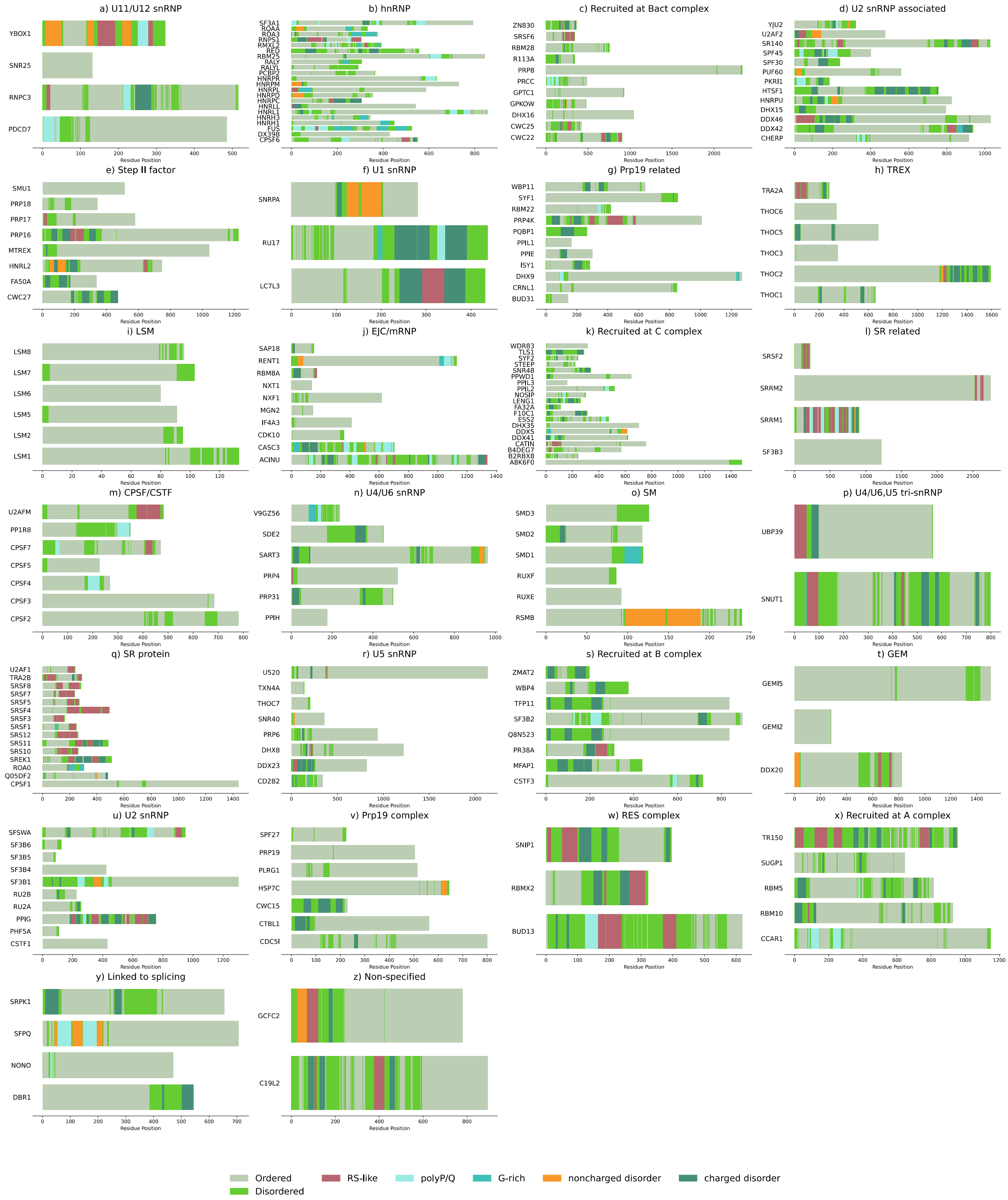


**Figure S2. Segmentation plot of disorder with compositional bias.** A-Z) Class/Family of spliceosome proteins. Each bar represents a different protein. Ordered residues are represented in grey and disordered residues are represented in green. Superimposed is the information about compositional bias: RS-like (dark red), poly P/Q (light blue), G-rich (cyan), non-charged (orange), charged (dark green).

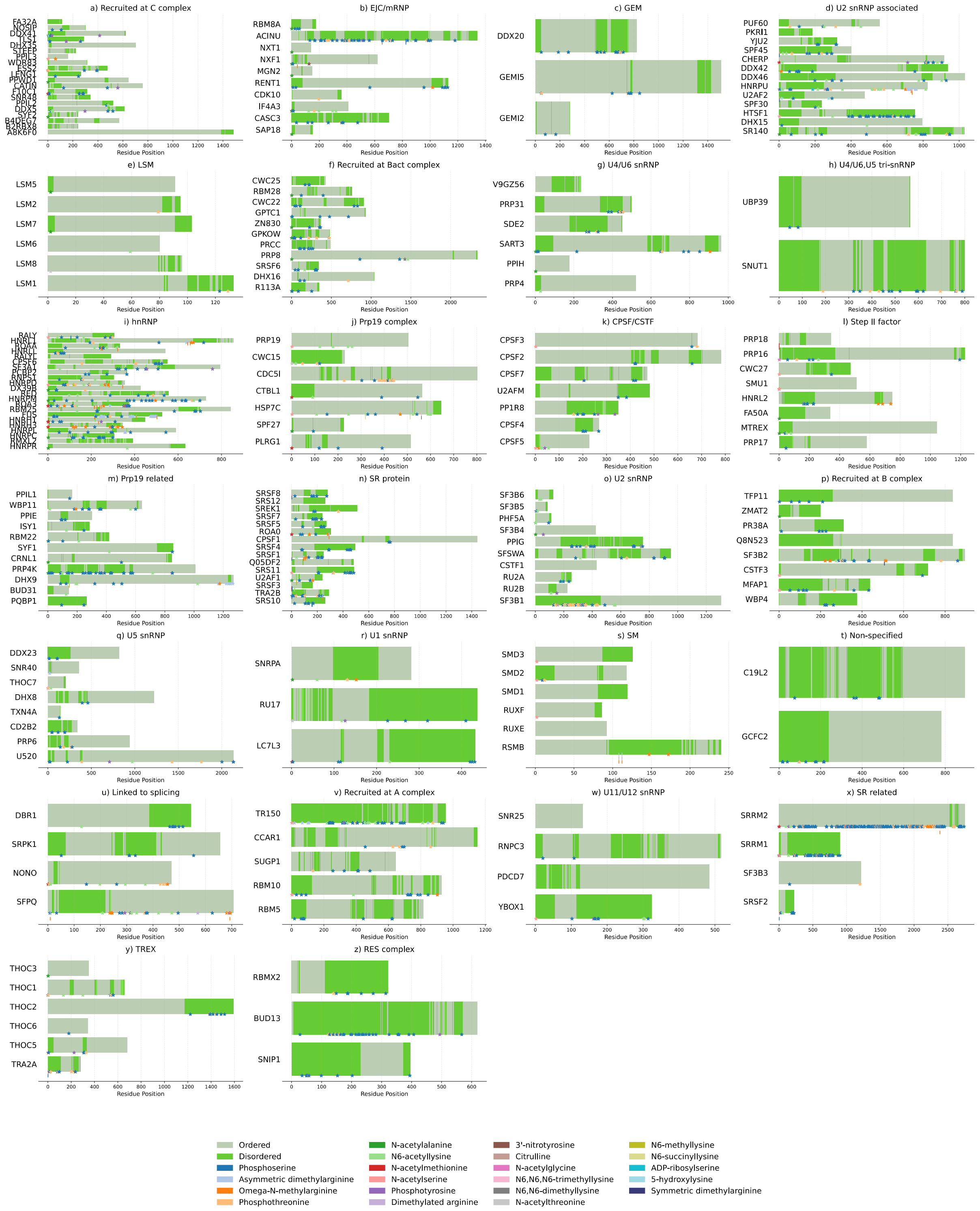


**Figure S3. Segmentation plot of post-translational modifications (PTM) in spliceosome proteins.** A-Z) Class/Family of spliceosome proteins. Each bar represents a different protein. Ordered residues are represented in grey and disordered residues are represented in green. Underlined in colored asterisks are the different types of PTM present per protein.

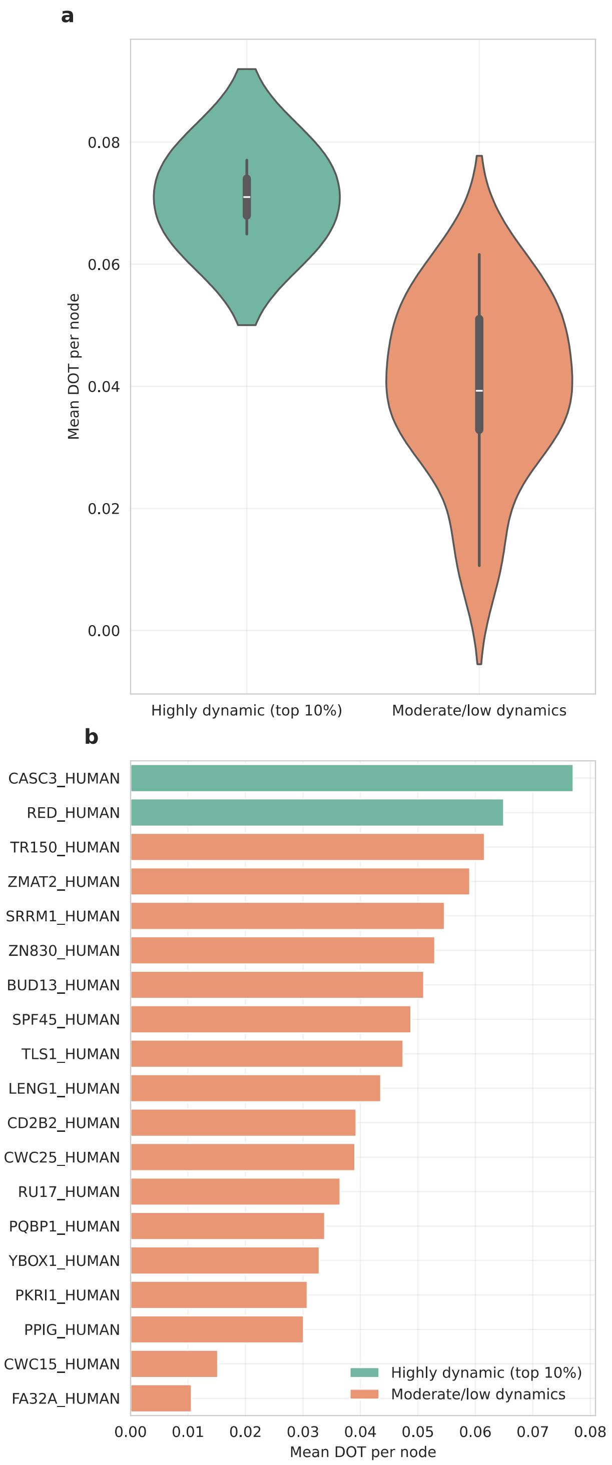


**Figure S4. Mean disorder-to-order (DOT) transition in SPL IDPs.** A) Violin plot of highly dynamic sequence versus average sequences. B) Mean DOT per protein. Green and orange refer to sequences that are highly or moderately/low dynamic, respectively.

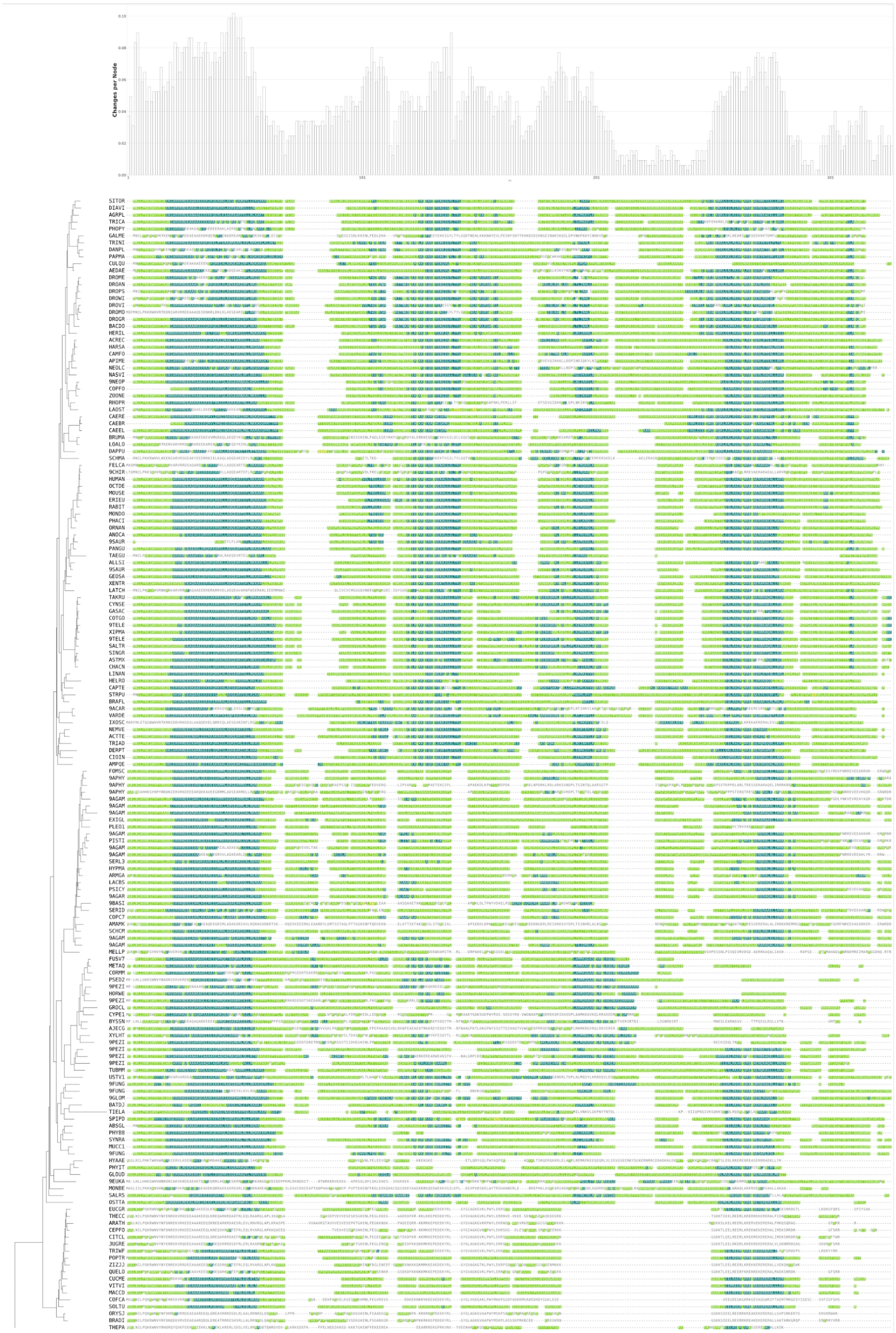

**Figure S5. Disorder content in protein LENG1_HUMAN orthologs.** The multiple sequence alignment (MSA) represents disordered (green) and ordered (not colored) regions. Disorder with secondary structure is represented by disorder with alpha-helix (blue) and beta-sheet (yellow) secondary structure. The phylogeny is shown on the right. The parsimony reconstruction of disorder-to-order transitions across the phylogenetic tree is displayed above the MSA, indicating the number of changes per site at each node.

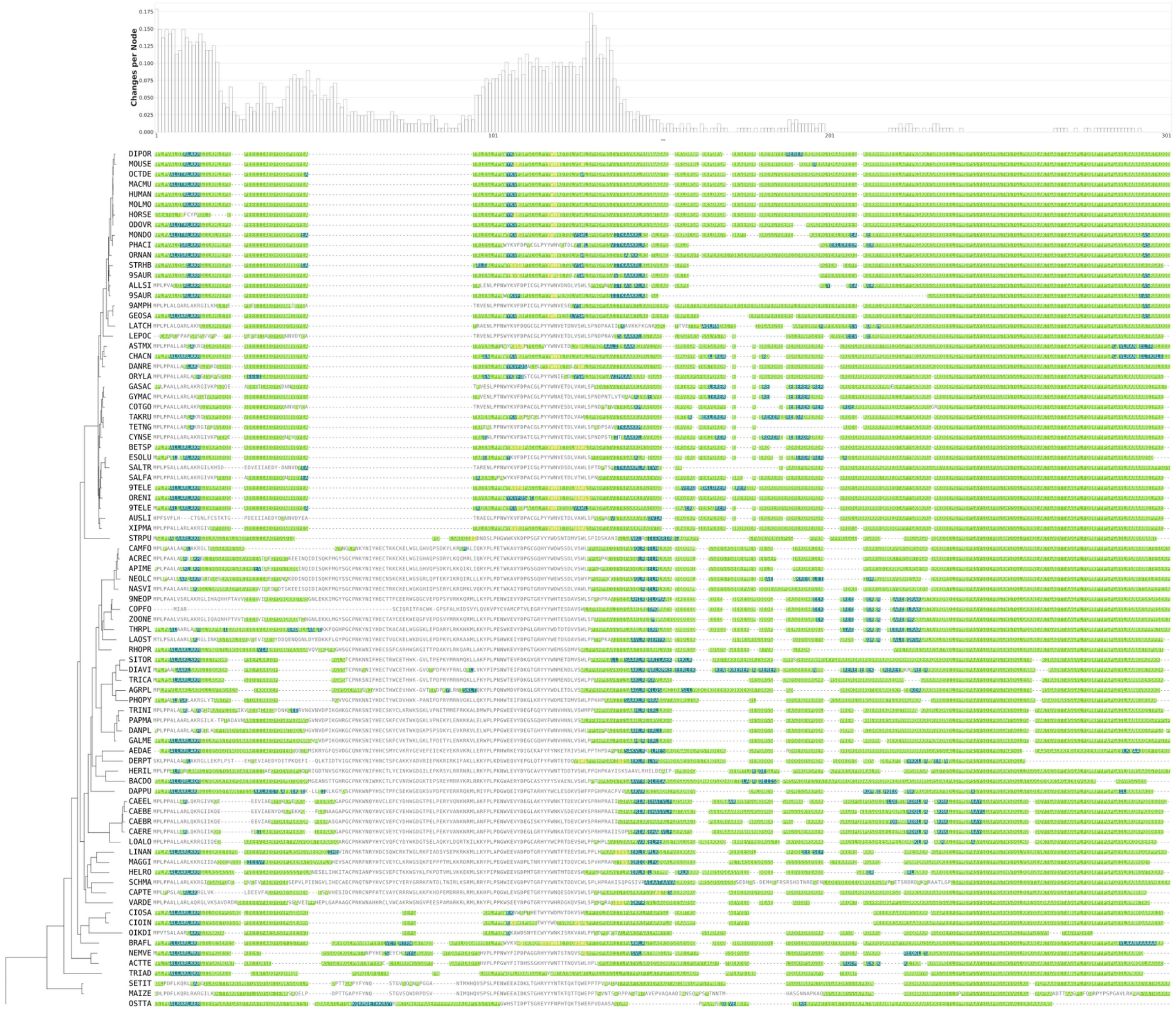

**Figure S6. Disorder content in protein PQBP1_HUMAN orthologs.** The MSA represents disordered (green) and ordered (not colored) regions. Disorder with secondary structure is represented by disorder with alpha-helical (blue) and with beta-sheet secondary structure (yellow). The phylogeny is shown on the right. The parsimony reconstruction of disorder-to-order transitions across the phylogenetic tree is displayed above the MSA, indicating the number of changes per site at each node.

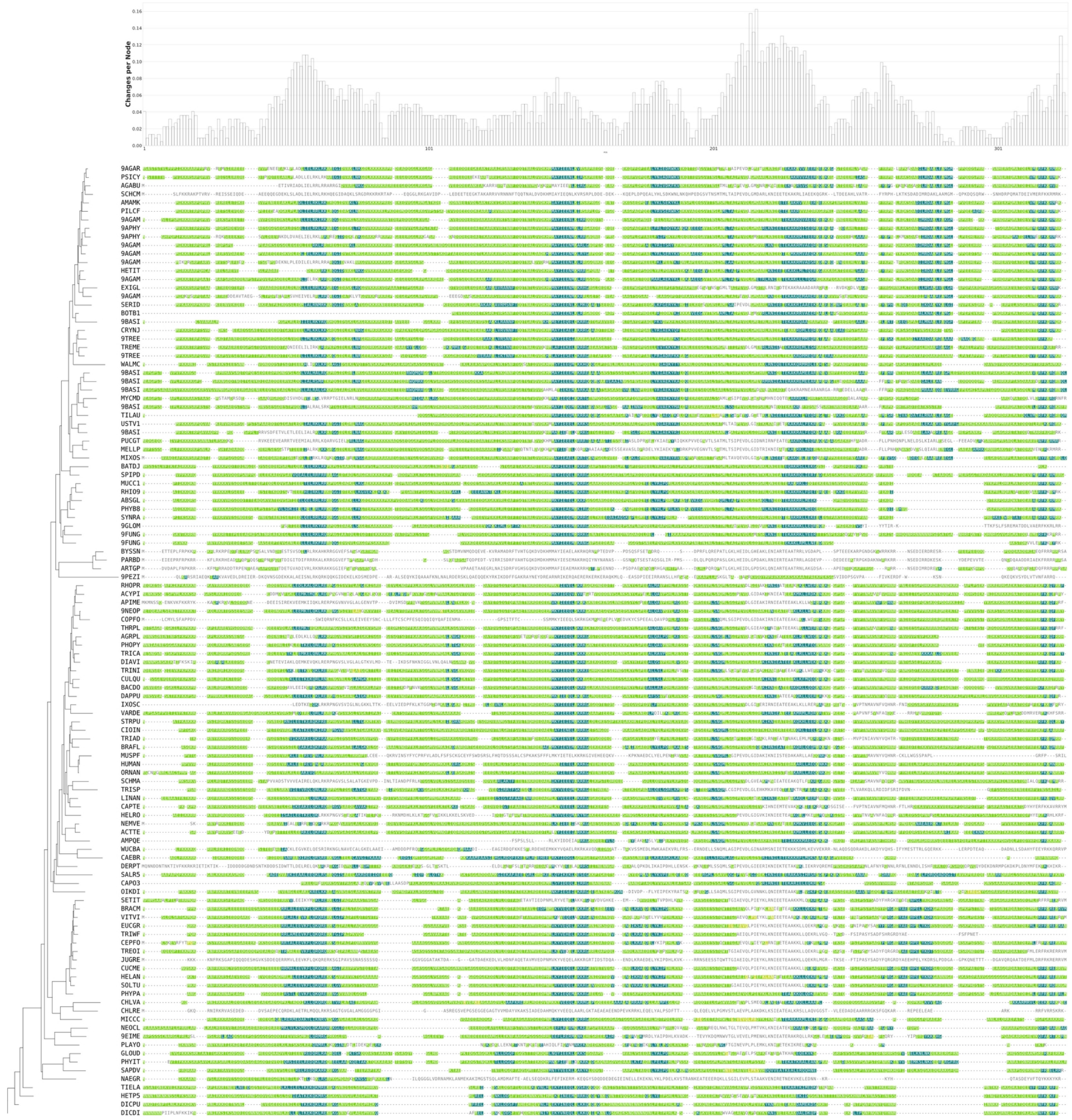

**Figure S7. Disorder content in protein TLS1_HUMAN orthologs.** The MSA represents disordered (green) and ordered (not colored) regions. Disorder with secondary structure is represented by disorder with alpha-helical (blue) and with beta-sheet secondary structure (yellow).The phylogeny is shown on the right. The parsimony reconstruction of disorder-to-order transitions across the phylogenetic tree is displayed above the MSA, indicating the number of changes per site at each node.

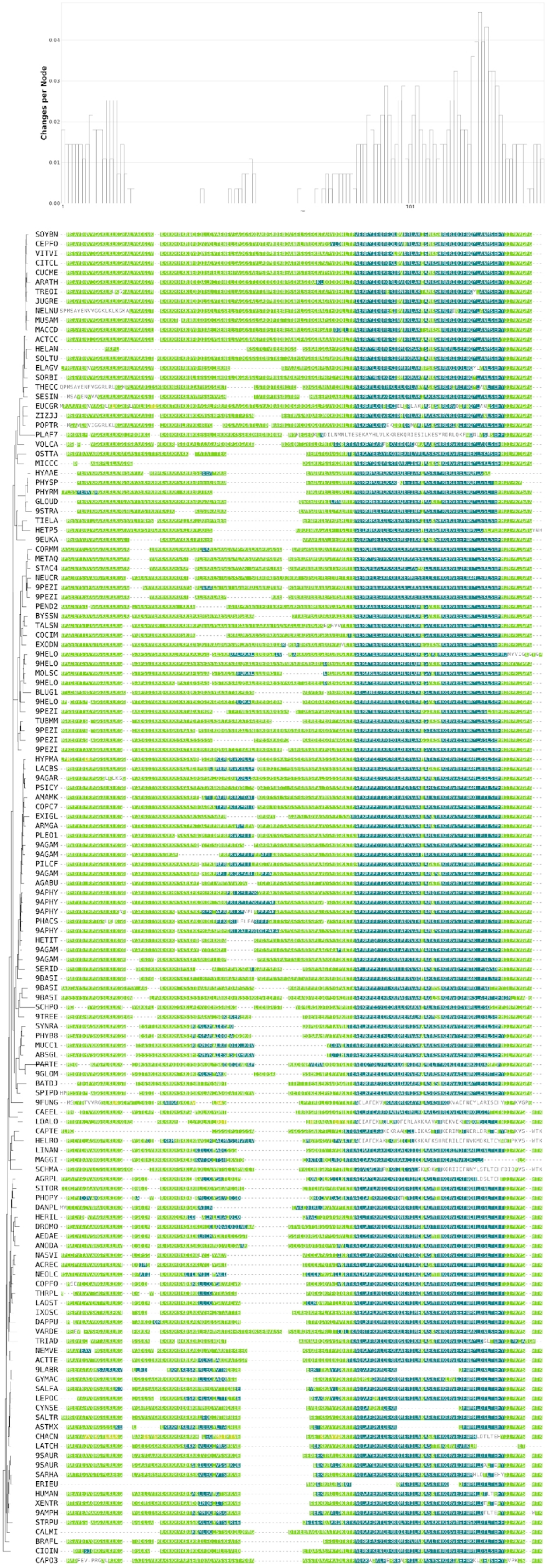

**Figure S8. Disorder content in protein FA32A_HUMAN orthologs.** The MSA represents disordered (green) and ordered (not colored) regions. Disorder with secondary structure is represented by disorder with alpha-helical (blue) and with beta-sheet secondary structure (yellow). The phylogeny is shown on the right. The parsimony reconstruction of disorder-to-order transitions across the phylogenetic tree is displayed above the MSA, indicating the number of changes per site at each node.

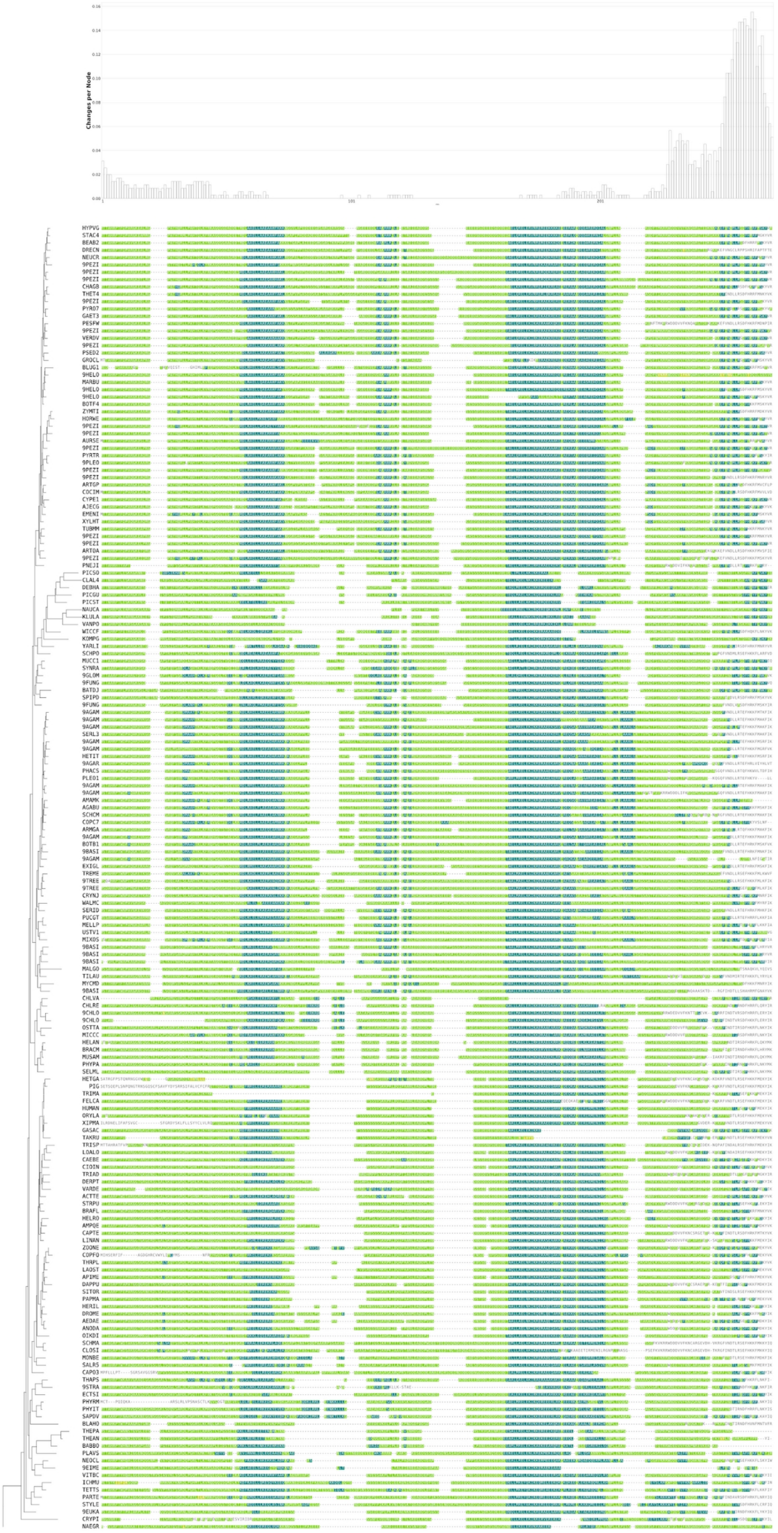

**Figure S9. Disorder content in protein CWC15_HUMAN orthologs.** The MSA represents disordered (green) and ordered (not colored) regions. Disorder with secondary structure is represented by disorder with alpha-helical (blue) and with beta-sheet secondary structure (yellow). The phylogeny is shown on the right. The parsimony reconstruction of disorder-to-order transitions across the phylogenetic tree is displayed above the MSA, indicating the number of changes per site at each node.

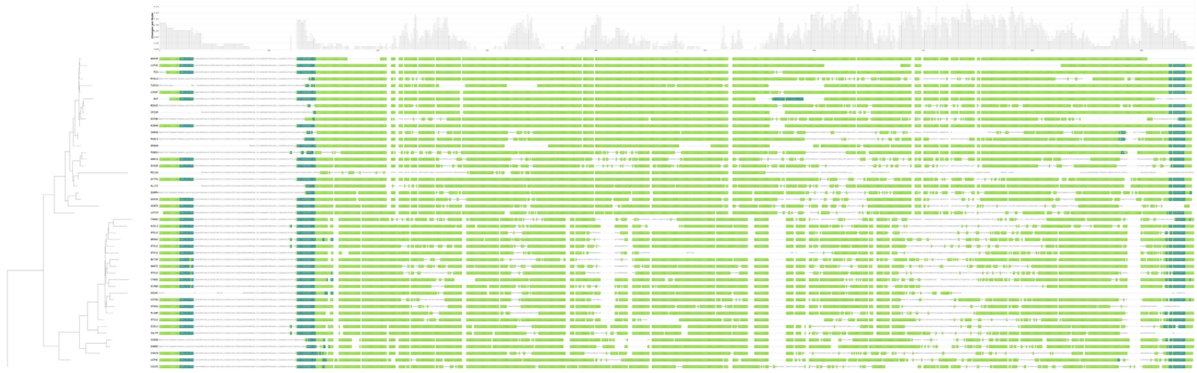

**Figure S10. Disorder content in protein SRRM1_HUMAN orthologs.** The MSA represents disordered (green) and ordered (not colored) regions. Disorder with secondary structure is represented by disorder with alpha-helical (blue) and with beta-sheet secondary structure (yellow). The phylogeny is shown on the right. The parsimony reconstruction of disorder-to-order transitions across the phylogenetic tree is displayed above the MSA, indicating the number of changes per site at each node.

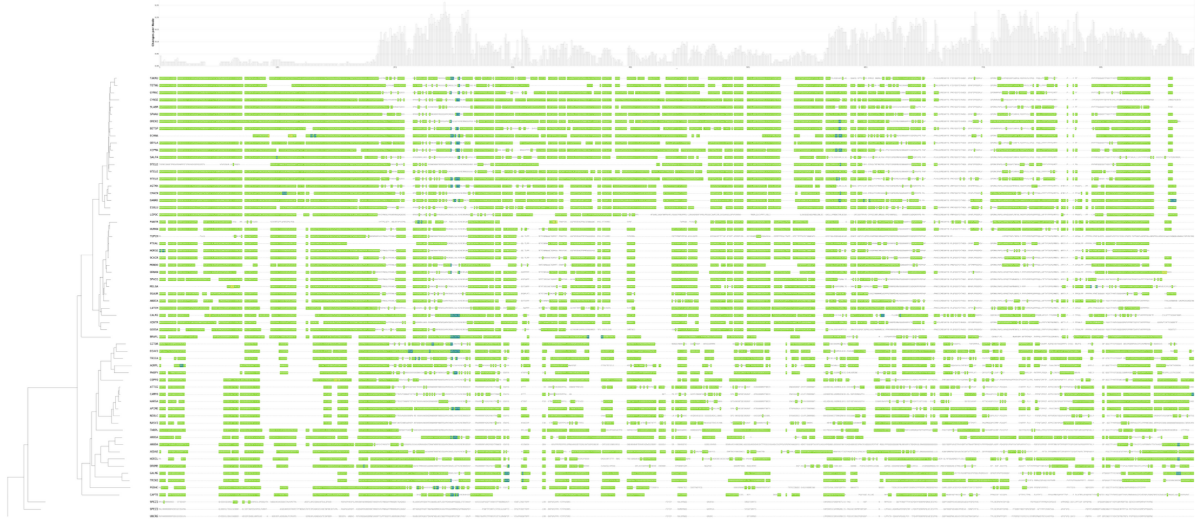

**Figure S11. Disorder content in protein CASC3_HUMAN orthologs.** The MSA represents disordered (green) and ordered (not colored) regions. Disorder with secondary structure is represented by disorder with alpha-helical (blue) and with beta-sheet secondary structure (yellow). The phylogeny is shown on the right. The parsimony reconstruction of disorder-to-order transitions across the phylogenetic tree is displayed above the MSA, indicating the number of changes per site at each node.

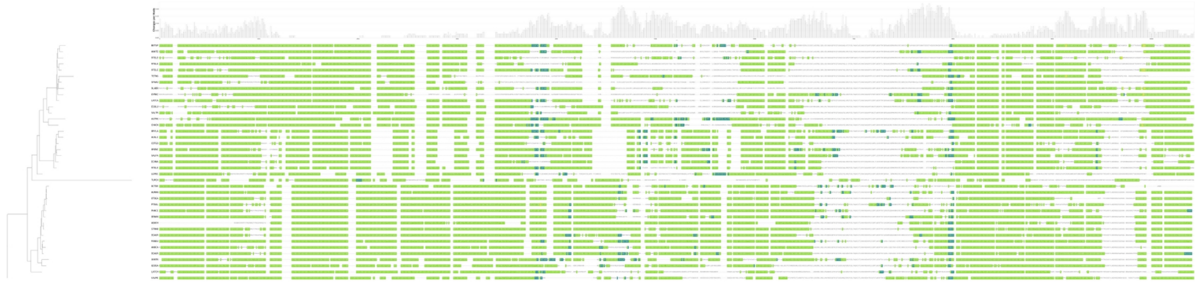

**Figure S12. Disorder content in protein TR150_HUMAN orthologs.** The MSA represents disordered (green) and ordered (not colored) regions. Disorder with secondary structure is represented by disorder with alpha-helical (blue) and with beta-sheet secondary structure (yellow). The phylogeny is shown on the right. The parsimony reconstruction of disorder-to-order transitions across the phylogenetic tree is displayed above the MSA, indicating the number of changes per site at each node.

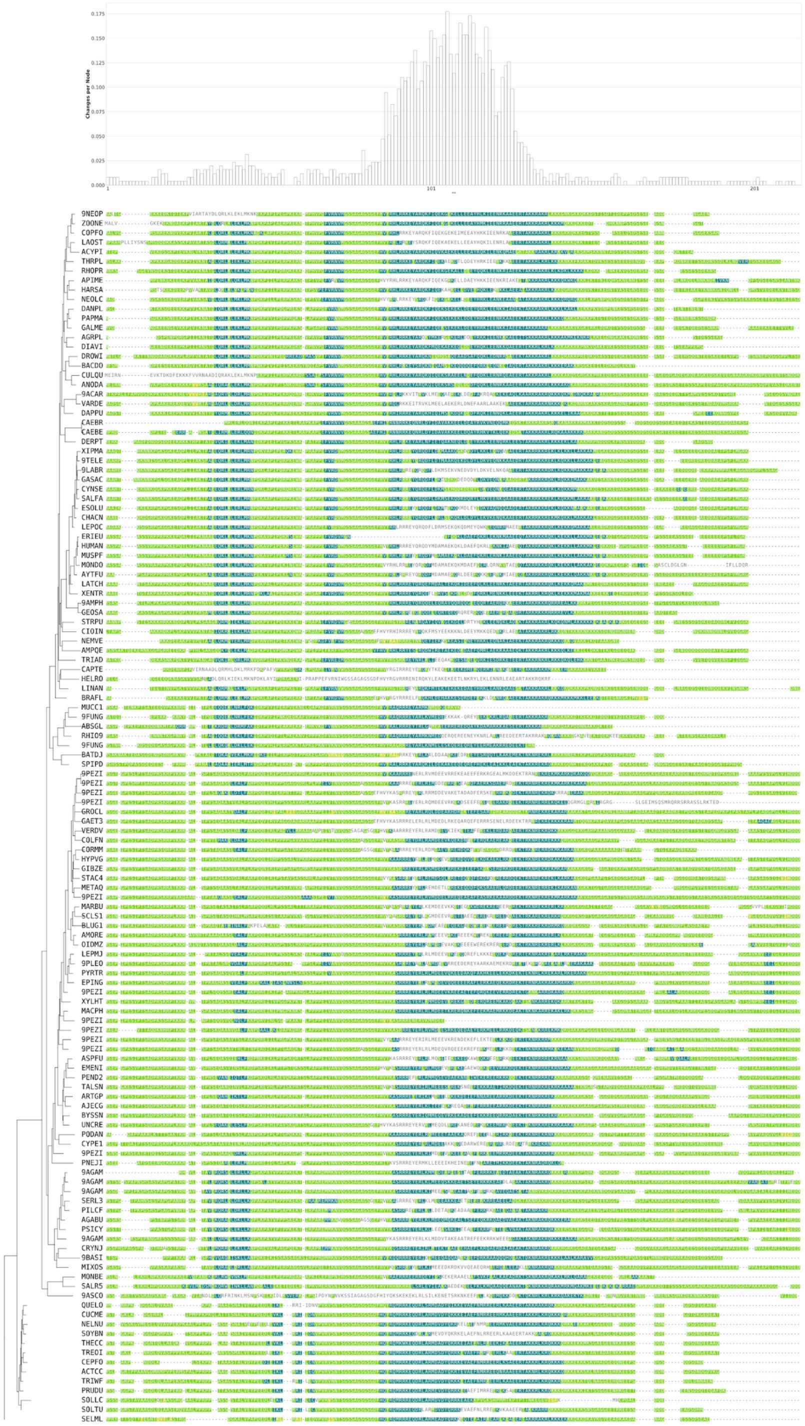

**Figure S13. Disorder content in protein PKRI1_HUMAN orthologs.** The MSA represents disordered (green) and ordered (not colored) regions. Disorder with secondary structure is represented by disorder with alpha-helical (blue) and with beta-sheet secondary structure (yellow). The phylogeny is shown on the right. The parsimony reconstruction of disorder-to-order transitions across the phylogenetic tree is displayed above the MSA, indicating the number of changes per site at each node.

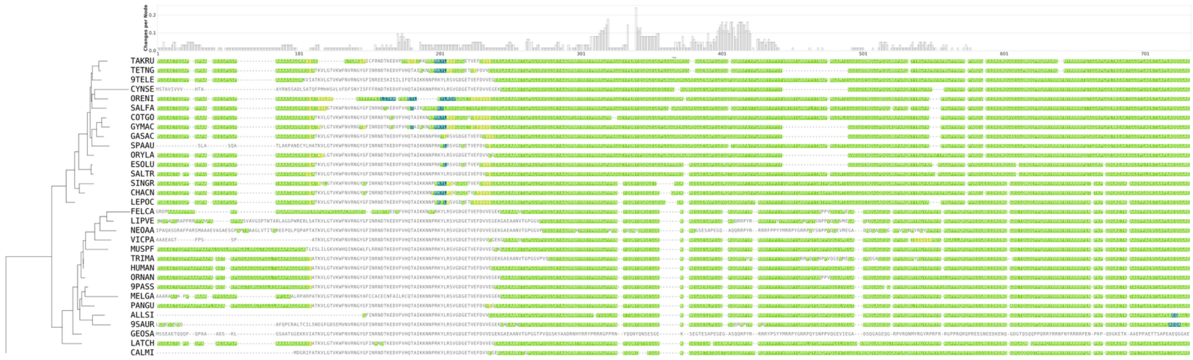

**Figure S14. Disorder content in protein YBOX1_HUMAN orthologs.** The MSA represents disordered (green) and ordered (not colored) regions. Disorder with secondary structure is represented by disorder with alpha-helical (blue) and with beta-sheet secondary structure (yellow). The phylogeny is shown on the right. The parsimony reconstruction of disorder-to-order transitions across the phylogenetic tree is displayed above the MSA, indicating the number of changes per site at each node.

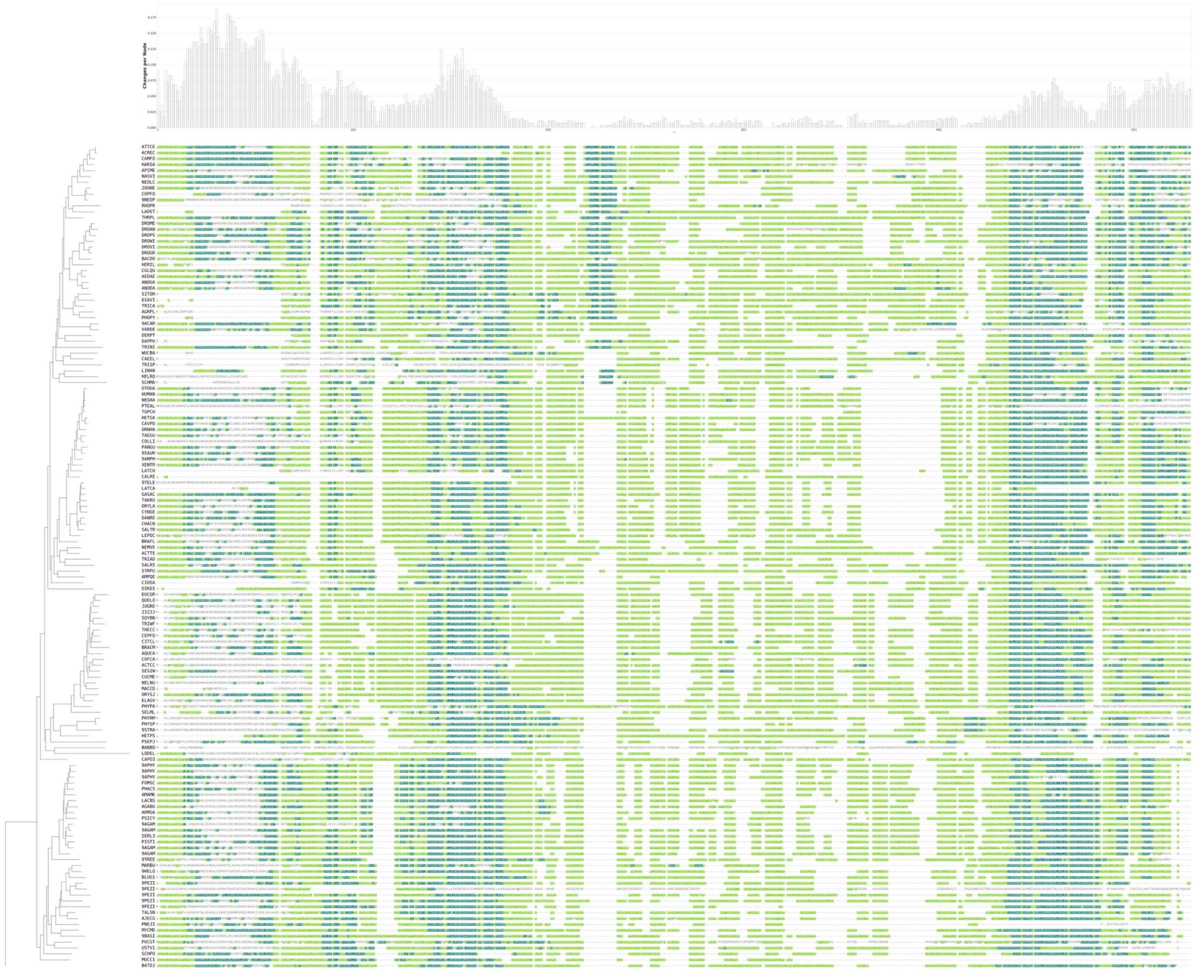

**Figure S15. Disorder content in protein CWC25_HUMAN orthologs.** The MSA represents disordered (green) and ordered (not colored) regions. Disorder with secondary structure is represented by disorder with alpha-helical (blue) and with beta-sheet secondary structure (yellow). The phylogeny is shown on the right. The parsimony reconstruction of disorder-to-order transitions across the phylogenetic tree is displayed above the MSA, indicating the number of changes per site at each node.

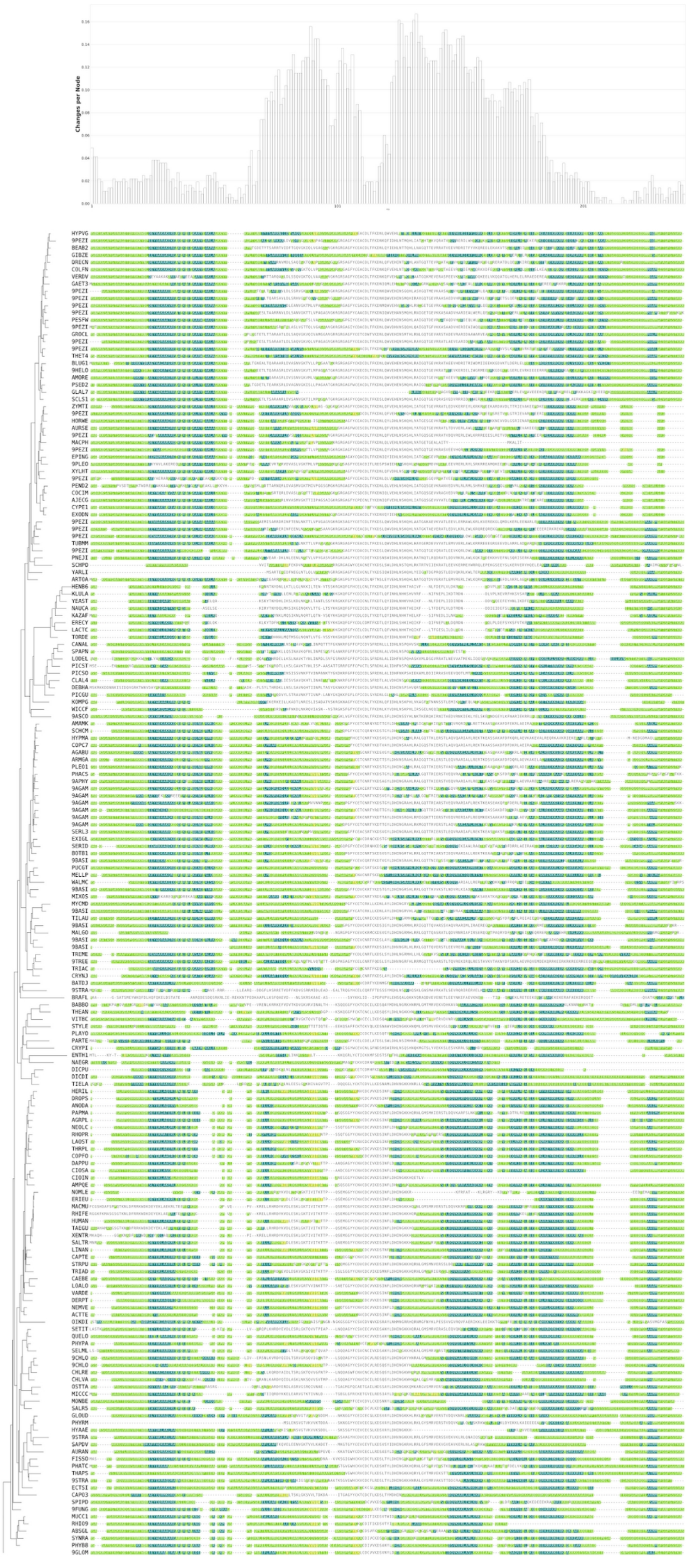


**Figure S16. Disorder content in protein ZMAT2_HUMAN orthologs.** The MSA represents disordered (green) and ordered (not colored) regions. Disorder with secondary structure is represented by disorder with alpha-helical (blue) and with beta-sheet secondary structure (yellow). The phylogeny is shown on the right. The parsimony reconstruction of disorder-to-order transitions across the phylogenetic tree is displayed above the MSA, indicating the number of changes per site at each node.

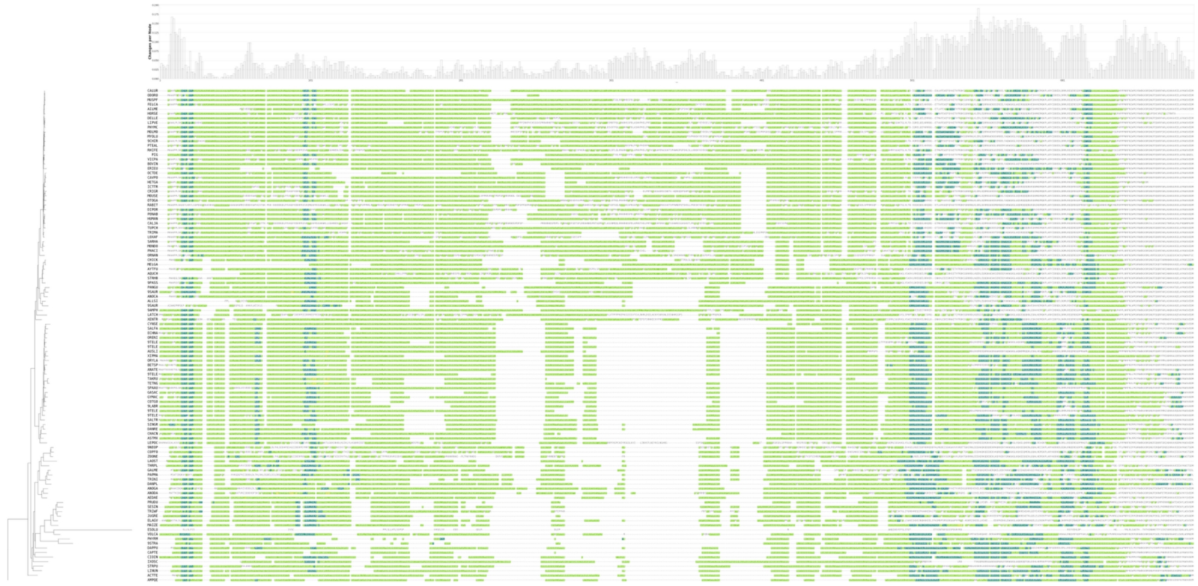

**Figure S17. Disorder content in protein BUD13_HUMAN orthologs.** The MSA represents disordered (green) and ordered (not colored) regions. Disorder with secondary structure is represented by disorder with alpha-helical (blue) and with beta-sheet secondary structure (yellow). The phylogeny is shown on the right. The parsimony reconstruction of disorder-to-order transitions across the phylogenetic tree is displayed above the MSA, indicating the number of changes per site at each node.

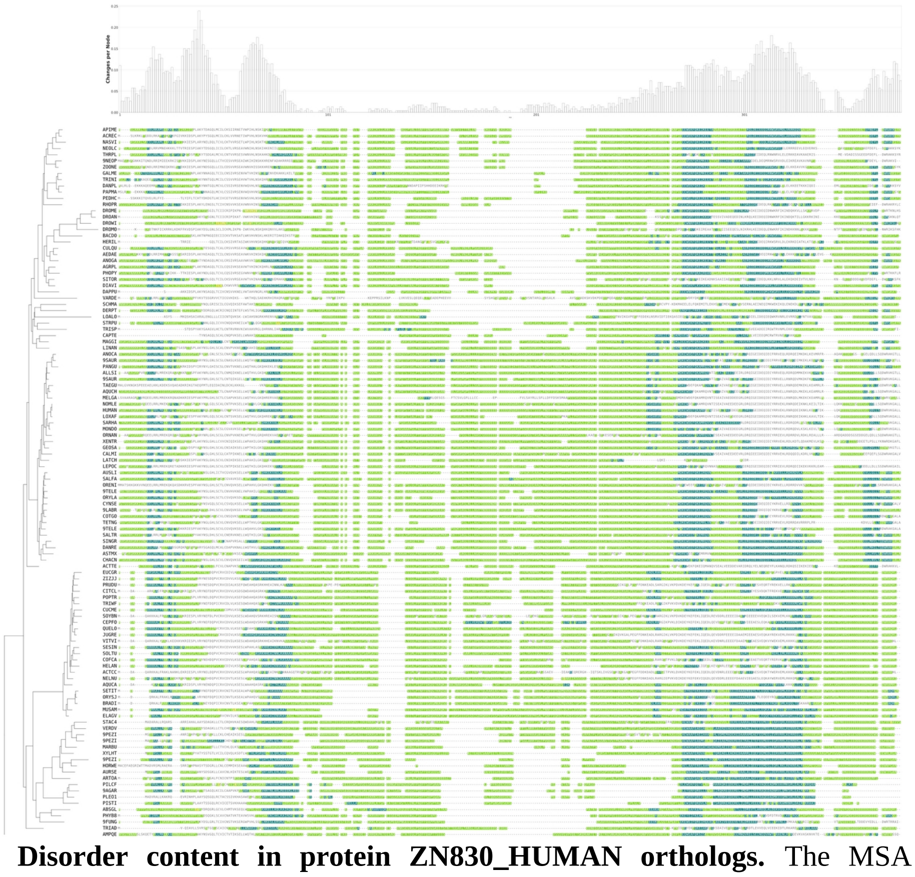

**Figure S18. Disorder content in protein ZN830_HUMAN orthologs.** The MSA represents disordered (green) and ordered (not colored) regions. Disorder with secondary structure is represented by disorder with alpha-helical (blue) and with beta-sheet secondary structure (yellow). The phylogeny is shown on the right. The parsimony reconstruction of disorder-to-order transitions across the phylogenetic tree is displayed above the MSA, indicating the number of changes per site at each node.

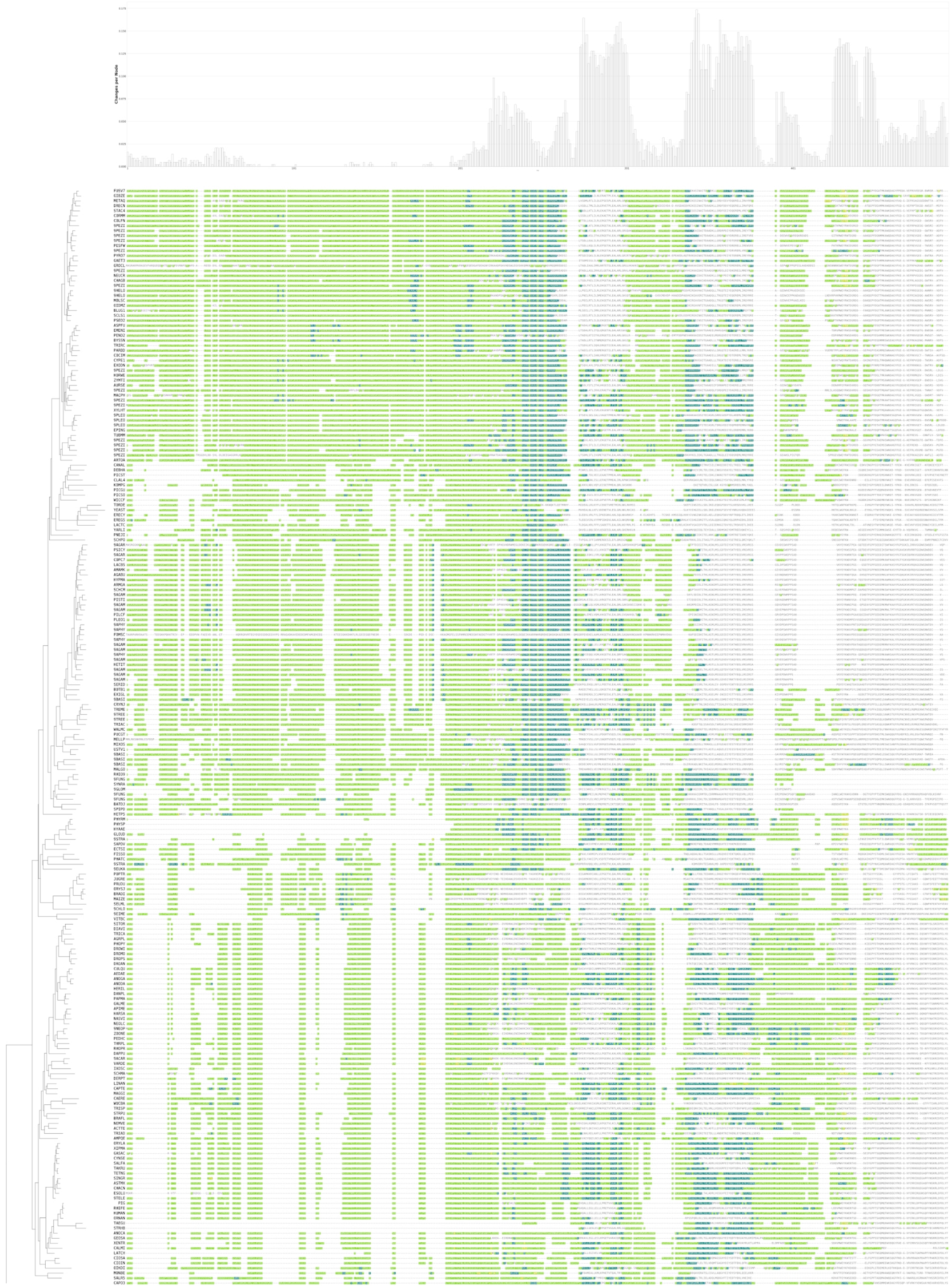

**Figure S19. Disorder content in protein CD2B2_HUMAN orthologs.** The MSA represents disordered (green) and ordered (not colored) regions. Disorder with secondary structure is represented by disorder with alpha-helical (blue) and with beta-sheet secondary structure (yellow). The phylogeny is shown on the right. The parsimony reconstruction of disorder-to-order transitions across the phylogenetic tree is displayed above the MSA, indicating the number of changes per site at each node.

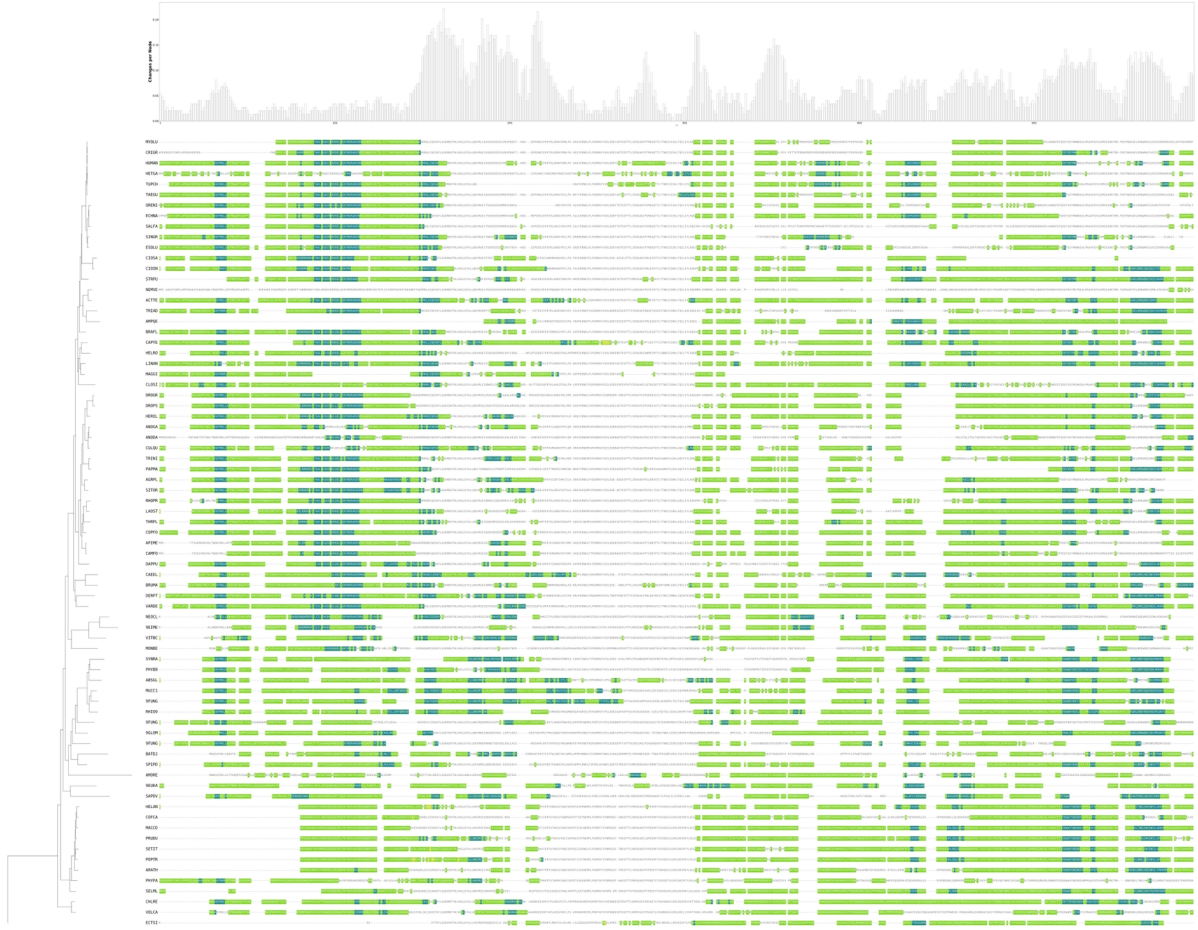

**Figure S20. Disorder content in protein RED_HUMAN orthologs.** The MSA represents disordered (green) and ordered (not colored) regions. Disorder with secondary structure is represented by disorder with alpha-helical (blue) and with beta-sheet secondary structure (yellow). The phylogeny is shown on the right. The parsimony reconstruction of disorder-to-order transitions across the phylogenetic tree is displayed above the MSA, indicating the number of changes per site at each node.

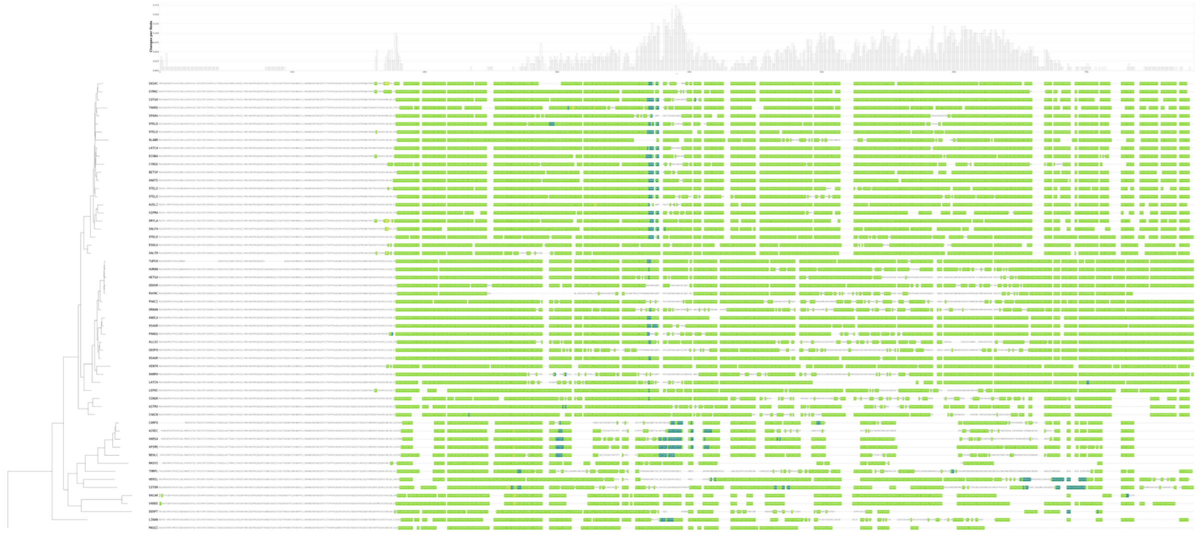

**Figure S21. Disorder content in protein PPIG_HUMAN orthologs.** The MSA represents disordered (green) and ordered (not colored) regions. Disorder with secondary structure is represented by disorder with alpha-helical (blue) and with beta-sheet secondary structure (yellow). The phylogeny is shown on the right. The parsimony reconstruction of disorder-to-order transitions across the phylogenetic tree is displayed above the MSA, indicating the number of changes per site at each node.

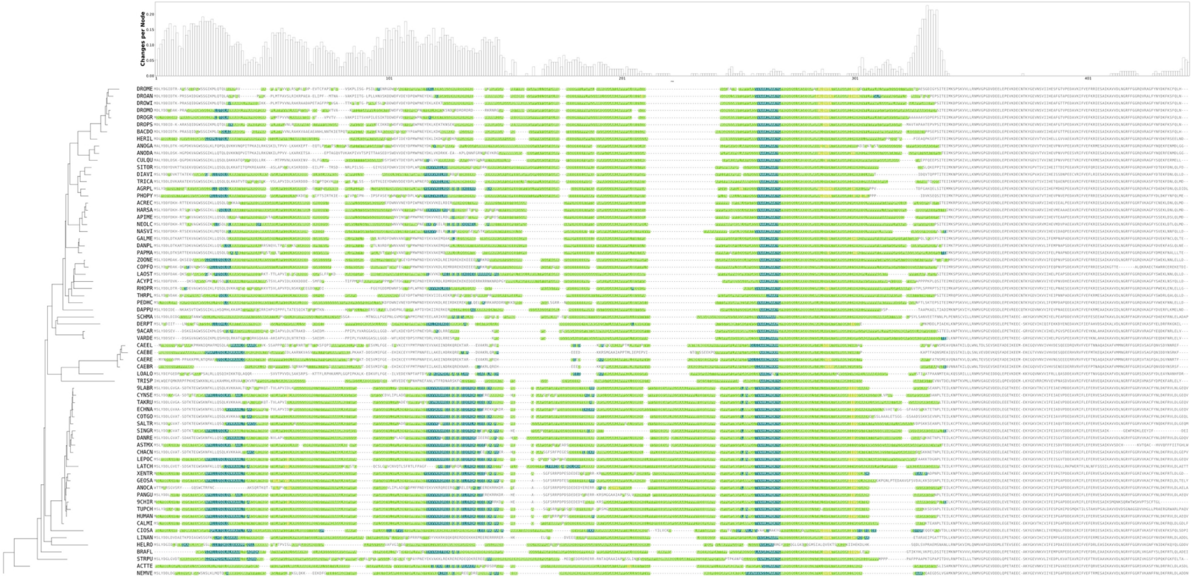

**Figure S22. Disorder content in protein SPF45_HUMAN orthologs.** The MSA represents disordered (green) and ordered (not colored) regions. Disorder with secondary structure is represented by disorder with alpha-helical (blue) and with beta-sheet secondary structure (yellow). The phylogeny is shown on the right. The parsimony reconstruction of disorder-to-order transitions across the phylogenetic tree is displayed above the MSA, indicating the number of changes per site at each node.

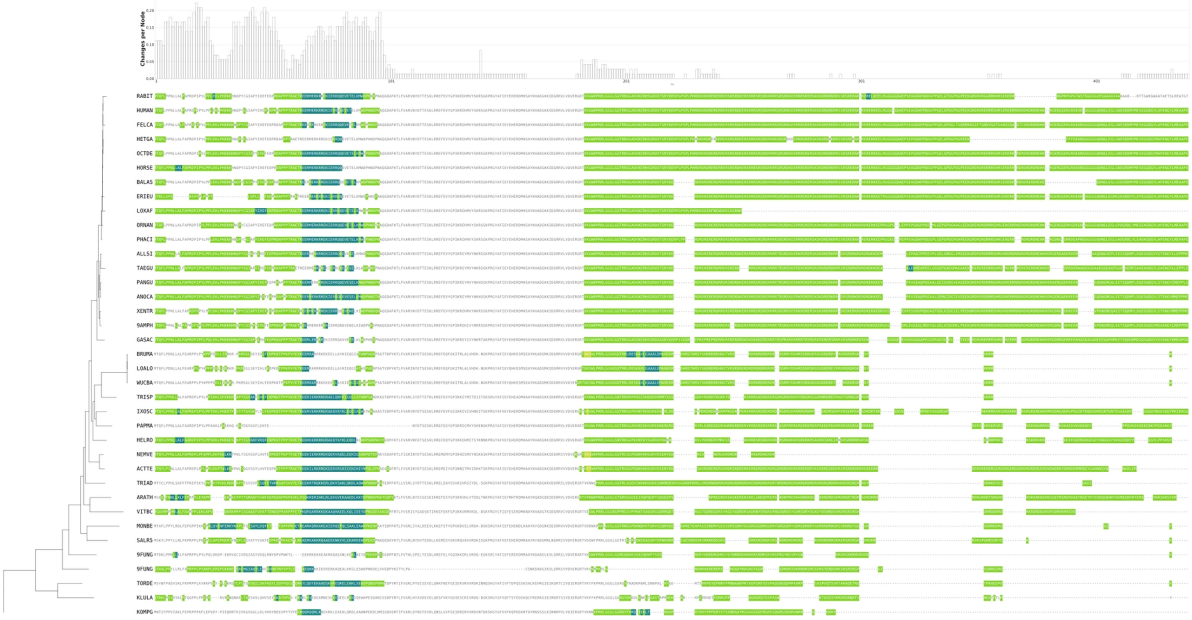

**Figure S23. Disorder content in protein RU17_HUMAN orthologs.** The MSA represents disordered (green) and ordered (not colored) regions. Disorder with secondary structure is represented by disorder with alpha-helical (blue) and with beta-sheet secondary structure (yellow). The phylogeny is shown on the right. The parsimony reconstruction of disorder-to-order transitions across the phylogenetic tree is displayed above the MSA, indicating the number of changes per site at each node.

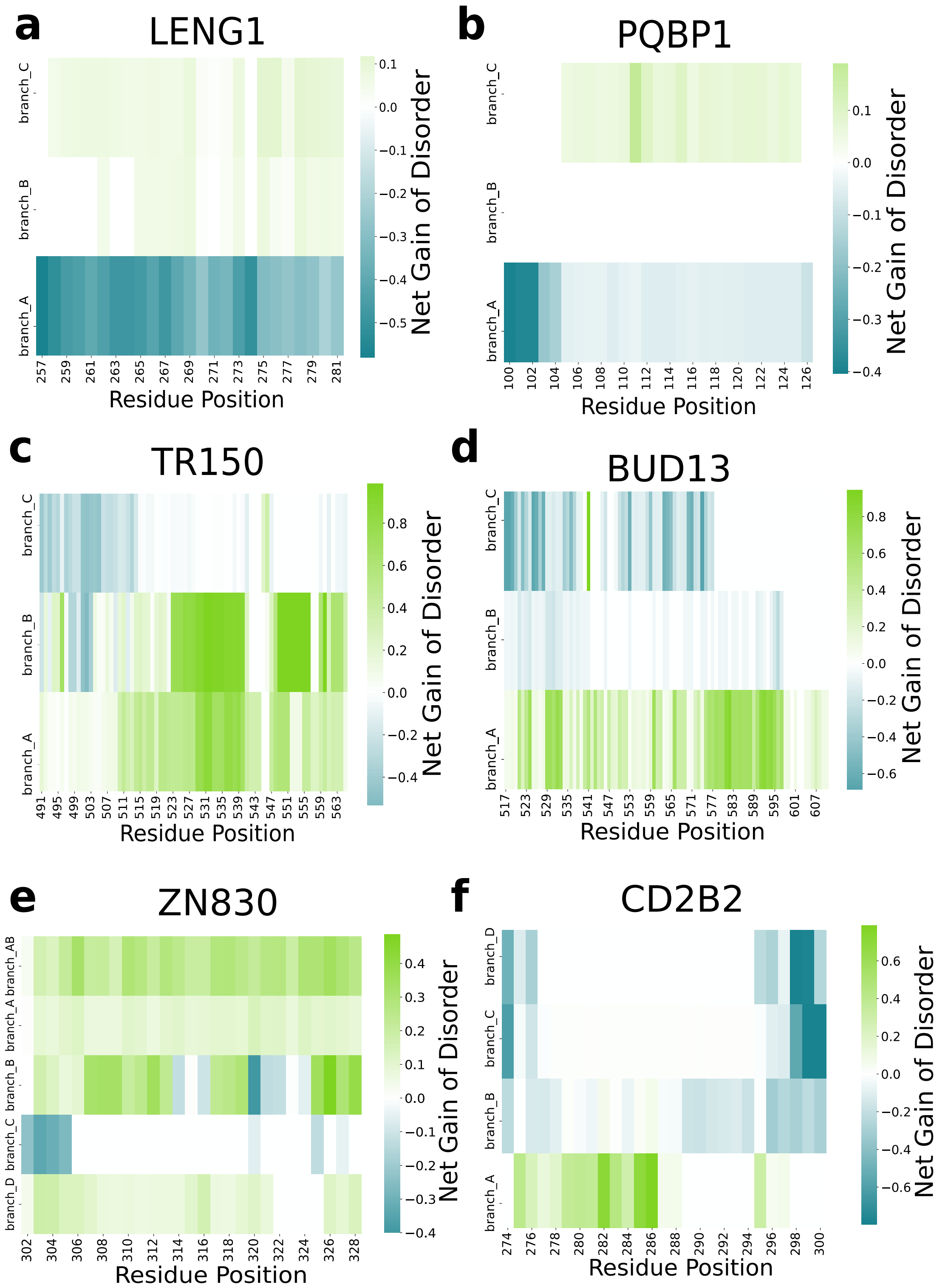


**Figure S24. Disorder-to-order transitions in SPL proteins.** Heatmap with the net gain of disorder per residue position. In blue and green are shown ordered regions and disordered regions, respectively.

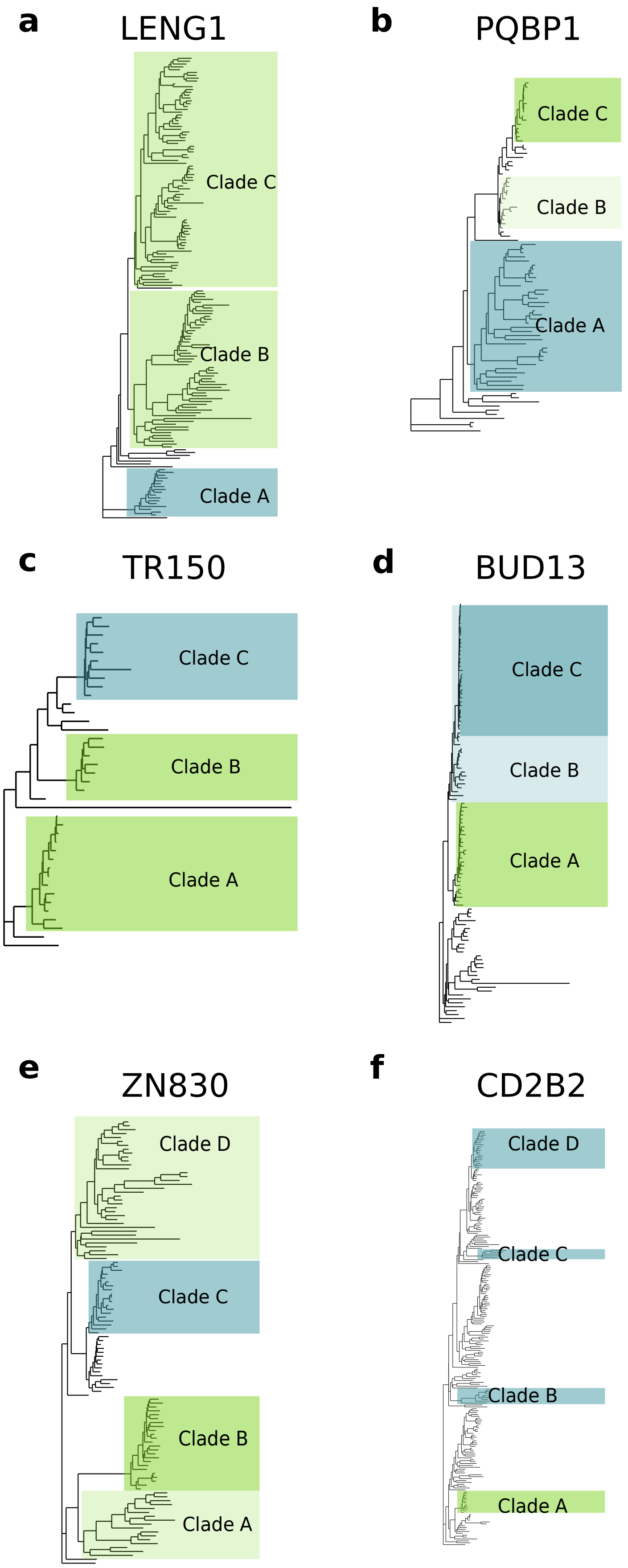


**Figure S25. Disorder-to-order evolution in SPL proteins.** Phylogenetic trees, highlighting the clades evaluated. In blue and green are shown ordered regions and disordered regions, respectively.

### Supplementary Tables

**Table S1. CB-IDR composition relative abundance.**

| Class/CB-IDR | RS-like | Poly P/Q | Charged res. | Non-charged res. | G-rich |
|---|---|---|---|---|---|
| RES complex | 23.99 | 4.22 | 14.72 | - | - |
| U1 snRNP | 7.83 | 2.82 | 36.36 | 11.2 | - |
| SR | 57.91 | - | 12.19 | - | 4.3 |
| Recruited at B complex | 2.94 | 4.76 | 32.19 | - | - |
| U2 snRNP associated | 8.63 | 5.87 | 27.02 | - | - |
| Recruited at $B^{act}$ complex | 12.49 | 6.3 | 10.43 | - | - |

**Table S2. Post translational modifications (PTMs) in Spliceosome Proteins with High Disorder Content (> 70%) ordered by PTM amount.**

| Protein | Class/ Family | Disorder % | PTM Type | PTM Position |
|---|---|---|---|---|
| SRRM1 | SR related | 89.49 | Phosphoserine | 227, 234, 240, 260, 389, 391, 393, 402, 414, 420, 429, 431, 436, 450, 452, 463, 465, 478, 524, 526, 528, 530, 532, 549, 551, 555, 560, 562, 583, 597, 605, 607, 616, 626, 628, 636, 638, 694, 695, 696, 705, 707, 713, 715, 738, 740, 748, 752, 754, 756, 769, 773, 775, 777, 781, 791, 795, 797, 802, 874,901, |
| | | | Phosphothreonine | 220, 241, 406, 416, 555, 572, 574, 581, 614, 718, 778, 793, 872 |
| | | | N-acetylmethionine | 1 |
| | | | Citrulline | 7 |
| | | | N6-acetyllysine | 140 |
| TR150 | Recruited at A complex | 86.07 | Phosphoserine | 220, 232, 237, 240, 243, 248, 253, 257, 315, 320, 323, 326, 339, 377, 379, 406, 408, 444, 468, 535, 575, 619, 622, 698, 928, 939 |
| | | | N6-acetyllysine | 221, 346, 455, 470, 481, 519, 527, 558, 811 |
| | | | Dimethylated arginine | 17, 66, 101, 108, 845 |
| | | | Phosphothreonine | 324, 397, 874 |
| | | | N-acetylserine | 2 |
| FUS | hnRNP | 89.35 | Dimethylarginine | 216, 218, 242, 244, 248, 251, 259, 377, 383, 386, |

| | | | | |
|---|---|---|---|---|
| | | | | 388, 394, 407, 473, 476, 481, 485, 487, 491, 495, 498, 503 |
| | | | Phosphoserine | 26,30, 42, 221, 277, 340 |
| | | | Omega-N-methylarginine | 216, 218, 407, 503 |
| BUD13 | RES complex | 80.61 | Phosphoserine | 28, 127, 139, 151, 163, 172, 175, 197, 201, 214, 222, 226, 235, 236, 240, 248, 258, 259, 271, 281, 325, 354, 357, 358, 391, 407, 567 |
| | | | Phosphothreonine | 123, 135, 147, 159 |
| YBOX1 | U11/U12 snRNP | 81.48 | Phosphoserine | 165, 167, 174, 176, 314 |
| | | | N-acetylserine | 2, 301, 304 |
| | | | N6-acetyllysine | 301, 304 |
| | | | Phosphotyrosine | 162 |
| CASC3 | EJC/mRNP | 86.91 | Phosphoserine | 35, 117, 148, 265, 363, 373, 477 |
| | | | Phosphothreonine | 357 |
| CWC15 | Prp19 complex | 95.20 | Phosphothreonine | 47, 110 |
| | | | N-acetylthreonine | 2 |
| | | | N6-acetyllysine | 18 |
| | | | Phosphoserine | 121 |
| TLS1 | Recruited at C complex | 100 | Phosphoserine | 15, 17, 261 |
| | | | Phosphotyrosine | 147 |
| | | | Phosphothreonine | 253 |
| PQBP1 | Prp19 related | 100 | Phosphoserine | 94, 247, 946, 948 |
| CWC25 | Recruited at Bact complex | 81.18 | Phosphoserine | 170, 218, 222 |
| LENG1 | Recruited at C complex | 100 | Phosphoserine | 59, 245 |
| SRSF3 | SR protein | 100 | - | - |
| FA32A | Recruited at C complex | 95.54 | - | - |
| PKRI1 | U2 snRNP associated | 82.07 | - | - |
| ZMAT2 | Recruited at B complex | 80.90 | N-acetylalanine | 2 |

**Table S3. Spliceosome proteins involved in tumor biogenesis (complete).**

| Gene Symbol | Mutation AA | Tumor Location |
|---|---|---|
| **ACIN1** | ['p.I271M', 'p.I253M', 'p.R646_S647dup', 'p.A407P', 'p.R1102Q', 'p.E281del', 'p.S478F', 'p.R588_S589dup', | Soft tissue; Gastrointestinal |

| | | |
|---|---|---|
| | 'p.S603_R604insHS', 'p.R1160Q', 'p.S77N', 'p.A389P', 'p.S135N', 'p.I311M', 'p.E223del', 'p.S585_R586insHS', 'p.S409P', 'p.R402Q', 'p.S467P', 'p.R401Q', 'p.E241del', 'p.A447P', 'p.S643_R644insHS', 'p.S438F', 'p.R433Q', 'p.S420F', 'p.R606_S607dup', 'p.R1147Q', 'p.R1120Q', 'p.S427P', nan] | stromal tumour |
| **BCAS2** | ['p.N139S', 'p.?', nan] | Large intestine; Carcinoma; Adenocarcinoma |
| **BUD13** | ['p.P190S', 'p.*620Yext*3', 'p.*486Yext*3', nan] | Soft tissue; Gastrointestinal stromal tumour |
| **BUD31** | ['p.?', nan] | Liver; Carcinoma; Hepatocellular carcinoma |
| **C9orf78** | ['p.?', nan] | Liver; Carcinoma; Hepatocellular carcinoma |
| **CACTIN** | ['p.?', 'p.A499V', 'p.H276Y', 'p.R378P', nan] | Soft tissue; Gastrointestinal stromal tumour |
| **CASC3** | [nan] | Central nervous system; Primitive neuroectodermal tumour-medulloblastoma; classic |
| **CCAR1** | ['p.P344S', 'p.E607K', 'p.?', 'p.P329S', 'p.E592K', nan] | Skin; Malignant melanoma; desmoplastic |
| **CD2BP2** | ['p.T88S', 'p.E17*', 'p.F329L', nan] | Soft tissue; Gastrointestinal stromal tumour |
| **CDC40** | ['p.?', nan] | Lung; Carcinoma; Small Cell Carcinoma |
| **CDK10** | ['p.L202P', 'p.?', 'p.L131P', 'p.L155P', nan] | Breast; Carcinoma; Basal (triple-negative) Carcinoma |
| **CHERP** | ['p.Q352del', 'p.Q341del', 'p.L47I', nan] | Upper Aerodigestive Tract; Carcinoma; Squamous Cell Carcinoma |
| **CPSF1** | ['p.V1029L', 'p.T838M', 'p.T561I', nan] | Soft Tissue; Gastrointestinal stromal tumour |
| **CPSF2** | ['p.S322=', nan] | Breast; Carcinoma; luminal NS carcinoma |

| | | |
|---|---|---|
| **CPSF3** | ['p.?', 'p.D578N', 'p.*648Sext*6', 'p.P554T', 'p.I101V', 'p.I138V', 'p.E99Q', 'p.D541N', 'p.*685Sext*6', 'p.P591T', 'p.E136Q', nan] | Upper Aerodigestive Tract; Carcinoma; Squamous Cell Carcinoma |
| **CPSF4** | ['p.?', 'p.V146I', 'p.V166I', 'p.V224I', 'p.V199I', nan] | Liver; Carcinoma; Hepatocellular Carcinoma |
| **CPSF6** | [nan] | Central nervous System; Primitive Neuroectodermal tumour-medulloblastoma; Classic |
| **CPSF7** | ['p.L223F', 'p.L275F', 'p.L232F', nan] | Central Nervous System;glioma; Astrocytoma Grade IV |
| **CRNKL1** | ['p.E737K', 'p.E181Q', 'p.Q111H', 'p.F39L', 'p.F51L', 'p.E576K', 'p.R700W', 'p.E342Q', 'p.E330Q', 'p.G35R', 'p.P124S', 'p.P112S', 'p.A548T', 'p.A560T', 'p.T158A', 'p.?', 'p.R539W', 'p.E725K', 'p.T146A', 'p.R688W', 'p.Q99H', 'p.A399T', nan] | Urinary Tract; Carcinoma |
| **CSTF1** | ['p.G33=', nan] | Central Nervous System; Glioma; Astrocytoma Grade IV |
| **CSTF3** | [nan] | Central nervous system; Primitive neuroectodermal tumour-medulloblastoma; Classic |
| **CTNNBL1** | [nan] | Central nervous System; Primitive neuroectodermal tumour-medulloblastoma; Classic |
| **CWC22** | ['p.S773R', 'p.R742K', nan] | Soft tissue; Gastrointestinal Stromal Tumour |
| **CWC27** | ['p.V335*', 'p.?', 'p.P256A', nan] | Oesophagus; Carcinoma; Squamous Cell Carcinoma |
| **DDX20** | ['p.V250=', 'p.I244T', 'p.I636T', 'p.L536V', 'p.L144V', 'p.V642=', nan] | Upper Aerodigestive Tract; Carcinoma; Squamous Cell Carcinoma |
| **DDX23** | ['p.S683F', 'p.S39P', 'p.R355Lfs*8', 'p.?', 'p.L309F', | Breast; Carcinoma; |

| | | |
|---|---|---|
| | 'p.L247=', 'p.R374C', 'p.K271R', 'p.D95H', 'p.D33E', 'p.I284N', nan] | Ductal Carcinoma |
| **DDX39B** | ['p.?', nan] | Liver; Carcinoma; Hepatocellular Carcinoma |
| **DDX41** | ['p.R311Q', 'p.A133T', 'p.A25T', 'p.?', nan] | Central nervous system; Glioma; Astrocytoma Grade IV |
| **DDX42** | ['p.E769del', 'p.E888del', 'p.N24I', 'p.?', 'p.N143I', nan] | Lung; Carcinoma; Adenocarcinoma |
| **DDX46** | ['p.E594=', nan] | Lung; Carcinoma; Small Cell Carcinoma |
| **DDX5** | ['p.?', 'p.Q209=', 'p.I153=', 'p.R240K', 'p.Y244=', 'p.Y165=', 'p.P102S', 'p.E116K', 'p.L245F', 'p.H348Y', 'p.K343E', 'p.H427Y', 'p.K264E', 'p.P97L', 'p.I169V', 'p.V453=', 'p.E408K', 'p.Q288=', 'p.L166F', 'p.P97S', 'p.E329K', 'p.H389Y', 'p.R161K', 'p.H310Y', 'p.E92=', 'p.V374=', 'p.I90V', nan] | Soft tissue; haemangioblastoma |
| **DHX15** | ['p.?', 'p.V713E', nan] | Liver; Carcinoma; Hepatocellular carcinoma |
| **DHX16** | ['p.S348Y', 'p.K814T', 'p.K333T', nan] | Kidney; Carcinoma; Renal Cell |
| **DHX35** | ['p.D186Y', 'p.?', 'p.A675S', 'p.T225=', 'p.T194=', 'p.A651S', 'p.D155Y', 'p.A644S', nan] | Liver; Carcinoma; Hepatocellular carcinoma |
| **DHX38** | ['p.?', 'p.D630=', 'p.T16S', nan] | Liver; Carcinoma; Hepatocellular carcinoma |
| **DHX8** | ['p.R256H', 'p.R978C', 'p.T599=', 'p.L1006M', 'p.F448=', nan] | Haematopoietic and Lymphoid Tissue; Lymphoid Neoplasm; Acute Lymphoblastic T Cell Leukaemia |
| **DHX9** | ['p.D511N', 'p.T443I', 'p.I478M', 'p.A61T', 'p.E515K', 'p.E629Q', 'p.Y1167*', nan] | Eye; Malignant melanoma; Spindle |
| **EIF4A3** | ['p.?', nan] | Upper Aerodigestive Tract; Carcinoma; Squamous Cell Carcinoma |
| **ESS2** | ['p.?', 'p.G121S', nan] | Breast; Carcinoma; Luminal NS Carcinoma |
| **FAM32A** | [nan] | Central nervous system; Primitive neuroectodermal tumour- |

| | | |
|---|---|---|
| | | medulloblastoma; Classic |
| **FAM50A** | ['p.S276T', nan] | Lung; Carcinoma; Small Cell Carcinoma |
| **FRA10AC1** | ['p.T78R', nan] | Soft tissue; Gastrointestinal stromal tumour |
| **FUS** | ['p.G225S', 'p.S366L', 'p.F437=', 'p.?', 'p.G228_G230dup', 'p.S438F', 'p.R474C', 'p.P151S', 'p.G500S', 'p.K313Q', 'p.R488C', 'p.S440F', 'p.A10T', 'p.K311Q', 'p.G229_G231del', 'p.R377W', 'p.S368L', 'p.P152S', 'p.G221S', 'p.C429S', 'p.R216H', 'p.E278K', 'p.G228_G230del', 'p.A278T', 'p.R473C', 'p.D490=', 'p.G485V', 'p.G505S', 'p.G170=', 'p.G292=', 'p.G266=', 'p.D491=', 'p.Q179H', 'p.R244C', 'p.R378W', 'p.G224S', 'p.P432S', 'p.F438=', 'p.Q178H', 'p.S439F', 'p.P150S', 'p.R213C', 'p.G291=', 'p.R521H', 'p.G503S', 'p.R522H', 'p.S367L', 'p.P430S', 'p.G267=', 'p.G222S', 'p.P150=', 'p.R393H', 'p.G498S', 'p.P151=', 'p.G290=', 'p.E277K', 'p.R486C', 'p.R376W', 'p.G497V', 'p.R243C', 'p.R395H', 'p.R487C', 'p.R520H', 'p.G229_G231dup', 'p.F439=', 'p.C427S', 'p.G495V', 'p.R212C', 'p.R472C', 'p.G499S', 'p.R394H', 'p.P431S', 'p.G486V', 'p.G487V', 'p.G504S', 'p.G169=', 'p.K312Q', 'p.G496V', 'p.D489=', 'p.C428S', nan] | Skin; Carcinoma; Merkel Cell Carcinoma |
| **GEMIN2** | [nan] | Central nervous system; Primitive neuroectodermal tumour-medulloblastoma; Classic |
| **GEMIN5** | ['p.T244S', 'p.R682Q', 'p.?', nan] | Liver; Carcinoma; Hepatocellular carcinoma |
| **GPATCH1** | ['p.L728S', nan] | Soft tissue; Gastrointestinal stromal tumour |
| **GPKOW** | [nan] | Central nervous system; Primitive neuroectodermal tumour-medulloblastoma; Classic |
| **HNRNPA0** | [nan] | Central nervous system; Primitive neuroectodermal tumour- |

| | | |
|---|---|---|
| | | medulloblastoma; Large Cell |
| **HNRNPA3** | [nan] | Central nervous system; Primitive neuroectodermal tumour-medulloblastoma; Classic |
| **HNRNPAB** | [nan] | Central nervous system; Primitive neuroectodermal tumour-medulloblastoma; Classic |
| **HNRNPC** | ['p.K206N', 'p.K193N', 'p.?', 'p.N272Y', 'p.N192Y', 'p.N258Y', 'p.N259Y', 'p.K150N', 'p.N271Y', 'p.K126N', 'p.N216Y', nan] | Lung; Carcinoma; Small Cell Carcinoma |
| **HNRNPD** | [nan] | Central nervous system; Primitive neuroectodermal tumour-medulloblastoma; Large Cell |
| **HNRNPH1** | ['p.?', nan] | Liver; Carcinoma; Hepatocellular carcinoma |
| **HNRNPH3** | ['p.?', nan] | Liver; Carcinoma; Hepatocellular carcinoma |
| **HNRNPL** | ['p.P367=', 'p.P234=', 'p.?', 'p.P238=', 'p.P371=', nan] | Liver; Carcinoma; Hepatocellular carcinoma |
| **HNRNPLL** | ['p.E119Q', 'p.E114Q', nan] | Liver; Carcinoma; Hepatocellular carcinoma |
| **HNRNPM** | ['p.K214I', 'p.K175I', nan] | Large Intestine; Carcinoma; Adenocarcinoma |
| **HNRNPR** | ['p.M1?', 'p.N571S', 'p.N470S', 'p.N568S', 'p.N530S', 'p.?', 'p.N429S', nan] | Haematopoietic and Lymphoid Tissue; Lymphoid Neoplasm; Acute Lymphoblastic T Cell Leukaemia |
| **HNRNPU** | ['p.D83H', nan] | Haematopoietic and Lymphoid Tissue; Haematopoietic Neoplasm; Chronic |

| | | |
|---|---|---|
| | | Myelomonocytic Leukaemia |
| **HNRNPUL 1** | ['p.?', 'p.A278V', 'p.Y250C', 'p.A378V', 'p.G636E', 'p.E407Vfs*17', 'p.E198Dfs*31', 'p.G536E', 'p.P700=', 'p.E307Vfs*17', 'p.A110S', 'p.E293Vfs*17', 'p.E298Dfs*31', 'p.E318Vfs*17', 'p.P711=', 'p.P710=', 'p.P800=', 'p.G547E', 'p.Y150C', 'p.P696=', 'p.A264V', 'p.G522E', 'p.A289V', 'p.Y161C', 'p.E209Dfs*31', 'p.Y207C', nan] | Liver; Carcinoma; Hepatocellular carcinoma |
| **HNRNPUL 2** | ['p.N650=', nan] | Lung; Carcinoma; Adenocarcinoma |
| **HSPA8** | ['p.?', 'p.P344=', 'p.Q424P', 'p.N141S', 'p.S286F', 'p.L210F', 'p.K539R', 'p.V195M', 'p.P108=', 'p.K393R', 'p.K520R', 'p.S2F', 'p.G560D', 'p.G414D', 'p.Q188P', 'p.P325=', 'p.Q405P', 'p.G324D', 'p.S50F', 'p.V176M', 'p.G541D', 'p.S267F', 'p.K303R', 'p.P198=', 'p.Q278P', 'p.L191F', 'p.S140F', nan] | Lung; Carcinoma; Small Cell Carcinoma |
| **HTATSF1** | ['p.R618I', 'p.?', nan] | Lung; Carcinoma; Small Cell Carcinoma |
| **IK** | ['p.F384V', 'p.F69V', nan] | Large Intestine; Carcinoma; Adenocarcinoma |
| **ISY1** | [nan] | Central nervous system; Primitive neuroectodermal tumour-medulloblastoma |
| **LENG1** | ['p.?', nan] | Skin; Malignant melanoma; Nodular |
| **LSM1** | [nan] | Central Nervous system; Primitive neuroectodermal tumour-medulloblastoma; Classic |
| **LSM2** | ['p.?', 'p.A85=', 'p.A86V', nan] | Liver; Carcinoma; Hepatocellular carcinoma |
| **LSM4** | [nan] | Central nervous system; Primitive neuroectodermal tumour-medulloblastoma; Desmoplastic |
| **LSM5** | ['p.G13R', 'p.G42R', nan] | Liver; Carcinoma; Hepatocellular carcinoma |

| | | |
|---|---|---|
| **LSM6** | [nan] | Central nervous system; Primitive neuroectodermal tumour-medulloblastoma; Classic |
| **LSM7** | ['p.?'] | Upper Aerodigestive Tract; Carcinoma; Squamous Cell Carcinoma |
| **LSM8** | [nan] | Central nervous system; Primitive neuroectodermal tumour-medulloblastoma; Desmoplastic |
| **LUC7L3** | [nan] | Central nervous system; Primitive neuroectodermal tumour-medulloblastoma; Classic |
| **MAGOHB** | ['p.E73K', 'p.E119K', nan] | Breast; Carcinoma; Ductal Carcinoma |
| **MFAP1** | ['p.R153H', nan] | Liver; Carcinoma; Hepatocellular carcinoma |
| **MTREX** | ['p.?', 'p.P720T', nan] | Lung; Carcinoma; Small Cell Carcinoma |
| **NOSIP** | [nan] | Central nervous system; Primitive neuroectodermal tumour-medulloblastoma; Classic |
| **NUDT21** | [nan] | Central nervous system; Primitive neuroectodermal tumour-medulloblastoma; Large Cell |
| **NXF1** | ['p.L212R', 'p.Q473*', 'p.S137R', 'p.?', 'p.R71*', 'p.V387Sfs*32', 'p.R515G', 'p.K22*', 'p.L75R', nan] | Haematopoietic and Lymphoid Tissue; Lymphoid Neoplasm; Chronic Lymphocytic Leukaemia-Small Lymphocytic |

| | | |
|---|---|---|
| | | Lymphoma |
| **NXT1** | [nan] | Breast; Carcinoma; Ductal Carcinoma |
| **PCBP2** | [nan] | Central nervous system; Primitive neuroectodermal tumour-medulloblastoma; Classic |
| **PDCD7** | ['p.A296E', nan] | Liver; Carcinoma; Hepatocellular carcinoma |
| **PHF5A** | ['p.R57C', nan] | Breast; Carcinoma; Luminal NS Carcinoma |
| **PLRG1** | [nan] | Central nervous system; Primitive Neuroectodermal tumour-medulloblastoma; Classic |
| **PPIE** | ['p.?', 'p.E282G', nan] | Breast; Carcinoma; Ductal Carcinoma |
| **PPIG** | ['p.N699D', 'p.L309F', 'p.S181=', 'p.D445E', 'p.L324F', 'p.N684D', 'p.D430E', 'p.S166=', nan] | Soft tissue; Gastrointestinal Stromal Tumour |
| **PPIH** | [nan] | Central nervous system; Primitive neuroectodermal tumour-medulloblastoma; Classic |
| **PPIL1** | [nan] | Central nervous system; Primitive neuroectodermal tumour-medulloblastoma; Classic |
| **PPIL2** | [nan] | Central nervous system; Primitive neuroectodermal tumour-medulloblastoma; Classic |
| **PPIL3** | ['p.?', nan] | Liver; Carcinoma; Hepatocellular carcinoma |

| | | |
|---|---|---|
| **PPP1R8** | ['p.?', 'p.K142Q', nan] | Liver; Carcinoma; Hepatocellular carcinoma |
| **PPWD1** | ['p.A454G', 'p.?', 'p.A328G', 'p.A484G', 'p.T577_V578delinsI', 'p.T451_V452delinsI', 'p.L166=', 'p.L322=', 'p.L292=', 'p.T607_V608delinsI', nan] | Large Intestine; Carcinoma; Adenocarcinoma |
| **PQBP1** | ['p.R129W', 'p.R224W', nan] | Large Intestine; Carcinoma; Adenocarcinoma |
| **PRCC** | ['p.K442E', 'p.Y364C', 'p.E466*', 'p.P136S', 'p.?', 'p.V233M', 'p.P81=', 'p.V232A', 'p.S159A', 'p.E468G', 'p.A353V', 'p.P303L', 'p.G449D', 'p.D341G', nan] | Breast; Carcinoma; Ductal Carcinoma |
| **PRKRIP1** | ['p.?', 'p.A141S', 'p.A84S', 'p.E54K', 'p.S7A', nan] | Liver; Carcinoma; Hepatocellular carcinoma |
| **PRPF18** | ['p.V143=', 'p.?', nan] | Central nervous system;glioma; Astrocytoma Grade IV |
| **PRPF19** | ['p.?', 'p.K56Qfs*26', 'p.T287P', nan] | Liver; Carcinoma; Hepatocellular carcinoma |
| **PRPF31** | ['p.N144K', nan] | Liver; Carcinoma; Hepatocellular carcinoma |
| **PRPF38A** | ['p.?', 'p.D49=', nan] | Liver; Carcinoma; Hepatocellular carcinoma |
| **PRPF4** | ['p.D292N', 'p.G62E', 'p.T267A', 'p.T268A', 'p.D293N', 'p.G63E', nan] | Breast; Carcinoma; Ductal Carcinoma |
| **PRPF6** | ['p.E348=', 'p.?', 'p.R31Q', nan] | Liver; Carcinoma; Hepatocellular carcinoma |
| **PRPF8** | ['p.Y1687=', 'p.R934C', 'p.R474C', 'p.T1025=', 'p.?', 'p.R1112H', 'p.K2098=', 'p.H2196=', 'p.R8Q', 'p.R565W', 'p.L1836W', 'p.Q1337=', 'p.T968=', 'p.R414C', 'p.N297=', 'p.S2060G', 'p.Y431*', 'p.T2219=', 'p.I1662V', 'p.R1094H', 'p.P949=', 'p.R861S', 'p.T918A', 'p.E1292D', 'p.A543V', 'p.R820C', 'p.S1343=', 'p.P828S', 'p.V906I', 'p.L213=', 'p.R2266C', 'p.R1935H', 'p.P1313=', 'p.R1354C', 'p.L238I', 'p.N1804=', 'p.L638=', nan] | Large Intestine; Carcinoma; Adenocarcinoma |
| **PUF60** | ['p.?'] | Liver; Carcinoma; Hepatocellular carcinoma |
| **RALY** | ['p.A214_G215insS', 'p.G251S', 'p.G232S', 'p.G248S', 'p.A230_G231insS', 'p.G235S', nan] | Soft tissue;Gastrointestinal stromal tumour |
| **RALYL** | ['p.?', 'p.A77S', 'p.A90S', nan] | Liver; Carcinoma; |

| | | |
|---|---|---|
| | | Hepatocellular carcinoma |
| **RBM10** | ['p.W678*', 'p.R649*', 'p.R46C', 'p.D681Efs*36', 'p.Y797Gfs*68', 'p.R698Q', 'p.?', 'p.K550Rfs*77', 'p.R83W', 'p.W490Vfs*11', 'p.E755*', 'p.E132*', 'p.R94W', 'p.H367Y', 'p.Q674K', 'p.A97T', 'p.G441V', 'p.*996Lext*43', 'p.Q190*', 'p.Q707*', 'p.K711N', 'p.R309M', 'p.Y504*', 'p.Q245*', 'p.P566Afs*18', 'p.A386P', 'p.S329del', 'p.T436Nfs*10', 'p.S115Tfs*84', 'p.R739C', 'p.G409C', 'p.Q432*', 'p.K889Sfs*23', 'p.E47Afs*81', 'p.E832*', 'p.R740C', 'p.G43Wfs*27', 'p.Q148*', 'p.E770*', 'p.I284_Q286dup', 'p.T299=', 'p.E138K', 'p.N222I', 'p.E674G', 'p.S2P', 'p.A382Vfs*104', 'p.Q263*', 'p.R146K', 'p.A690V', 'p.R147L', 'p.Q349*', 'p.S846L', 'p.I811M', 'p.M809I', 'p.M1?', 'p.T316M', 'p.I348S', 'p.E47*', 'p.W756*', 'p.D820Sfs*35', 'p.R815Hfs*33', 'p.G326C', 'p.E889K', 'p.E686*', 'p.T293Nfs*10', 'p.N585Kfs*10', 'p.R608L', 'p.G965V', 'p.L104*', 'p.R149Q', 'p.R107C', 'p.E534*', 'p.A838V', 'p.V241I', 'p.S557*', 'p.A465Gfs*37', 'p.N594Ifs*33', 'p.S692Afs*32', 'p.Q742*', 'p.G762D', 'p.L763R', 'p.P194Afs*30', 'p.G267C', 'p.G91Vfs*98', 'p.A166S', 'p.D531V', 'p.S306*', 'p.T294Sfs*113', 'p.G55D', 'p.E329Sfs*2', 'p.E264Sfs*2', 'p.R739H', 'p.T808M', 'p.R152C', 'p.R579C', 'p.S687_A688delins*', 'p.V99Afs*27', 'p.K568N', 'p.E735*', 'p.G763C', 'p.Q447*', 'p.R153G', 'p.S90L', 'p.H544Ifs*3', 'p.S661I', 'p.E546*', 'p.L382P', 'p.E231*', 'p.G650*', 'p.R37Q', 'p.I697_R698insTL', 'p.Q707Afs*12', 'p.M788I', 'p.R84W', 'p.P506=', 'p.W252*', 'p.Q674*', 'p.V569A', 'p.Q200*', 'p.F150L', 'p.H289Afs*14', 'p.R255H', 'p.F734Sfs*31', 'p.E215*', 'p.Q790*', 'p.Q71L', 'p.D634Ifs*90', 'p.E353*', 'p.Y838*', 'p.R48W', 'p.V277E', 'p.A331V', 'p.Q148R', 'p.N330S', 'p.H367Afs*14', 'p.S60*', 'p.R563C', 'p.A883V', 'p.H289Y', 'p.T174M', 'p.E606K', 'p.N516Qfs*35', 'p.P477S', 'p.S723Pfs*14', 'p.L363P', 'p.R607Efs*36', 'p.S126Cfs*64', 'p.E800*', 'p.R61M', 'p.E560*', 'p.H566Rfs*25', 'p.A553Hfs*73', 'p.G203*', 'p.N709Tfs*15', 'p.K549Rfs*77', 'p.N728Kfs*10', 'p.A365T', 'p.E658*', 'p.Y596*', 'p.G690V', 'p.P648=', 'p.Q563*', 'p.S102Rfs*87', 'p.D731V', 'p.E894*', 'p.E812K', 'p.Y661*', 'p.L363Sfs*115', 'p.P354S', 'p.G184C', 'p.G261Vfs*5', 'p.K638Gfs*3', 'p.R42C', 'p.G477E', 'p.T809M', 'p.S619Afs*150', 'p.G326Vfs*5', 'p.I353N', 'p.H467D', 'p.K699*', 'p.R726*', 'p.C298F', 'p.D139N', 'p.E112Afs*81', 'p.N253S', 'p.S323F', 'p.E52*', 'p.S268Cfs*64', 'p.L121Pfs*68', 'p.R387M', 'p.Q274*', | thyroid; Carcinoma |

'p.E67*', 'p.S687*', 'p.S414*', 'p.G389Afs*52',
'p.K459Nfs*168', 'p.D365N', 'p.H708P', 'p.R836Xfs*?',
'p.Q628H', 'p.D85N', 'p.K705*', 'p.E751*', 'p.K715Vfs*4',
'p.P647S', 'p.S720L', 'p.R5Dfs*129', 'p.D139E',
'p.E184del', 'p.I418N', 'p.S313Vfs*87', 'p.L285P',
'p.E732D', 'p.D641*', 'p.S218*', 'p.G855V', 'p.Q523*',
'p.P409H', 'p.A247P', 'p.G922Afs*29', 'p.H699R',
'p.P497S', 'p.E63D', 'p.G296Afs*111', 'p.R168Q',
'p.A287T', 'p.A305Vfs*104', 'p.P725=', 'p.I472Tfs*76',
'p.P507=', 'p.I775_R776insTL', 'p.E670*', 'p.Q414*',
'p.K133Sfs*56', 'p.E278*', 'p.I697V', 'p.I271S', 'p.A239T',
'p.Q517L', 'p.S548=', 'p.I953Sfs*36', 'p.G857Afs*29',
'p.S415=', 'p.R648*', 'p.R111C', 'p.R295*', 'p.R775W',
'p.E352*', 'p.Q779*', 'p.H609D', 'p.G297V', 'p.G344C',
'p.Q290H', 'p.D816=', 'p.Y879_R880delins*', 'p.L246*',
'p.R228Q', 'p.I276M', 'p.W110*', 'p.V419=', 'p.E720D',
'p.D281N', 'p.Q178*', 'p.K646N', 'p.T552I', 'p.R740H',
'p.S274F', 'p.A389P', 'p.S738Rfs*63', 'p.R42Ffs*92',
'p.Q426*', 'p.W425*', 'p.Q168*', 'p.Y737_R738delins*',
'p.L428Sfs*115', 'p.R5*', 'p.E100K', 'p.R230*', 'p.Y438*',
'p.R776Q', 'p.I427_Q429dup', 'p.E422K', 'p.R70Dfs*129',
'p.R765*', 'p.W140*', 'p.R149W', 'p.R318H',
'p.D603Efs*36', 'p.Q526K', 'p.K837Sfs*10', 'p.H289P',
'p.F848=', 'p.S622L', 'p.S50Tfs*84', 'p.S797Lfs*13',
'p.N586Kfs*10', 'p.W477C', 'p.R150del', 'p.R86Q',
'p.F591Sfs*31', 'p.R776W', 'p.V16I', 'p.D498G',
'p.D310N', 'p.R714W', 'p.R332H', 'p.R786L', 'p.G857V',
'p.Q789*', 'p.W793_K794delinsCR', 'p.Q485H',
'p.S555Rfs*71', 'p.S324F', 'p.A323Gfs*37', 'p.A740V',
'p.D216N', 'p.S660Rfs*63', 'p.Q197*', 'p.R319H',
'p.E485K', 'p.L921Q', 'p.E420*', 'p.R749Sfs*20',
'p.N593Ifs*33', 'p.R32*', 'p.E732Q', 'p.Q850*', 'p.G409A',
'p.A818V', 'p.S545*', 'p.R6H', 'p.K915Tfs*5', 'p.R48Q',
'p.H849Y', 'p.R698W', 'p.K569N', 'p.G467Afs*52',
'p.R563Wfs*12', 'p.Q92Efs*10', 'p.D640G', 'p.R744C',
'p.D20N', 'p.S360*', 'p.A554Tfs*41', 'p.R831P',
'p.Q772Afs*12', 'p.E184dup', 'p.A773V', 'p.Q448*',
'p.Q64*', 'p.Q485*', 'p.S633Kfs*10', 'p.E154=', 'p.F410C',
'p.S369Nfs*68', 'p.A631Tfs*41', 'p.R699Q', 'p.P790=',
'p.A244P', 'p.G261C', 'p.Q267*', 'p.G766R', 'p.D808V',
'p.R420Wfs*12', 'p.A304Vfs*104', 'p.L198=', 'p.V202M',
'p.W335C', 'p.S43*', 'p.R830*', 'p.S645Pfs*14', 'p.Q630*',
'p.R684Efs*36', 'p.E663*', 'p.K637Gfs*3', 'p.S797L',
'p.F592Sfs*31', 'p.E665*', 'p.S218L', 'p.T493Vfs*134',
'p.G905D', 'p.D295Y', 'p.E296=', 'p.V599Nfs*168',
'p.I840_R841insTL', 'p.E513*', 'p.I353M', 'p.L685V',
'p.R102Q', 'p.A409V', 'p.G584V', 'p.A247T', 'p.Q208*',
'p.Q342*', 'p.E748*', 'p.E177*', 'p.W678Dfs*37',

'p.Q638*', 'p.Q574*', 'p.E795Q', 'p.S288*', 'p.Q739K',
'p.E805*', 'p.Q255*', 'p.S660I', 'p.I810Sfs*36', 'p.E177K',
'p.G345*', 'p.E190V', 'p.S545_A546delins*',
'p.I362_Q364dup', 'p.Q772*', 'p.R990H', 'p.Q773*',
'p.N671Ifs*33', 'p.R171W', 'p.V456Nfs*168', 'p.Q668K',
'p.Q51*', 'p.Y519*', 'p.S164*', 'p.Q486*', 'p.R295G',
'p.E671*', 'p.L827V', 'p.V354=', 'p.G76S', 'p.G542E',
'p.Q915*', 'p.R749Efs*36', 'p.Q571*', 'p.E641*',
'p.S554Afs*150', 'p.G184Vfs*5', 'p.G22S', 'p.L428P',
'p.E451*', 'p.Q767Rfs*18', 'p.Q234*', 'p.D604Efs*36',
'p.E703Rfs*66', 'p.Q628*', 'p.I285_Q287dup', 'p.R21H',
'p.F347L', 'p.E816G', 'p.E182K', 'p.E611*',
'p.P501Afs*18', 'p.A313V', 'p.Q558*', 'p.G840C',
'p.H784Y', 'p.I840V', 'p.E578*', 'p.S406=', 'p.A324T',
'p.H777R', 'p.R420C', 'p.I775V', 'p.R689P', 'p.P896S',
'p.Y647*', 'p.Y814_R815delins*', 'p.G623R', 'p.G153S',
'p.I418M', 'p.W821Dfs*37', 'p.S633Rfs*71', 'p.R186*',
'p.E561Rfs*66', 'p.E672*', 'p.Y505*', 'p.D296Y',
'p.C686W', 'p.G538E', 'p.D185E', 'p.D35_Y36delinsGC',
'p.K637Vfs*4', 'p.E627*', 'p.E117K', 'p.Q481L',
'p.A247S', 'p.E808Q', 'p.D364N', 'p.N736Ifs*33',
'p.L292F', 'p.R677M', 'p.W679Dfs*37', 'p.G449S',
'p.E750*', 'p.A324S', 'p.A618G', 'p.R103W', 'p.R887*',
'p.L828P', 'p.G396E', 'p.E560Rfs*66', 'p.K458Nfs*168',
'p.K562*', 'p.P409Rfs*218', 'p.H707L', 'p.R929L',
'p.R787L', 'p.Q597*', 'p.N515Qfs*35', 'p.D854Tfs*32',
'p.T436Sfs*113', 'p.E357*', 'p.R689C', 'p.P916L',
'p.V457Nfs*168', 'p.E576*', 'p.V569I', 'p.E589Q',
'p.P620S', 'p.Y537*', 'p.T350I', 'p.G616C',
'p.T371Nfs*10', 'p.M732I', 'p.E2Dfs*80', 'p.V492I',
'p.I811Sfs*36', 'p.E242K', 'p.T298=', 'p.R153*',
'p.E45Kfs*2', 'p.E578del', 'p.V16F', 'p.S483=', 'p.Q277*',
'p.G615C', 'p.E154*', 'p.W490*', 'p.P198S', 'p.F292L',
'p.S632Kfs*10', 'p.A422T', 'p.P644T', 'p.F771=',
'p.R421Wfs*12', 'p.Q668Lfs*101', 'p.Y36Ifs*90',
'p.R744*', 'p.I276N', 'p.H432Y', 'p.M731I', 'p.R37L',
'p.A308S', 'p.S803Rfs*63', 'p.V99I', 'p.E605K', 'p.E673*',
'p.N663Kfs*10', 'p.L240P', 'p.G727*', 'p.S477Afs*150',
'p.E513del', 'p.K747Sfs*23', 'p.E665Q', 'p.M930I',
'p.G693C', 'p.L685Pfs*41', 'p.S447Nfs*68',
'p.R660Gfs*46', 'p.S710Kfs*10', 'p.R109*',
'p.L286Sfs*115', 'p.P753S', 'p.R607L', 'p.W334C',
'p.E609*', 'p.A696V', 'p.Y732Gfs*68', 'p.Q203*',
'p.E954K', 'p.V491I', 'p.W679*', 'p.I329Tfs*76',
'p.E829*', 'p.Q714*', 'p.E112D', 'p.I810M', 'p.R397H',
'p.R864L', 'p.G514S', 'p.S193T', 'p.Q213L', 'p.F412L',
'p.R817H', 'p.K718N', 'p.E638Rfs*66', 'p.*853Lext*43',
'p.S318Tfs*88', 'p.S703L', 'p.E703*', 'p.Q739*',

| |
|---|
| 'p.Q832Rfs*18', 'p.A612V', 'p.I348F', 'p.E800Rfs*67', 'p.E344K', 'p.T441=', 'p.T251M', 'p.E125V', 'p.C156F', 'p.P355S', 'p.Q595L', 'p.L635Rfs*88', 'p.R836S', 'p.S775Kfs*10', 'p.V277=', 'p.S203Cfs*64', 'p.D497G', 'p.S693Afs*32', 'p.R743C', 'p.Q265*', 'p.E119dup', 'p.E666*', 'p.S846*', 'p.R535M', 'p.G900V', 'p.E533*', 'p.S391Vfs*87', 'p.E186D', 'p.S704L', 'p.R150Q', 'p.P773L', 'p.W821*', 'p.F175Xfs*88', 'p.D677Sfs*35', 'p.V492A', 'p.R461H', 'p.S704*', 'p.S556Rfs*71', 'p.S466F', 'p.V161M', 'p.T376=', 'p.A474V', 'p.E435*', 'p.S476N', 'p.R452M', 'p.R148W', 'p.L263=', 'p.E121D', 'p.D441N', 'p.V534Nfs*168', 'p.V125M', 'p.E345K', 'p.Y360*', 'p.E435del', 'p.E242Gfs*24', 'p.G763D', 'p.S401F', 'p.K601Nfs*168', 'p.N364I', 'p.Q123*', 'p.P722T', 'p.E751G', 'p.E667Kfs*37', 'p.S619N', 'p.R848H', 'p.K621*', 'p.R97*', 'p.V634I', 'p.D777Tfs*32', 'p.Y760*', 'p.G410A', 'p.Q339L', 'p.K275Sfs*56', 'p.E658Rfs*67', 'p.E280*', 'p.G92D', 'p.G439Afs*111', 'p.S416F', 'p.L686Pfs*41', 'p.S232L', 'p.P336Afs*30', 'p.A696Hfs*73', 'p.K780Vfs*4', 'p.W347*', 'p.G280*', 'p.F669Sfs*31', 'p.Q338L', 'p.H290Y', 'p.R699W', 'p.Y654Gfs*68', 'p.Q339*', 'p.D777V', 'p.E296*', 'p.A564T', 'p.E589*', 'p.R84Q', 'p.S295*', 'p.P831S', 'p.C764W', 'p.P555S', 'p.E403*', 'p.S781L', 'p.R880Hfs*33', 'p.E732Kfs*37', 'p.D575G', 'p.G532Afs*52', 'p.E430*', 'p.S108*', 'p.R421C', 'p.L317P', 'p.R853W', 'p.K760Sfs*10', 'p.P648S', 'p.R396H', 'p.I407Tfs*76', 'p.Q148L', 'p.E698*', 'p.Q629*', 'p.E117*', 'p.G792*', 'p.E190*', 'p.Q350*', 'p.G701R', 'p.Y615*', 'p.G547V', 'p.Q525*', 'p.D223*', 'p.E408K', 'p.D777Ifs*90', 'p.D100_Y101delinsGC', 'p.K133Qfs*6', 'p.C687W', 'p.I413F', 'p.R725Gfs*46', 'p.S541N', 'p.E215K', 'p.H290P', 'p.L713Rfs*88', 'p.G439V', 'p.D609V', 'p.P423Afs*18', 'p.P424Afs*18', 'p.R498Wfs*12', 'p.L260Sfs*71', 'p.E897*', 'p.S179Rfs*87', 'p.E494*', 'p.R42H', 'p.H849L', 'p.S460Tfs*88', 'p.Q225*', 'p.G624R', 'p.E100*', 'p.E280K', 'p.Y761*', 'p.R255G', 'p.R817C', 'p.Q701*', 'p.E888*', 'p.R107Ffs*92', 'p.I271_Q272delinsK', 'p.R168W', 'p.E165Gfs*24', 'p.T371Sfs*113', 'p.P851L', 'p.M720V', 'p.G922V', 'p.R766C', 'p.P340S', 'p.A79T', 'p.G625V', 'p.H565Rfs*25', 'p.E662*', 'p.G668Afs*56', 'p.D881=', 'p.H467Ifs*3', 'p.R5Dfs*80', 'p.R85del', 'p.E763*', 'p.Q320*', 'p.I145=', 'p.S476Afs*150', 'p.E47Afs*32', 'p.W412C', 'p.A156T', 'p.I698V', 'p.R397G', 'p.E67Dfs*129', 'p.H432P', 'p.S414=', 'p.H784L', 'p.Q822*', 'p.W282*', 'p.Y680*', 'p.Q596*', 'p.S646Pfs*14', 'p.E184D', 'p.S252del', 'p.G649*', |

'p.Q283*', 'p.G509S', 'p.Q807*', 'p.G822V', 'p.R715W',
'p.G371S', 'p.M874I', 'p.K201E', 'p.Y101C',
'p.F734Lfs*35', 'p.G780Afs*29', 'p.D300N', 'p.P645T',
'p.D363N', 'p.D746Efs*36', 'p.E187Sfs*2',
'p.W348Vfs*11', 'p.R163Q', 'p.G552A', 'p.D873V',
'p.Y361*', 'p.P754S', 'p.H290Afs*14', 'p.A755V',
'p.A390V', 'p.W756Dfs*37', 'p.Q509*', 'p.S781*',
'p.I698_R699insTL', 'p.Q660L', 'p.T427I', 'p.E344G',
'p.Q665*', 'p.I271F', 'p.E498*', 'p.K838Tfs*5', 'p.E875D',
'p.E735Rfs*67', 'p.E487K', 'p.S295L', 'p.Q689*',
'p.D281E', 'p.A688V', 'p.W871_K872delinsCR', 'p.I68=',
'p.R71H', 'p.W425Vfs*11', 'p.A312V', 'p.Y538*',
'p.G23*', 'p.V267M', 'p.H708Rfs*25', 'p.E590Kfs*37',
'p.S940Lfs*13', 'p.D635Ifs*90', 'p.R745*', 'p.Q419*',
'p.D674V', 'p.R925H', 'p.F913=', 'p.A499T',
'p.Q234Efs*10', 'p.W348*', 'p.H700R', 'p.Y508*',
'p.E608*', 'p.S703*', 'p.V491A', 'p.F770=', 'p.R749K',
'p.E745*', 'p.R831C', 'p.E666Q', 'p.E732*', 'p.E748K',
'p.G811Afs*56', 'p.E436del', 'p.K536Nfs*168',
'p.L636Rfs*88', 'p.G762C', 'p.E823*', 'p.G473E',
'p.P271Afs*30', 'p.E119del', 'p.L293F', 'p.Q603Lfs*101',
'p.Q213R', 'p.R657C', 'p.K124E', 'p.E495*', 'p.D635V',
'p.Q689Rfs*18', 'p.Y903*', 'p.K627Rfs*77', 'p.D223N',
'p.R688*', 'p.R792W', 'p.E813*', 'p.S698Rfs*71',
'p.V241Afs*27', 'p.G218S', 'p.Q630Afs*12', 'p.R5Kfs*7',
'p.R81K', 'p.K764*', 'p.E667*', 'p.H842R', 'p.S661Rfs*63',
'p.E589Kfs*37', 'p.G779Afs*29', 'p.*931Lext*43',
'p.R687*', 'p.R759S', 'p.E733D', 'p.D678Sfs*35',
'p.A174T', 'p.A553Tfs*41', 'p.E667Q', 'p.R882H',
'p.Q332*', 'p.D784*', 'p.G779V', 'p.I210=',
'p.P610Lfs*159', 'p.Q225H', 'p.Y504C', 'p.S125*',
'p.A554Hfs*73', 'p.R841Q', 'p.K759Sfs*10', 'p.Q664*',
'p.H544D', 'p.M787I', 'p.A321P', 'p.E436*', 'p.R103Q',
'p.A430T', 'p.N787Tfs*15', 'p.S244Rfs*87',
'p.T294Nfs*10', 'p.G296V', 'p.P467Lfs*159',
'p.K902Sfs*10', 'p.E112*', 'p.S492=', 'p.F268C',
'p.G778V', 'p.Y518*', 'p.E119D', 'p.A324P', 'p.S415*',
'p.Q272*', 'p.S317Tfs*88', 'p.M66I', 'p.C233F', 'p.A455V',
'p.E938Q', 'p.I413S', 'p.W678G', 'p.A695V', 'p.Q590*',
'p.R86H', 'p.D216E', 'p.S49=', 'p.E189D', 'p.E628*',
'p.E292*', 'p.Y503*', 'p.Y36Ifs*41', 'p.S554N', 'p.V303M',
'p.Q92*', 'p.S798Lfs*13', 'p.S370Nfs*68',
'p.P545Lfs*159', 'p.S360L', 'p.K746Sfs*23', 'p.R758S',
'p.R70*', 'p.T492I', 'p.T636Vfs*134', 'p.E550K',
'p.E590*', 'p.L778Q', 'p.R901S', 'p.K837Tfs*5',
'p.G157D', 'p.H609Ifs*3', 'p.T694I', 'p.Y101Ifs*90',
'p.H706L', 'p.G390Afs*52', 'p.D438Y', 'p.E683*',
'p.D438A', 'p.Q526Lfs*101', 'p.M865I', 'p.L195Xfs*71',

'p.P649=', 'p.H466Ifs*3', 'p.A389T', 'p.Q690Rfs*18',
'p.K210Sfs*56', 'p.Q686*', 'p.A617G', 'p.Q596K',
'p.A447Vfs*104', 'p.E657*', 'p.H432Afs*14', 'p.S545L',
'p.W187*', 'p.E743Q', 'p.D311N', 'p.C829W', 'p.S687L',
'p.S351F', 'p.S719L', 'p.Y582*', 'p.K88*', 'p.V634A',
'p.R498C', 'p.G374Afs*111', 'p.D453N', 'p.L285Sfs*115',
'p.R102L', 'p.F270L', 'p.L685R', 'p.E355*', 'p.L685P',
'p.S353*', 'p.S512Nfs*68', 'p.A243S', 'p.R822*',
'p.L286P', 'p.R80C', 'p.G669Afs*56', 'p.E2Dfs*129',
'p.T551I', 'p.S862L', 'p.E796Q', 'p.D365*', 'p.R612M',
'p.L686R', 'p.S622*', 'p.Q767*', 'p.Q649*', 'p.G384Vfs*6',
'p.L181*', 'p.R145C', 'p.L686P', 'p.P408Rfs*218',
'p.Q629Afs*12', 'p.P787T', 'p.E705*', 'p.G400E',
'p.S211*', 'p.S50Tfs*35', 'p.V176I', 'p.D642*', 'p.D300*',
'p.A389S', 'p.E590Q', 'p.Q544*', 'p.L763Pfs*41',
'p.Y36C', 'p.Y655Gfs*68', 'p.Q480*', 'p.Q496*',
'p.E723*', 'p.D506N', 'p.P275S', 'p.E138*', 'p.E683K',
'p.Q148H', 'p.E345G', 'p.G307Vfs*6', 'p.P408H',
'p.Q525Lfs*101', 'p.R98Efs*36', 'p.N710Tfs*15',
'p.P584=', 'p.S25F', 'p.G165A', 'p.Q543*', 'p.L763P',
'p.F345C', 'p.L684V', 'p.Q27K', 'p.Q138*', 'p.Q573*',
'p.F669Lfs*35', 'p.G306Vfs*6', 'p.W217*', 'p.E743*',
'p.K230*', 'p.S875Lfs*13', 'p.W794_K795delinsCR',
'p.S606N', 'p.E128D', 'p.S477N', 'p.Q524*', 'p.V238M',
'p.M655V', 'p.N658Qfs*35', 'p.G442V', 'p.L195Sfs*71',
'p.Q484*', 'p.R42Ffs*43', 'p.T494Vfs*134', 'p.R847H',
'p.E429*', 'p.Q844*', 'p.A695G', 'p.G746Afs*56',
'p.R607Sfs*20', 'p.Q495*', 'p.D388N', 'p.E721D',
'p.E746*', 'p.A631Hfs*73', 'p.E811K', 'p.P774L',
'p.P486Rfs*218', 'p.W679G', 'p.R294C', 'p.N852Tfs*15',
'p.Q867*', 'p.Q690*', 'p.D919Tfs*32', 'p.K980Tfs*5',
'p.S394del', 'p.I330Tfs*76', 'p.S463N', 'p.R580C',
'p.Q285*', 'p.E606*', 'p.R230G', 'p.E740*',
'p.I348_Q349delinsK', 'p.R821C', 'p.Q572*', 'p.Y582C',
'p.Y647C', 'p.L121=', 'p.A205V', 'p.D755Sfs*35',
'p.Q702*', 'p.L370F', 'p.S770Afs*32', 'p.E620*',
'p.S788Pfs*14', 'p.E657Rfs*67', 'p.G100A', 'p.D373Y',
'p.V176Afs*27', 'p.Q597K', 'p.K266E', 'p.R70Kfs*7',
'p.Q341*', 'p.K576N', 'p.S544L', 'p.R857W', 'p.I953M',
'p.R583Gfs*46', 'p.E873Q', 'p.S6P', 'p.G652S', 'p.H707Y',
'p.R737Hfs*33', 'p.R606Efs*36', 'p.E561*', 'p.E242*',
'p.P432S', 'p.Q621*', 'p.E351*', 'p.Q431*', 'p.F592Lfs*35',
'p.E865*', 'p.Q71R', 'p.E254D', 'p.E863D', 'p.A741V',
'p.I413_Q414delinsK', 'p.G519V', 'p.Q525K',
'p.G233Vfs*98', 'p.D712V', 'p.G372S', 'p.D738=',
'p.Q138Sfs*3', 'p.L435F', 'p.S492*', 'p.R685L', 'p.A288T',
'p.P478S', 'p.T293Sfs*113', 'p.A760G', 'p.K563*',
'p.F591Lfs*35', 'p.R886C', 'p.L762V', 'p.R841W',

| | | |
|---|---|---|
| | 'p.R688P', 'p.K780Gfs*3', 'p.G758C', 'p.G777V', 'p.E722*', 'p.K715Gfs*3', 'p.P486H', 'p.G780V', 'p.G395E', 'p.E752*', 'p.Q666*', 'p.Q623*', 'p.E798D', 'p.G823V', 'p.D730V', 'p.Q932*', 'p.G920V', 'p.R766P', 'p.N395S', 'p.S544*', 'p.G510S', 'p.F227L', 'p.K622*', 'p.R606Sfs*20', 'p.S395Tfs*88', 'p.D634V', 'p.S128T', 'p.Q563H', 'p.A140V', 'p.Q50*', 'p.T571Vfs*134', 'p.R684Sfs*20', 'p.Q486H', 'p.G297Afs*111', 'p.E125*', 'p.E638*', 'p.L856Q', 'p.R738Hfs*33', 'p.Q757*', 'p.P725S', 'p.R882C', 'p.S167L', 'p.P790S', 'p.H706Y', 'p.Q680*', 'p.D719*', 'p.R791*', 'p.N593Qfs*35', 'p.R750L', 'p.A613V', 'p.Q416L', 'p.L263Pfs*68', 'p.K165*', 'p.*854Lext*43', 'p.S405=', 'p.V419E', 'p.H643Rfs*25', 'p.E499*', 'p.D739=', 'p.Q405*', 'p.S241*', 'p.D532V', 'p.Q129*', 'p.G548V', 'p.Q603K', 'p.R163Efs*36', 'p.K692Rfs*77', 'p.R918W', 'p.Q679*', 'p.E407K', 'p.E277*', 'p.E422G', 'p.D120E', 'p.A696Tfs*41', 'p.S622_A623delins*', 'p.S557=', 'p.K638Vfs*4', 'p.R82L', 'p.S456Vfs*87', 'p.Q481*', 'p.K275Qfs*6', 'p.L198Pfs*68', 'p.Q116*', 'p.A11D', 'p.E676*', 'p.L828R', 'p.E605*', 'p.W821G', 'p.D296A', 'p.L118Sfs*71', 'p.L778Rfs*88', 'p.R107H', 'p.E481*', 'p.R310M', 'p.D712Ifs*90', 'p.E621*', 'p.Q601*', 'p.D776Tfs*32', 'p.T349I', 'p.V354E', 'p.A322Gfs*37', 'p.T951M', 'p.M66Ifs*4', 'p.Q518L', 'p.G449Vfs*6', 'p.Q832*', 'p.D373A', 'p.E754*', 'p.S314Vfs*87', 'p.P468Lfs*159', 'p.Q290*', 'p.A221T', 'p.R688C', 'p.N299I', 'p.G399E', 'p.G840D', 'p.P551Rfs*218', 'p.Q284*', 'p.M578V', 'p.H367P', 'p.A400Gfs*37', 'p.S738I', 'p.I888Sfs*36', 'p.Q340*', 'p.Y505C', 'p.S544_A545delins*', 'p.G168Vfs*98', 'p.G587S', 'p.R229C', 'p.R722C', 'p.P551H', 'p.H466D', 'p.R582Gfs*46', 'p.K210Qfs*6', 'p.Q361*', 'p.Q310*', 'p.K575N', 'p.E404*', 'p.L828Pfs*41', 'p.Q169Efs*10', 'p.R534M', 'p.P647=', 'p.W347Vfs*11', 'p.T629I', 'p.A421T', 'p.W936_K937delinsCR', 'p.G374V', 'p.E231=', 'p.E810D', 'p.R236W', 'p.E28K', 'p.S464N', 'p.D489N', 'p.G487A', 'p.L779Q', 'p.K653N', 'p.M577V', 'p.K640*', 'p.T886M', 'p.R251*', 'p.G905C', 'p.D295A', 'p.Y736_R737delins*', 'p.W756G', 'p.R332G', 'p.Q169*', 'p.A332V', 'p.R85Q', 'p.I888M', 'p.K824Sfs*23', 'p.S803I', 'p.E307Gfs*24', 'p.E815*', 'p.Q203Sfs*3', 'p.E673G', 'p.E808*', 'p.E487G', 'p.S835Afs*32', nan] | |
| **RBM17** | ['p.?', nan] | Liver; Carcinoma; Hepatocellular carcinoma |
| **RBM22** | ['p.M338I', 'p.G339*', 'p.G388*', 'p.M387I', nan] | Kidney; Carcinoma; Papillary Renal Cell |

| | | |
|---|---|---|
| | | Carcinoma |
| **RBM25** | ['p.E404Tfs*33', 'p.K527=', 'p.?', 'p.R409=', 'p.P49T', 'p.A266=', 'p.P643=', nan] | Haematopoietic and Lymphoid Tissue; Lymphoid Neoplasm; Plasma Cell Myeloma |
| **RBM28** | ['p.?', 'p.Q715L', 'p.Q574L', nan] | Liver; Carcinoma; Hepatocellular carcinoma |
| **RBM5** | ['p.G729S', 'p.G518D', nan] | Soft Tissue; fibrosarcoma |
| **RBM8A** | ['p.G41R'] | Soft tissue;Gastrointestinal stromal tumour |
| **RBMX2** | [nan] | Central nervous system; Primitive neuroectodermal tumour-medulloblastoma; Classic |
| **RBMXL2** | ['p.A4V', 'p.A66V', 'p.T134A', nan] | Soft Tissue; Gastrointestinal Stromal Tumour |
| **RNF113A** | ['p.A28T', 'p.E88D', nan] | Breast; Carcinoma; Ductal Carcinoma |
| **RNPC3** | ['p.N358S', 'p.?', 'p.P109=', 'p.P474A', 'p.T67Nfs*8', 'p.K368N', 'p.M160I', 'p.P415L', 'p.P473A', 'p.P4S', 'p.N359S', 'p.P414L', 'p.K367N', nan] | Ovary; Carcinoma; Mucinous Carcinoma |
| **RNPS1** | [nan] | Central nervous system; Primitive neuroectodermal tumour-medulloblastoma; Classic |
| **SAP18** | [nan] | Central nervous system; Primitive neuroectodermal tumour-medulloblastoma |
| **SART1** | ['p.G485A', 'p.?', nan] | Haematopoietic and Lymphoid Tissue; Lymphoid Neoplasm; T Cell Large Granular Lymphocytic Leukaemia |
| **SART3** | ['p.?', 'p.Y758H', 'p.Y794H', 'p.A336T', nan] | Liver; Carcinoma; Hepatocellular carcinoma |

| | | |
|---|---|---|
| **SDE2** | ['p.?', 'p.E321*', nan] | Liver; Carcinoma; Hepatocellular carcinoma |
| **SF3A1** | ['p.T299S', 'p.R511W', 'p.V586G', 'p.?', 'p.I159V', 'p.E331K', 'p.P558L', 'p.R699H', 'p.T482A', 'p.V648L', 'p.P618L', 'p.A528D', 'p.R511Q', 'p.E75K', 'p.N74K', 'p.R390H', nan] | Haematopoietic and Lymphoid Tissue; Haematopoietic Neoplasm; Acute Myeloid Leukaemia |
| **SF3A2** | [nan] | Central nervous system; Primitive neuroectodermal tumour-medulloblastoma |
| **SF3B1** | ['p.I556T', 'p.K80N', 'p.Q191*', 'p.I15V', 'p.?', 'p.K503R', 'p.L1247=', 'p.A672V', 'p.D476Y', 'p.D393Y', 'p.A706V', 'p.P370L', 'p.E312K', 'p.R625H', 'p.D856V', 'p.Q41H', 'p.Y570C', 'p.R702Dfs*12', 'p.R827K', 'p.K666E', 'p.E902Q', 'p.G1162R', 'p.G740X', 'p.P646S', 'p.A708T', 'p.Q220R', 'p.I704S', 'p.L478I', 'p.R124Q', 'p.R1180*', 'p.R315*', 'p.Q949H', 'p.P1243=', 'p.Q670E', 'p.D476H', 'p.S229=', 'p.E622G', 'p.R625L', 'p.F444=', 'p.L1129=', 'p.N804H', 'p.W658R', 'p.L575=', 'p.H662Y', 'p.R315Q', 'p.E902K', 'p.D302E', 'p.S286F', 'p.Q699_K700del', 'p.A920V', 'p.E1205K', 'p.R1045W', 'p.E1221Q', 'p.S685N', 'p.N626Y', 'p.A52T', 'p.T935K', 'p.R495I', 'p.E622V', 'p.A188S', 'p.Q14H', 'p.Y1284=', 'p.N626I', 'p.P87S', 'p.S73F', 'p.I875T', 'p.P130S', 'p.R924*', 'p.G664C', 'p.R625P', 'p.M367V', 'p.V701Sfs*14', 'p.E988K', 'p.M620V', 'p.S75N', 'p.H662D', 'p.K666R', 'p.R939C', 'p.G83*', 'p.L539dup', 'p.L90=', 'p.H8Y', 'p.Y623C', 'p.I667V', 'p.D894G', 'p.P1257T', 'p.L1280=', 'p.T766N', 'p.L600M', 'p.H1184=', 'p.H662Q', 'p.R1297C', 'p.Q19E', 'p.P380A', 'p.R390I', 'p.R1041H', 'p.G751V', 'p.W658S', 'p.H738Y', 'p.N291Y', 'p.G347V', 'p.V695L', 'p.D883N', 'p.D907Vfs*13', 'p.D68N', 'p.P489S', 'p.E9K', 'p.G305V', 'p.D725N', 'p.R625G', 'p.K22_A23delinsNL', 'p.K946T', 'p.F1081S', 'p.G264R', 'p.R196Q', 'p.G740E', 'p.N941S', 'p.S1051Y', 'p.Q699_K700delinsHE', 'p.Q699E', 'p.D856N', 'p.Y928C', 'p.E776A', 'p.L1024*', 'p.L1065H', 'p.N626S', 'p.N626D', 'p.D67=', 'p.A52V', 'p.K666N', 'p.R132C', 'p.C965S', 'p.K666M', 'p.G330E', 'p.K748M', 'p.D1290N', 'p.K666T', 'p.Y141C', 'p.I360Sfs*22', 'p.I704N', 'p.W658C', 'p.F777L', 'p.A1072=', 'p.M784_K785delinsI', 'p.L536F', 'p.E1033D', 'p.K741E', 'p.Y1136*', 'p.E688K', 'p.K741*', 'p.R594Q', 'p.E906K', 'p.R625C', 'p.N626H', 'p.S1150Ifs*7', 'p.D885Y', 'p.V701F', 'p.R451*', 'p.A745P', 'p.D1291H', 'p.A744P', 'p.G83=', 'p.A660V', 'p.S637F', 'p.E760K', | Prostate; Carcinoma; Adenocarcinoma |

'p.Q901R', 'p.L674P', 'p.E776G', 'p.P409Q', 'p.A678T', 'p.A284T', 'p.G742D', 'p.A1279V', 'p.E27K', 'p.Q1186H', 'p.Y864C', 'p.A176D', 'p.L467I', 'p.S611F', 'p.A672S', 'p.P780R', 'p.P355S', 'p.S129F', 'p.I1287M', 'p.R702Gfs*27', 'p.W1266*', 'p.D907Y', 'p.H306Y', 'p.D894Y', 'p.S1150L', 'p.V1219=', 'p.R939H', 'p.E1054K', 'p.K785R', 'p.A263G', 'p.L1132_M1133insQL', 'p.K22N', 'p.M784I', 'p.R132H', 'p.G53K', 'p.D144N', 'p.G797E', 'p.G1146R', 'p.P780T', 'p.Q1248*', 'p.V576L', 'p.S400F', 'p.R318*', 'p.I704V', 'p.K6E', 'p.T663I', 'p.L161F', 'p.S1124L', 'p.A1229V', 'p.L639=', 'p.K963N', 'p.D781E', 'p.Y623H', 'p.K666Q', 'p.E783_K785del', 'p.P813L', 'p.Q903R', 'p.L747W', 'p.R495S', 'p.A263T', 'p.P297L', 'p.V701I', 'p.T964A', 'p.K700E', 'p.K649E', 'p.D1278Y', 'p.L897I', 'p.Y898H', 'p.S229Y', 'p.V482I', 'p.R517H', 'p.E873K', 'p.Q931E', 'p.E64K', 'p.R630S', 'p.K182E', 'p.V576A', 'p.D65N', 'p.I704F', 'p.D781G', 'p.K653_S657delinsT', 'p.D799G', 'p.P224S', 'p.R451Q', 'p.Y421F', 'p.K748E', 'p.K653Q', 'p.K741N', 'p.A277V', 'p.D894N', 'p.E64D', 'p.C1123Y', 'p.E300Q', 'p.E902V', 'p.R775G', 'p.E622D', 'p.G676Afs*11', 'p.E968Kfs*3', 'p.E860K', 'p.I875V', 'p.V577del', 'p.S705dup', 'p.V635L', 'p.R736C', 'p.E1061A', 'p.I690T', 'p.G877A', 'p.E889K', 'p.L581=', 'p.G676D', 'p.0', 'p.R590K', 'p.P372S', 'p.W338L', 'p.I850V', 'p.Q949Nfs*6', 'p.R702W', 'p.E243K', 'p.L1182F', 'p.V1169I', 'p.V634I', 'p.R397H', 'p.R238H', 'p.R1286G', 'p.I787Lfs*3', 'p.A364V', 'p.P718L', 'p.P145R', 'p.P363S', 'p.E1135K', 'p.P1139S', 'p.D584N', 'p.R397C', 'p.S643C', 'p.P642S', 'p.L897V', 'p.W293R', 'p.D835N', 'p.A385=', 'p.S705_A706dup', 'p.L415P', 'p.V577=', 'p.P304L', 'p.E783K', 'p.T627P', 'p.A861T', 'p.G283=', 'p.N291I', 'p.L994I', 'p.Q1277*', 'p.A749T', 'p.V591M', 'p.M446I', 'p.A711V', 'p.P615S', 'p.L756F', 'p.R625S', 'p.K700delinsRVWEVRE', 'p.H1091R', 'p.G605S', 'p.G880R', 'p.A711D', 'p.Y623F', 'p.K1071N', 'p.Y561F', 'p.E592K', 'p.Q931H', 'p.R702_I704del', 'p.P212L', 'p.S229F', 'p.M453V', 'p.L773R', 'p.I704T', 'p.R238C', 'p.I1272V', 'p.C1204R', 'p.G271D', 'p.D781N', 'p.L399F', 'p.K141=', 'p.S637C', 'p.K741T', 'p.G1275A', 'p.I721F', 'p.A1085V', 'p.E776D', 'p.A1052Kfs*9', 'p.K943E', 'p.R425Q', 'p.P327L', 'p.M147I', 'p.P465S', 'p.I253V', 'p.W1266R', 'p.G972A', 'p.T703P', 'p.K1102I', 'p.D1291del', 'p.R625X', 'p.I673F', 'p.E595D', 'p.V184I', 'p.Q863del', 'p.G282R', 'p.E491G', 'p.T299A', 'p.P755S', 'p.G740R', 'p.S944A', 'p.D586H', 'p.I787N', 'p.A419P', 'p.E545Q', 'p.Q670Q', 'p.R1074C', 'p.A176V', 'p.E579_P580delinsA', 'p.D483Y', 'p.D546Y', 'p.P1224T',

| | | |
|---|---|---|
| | 'p.R512I', 'p.G605D', 'p.E809Dfs*7', 'p.P569S', 'p.E1160K', 'p.R157Q', 'p.T350S', 'p.K522R', 'p.A1076T', 'p.T663P', 'p.W658L', 'p.W232*', 'p.R1019Tfs*10', 'p.R196*', 'p.K666X', 'p.D618Y', 'p.L982=', 'p.Q534P', 'p.R549C', 'p.I787Nfs*21', 'p.P530L', 'p.G740V', 'p.N1270K', 'p.L502S', 'p.R775Q', 'p.R775L', 'p.P1171L', 'p.M609I', nan] | |
| **SF3B2** | ['p.?', 'p.R146W', 'p.E313=', 'p.E296=', 'p.P120S', 'p.S326F', 'p.A774V', 'p.A757V', 'p.S343F', nan] | Skin; Carcinoma; Merkel Cell Carcinoma |
| **SF3B3** | ['p.A632T', 'p.G60C', 'p.A892D', 'p.S659L', nan] | Lung; Carcinoma; Small Cell Carcinoma |
| **SF3B4** | [nan] | Central nervous system; Primitive neuroectodermal tumour-medulloblastoma; Classic |
| **SF3B5** | [nan] | Breast; Carcinoma; Ductal Carcinoma |
| **SF3B6** | [nan] | Central nervous system; Primitive neuroectodermal tumour-medulloblastoma; Classic |
| **SFSWAP** | ['p.R855W', 'p.R907W', 'p.G512S', 'p.L421P', nan] | Lung; Carcinoma; Small Cell Carcinoma |
| **SMNDC1** | [nan] | Central nervous system; Primitive neuroectodermal tumour-medulloblastoma |
| **SMU1** | ['p.?', nan] | Liver; Carcinoma; Hepatocellular carcinoma |
| **SNIP1** | [nan] | Central nervous system; Primitive neuroectodermal tumour-medulloblastoma; Large Cell |
| **SNRNP200** | ['p.L1788P', 'p.?', 'p.P1723=', 'p.Q64*', 'p.F1736L', nan] | Stomach; Carcinoma; Intestinal Adenocarcinoma |
| **SNRNP25** | [nan] | Central nervous system; Primitive |

| | | |
|---|---|---|
| | | neuroectodermal tumour-medulloblastoma; Classic |
| **SNRNP40** | [nan] | Central nervous system; Primitive neuroectodermal tumour-medulloblastoma; Classic |
| **SNRNP48** | ['p.P45L', nan] | Soft tissue;Gastrointestinal stromal tumour |
| **SNRNP70** | [nan] | Central nervous system; Primitive neuroectodermal tumour-medulloblastoma; Classic |
| **SNRPA** | ['p.I93Lfs*6', 'p.G99D', nan] | Liver; Carcinoma; Hepatocellular carcinoma |
| **SNRPA1** | ['p.R219=', 'p.L70*', 'p.?', nan] | Lung; Carcinoma; Small Cell Carcinoma |
| **SNRPB** | ['p.?', nan] | Liver; Carcinoma; Hepatocellular carcinoma |
| **SNRPB2** | ['p.K19Q', 'p.?', nan] | Large Intestine; Carcinoma; Adenocarcinoma |
| **SNRPD1** | [nan] | Central nervous system; Primitive neuroectodermal tumour-medulloblastoma; Large Cell |
| **SNRPD2** | [nan] | Central nervous system; Primitive neuroectodermal tumour-medulloblastoma; Large Cell |
| **SNRPD3** | [nan] | Central nervous system; Primitive neuroectodermal tumour-medulloblastoma; |

| | | |
|---|---|---|
| | | Desmoplastic |
| **SNRPE** | [nan] | Central nervous system; Primitive neuroectodermal tumour-medulloblastoma; Large Cell |
| **SNRPF** | ['p.?', nan] | Liver; Carcinoma; Hepatocellular carcinoma |
| **SREK1** | ['p.?', nan] | Liver; Carcinoma; Hepatocellular carcinoma |
| **SRRM1** | ['p.?', nan] | Liver; Carcinoma; Hepatocellular carcinoma |
| **SRRM2** | ['p.?', 'p.G360D', 'p.P804T', 'p.S2020=', 'p.A308=', 'p.R206*', 'p.S2020F', 'p.P2373=', 'p.A41D', 'p.A137D', 'p.R40C', 'p.G264D', 'p.E1134=', 'p.L71=', 'p.S1474A', 'p.R136C', 'p.K121del', 'p.R2396K', 'p.K768=', 'p.R302*', 'p.P897L', 'p.S1550=', 'p.A212=', 'p.R1944H', 'p.K217del', 'p.T1856I', 'p.R1776H', nan] | Urinary Tract; Carcinoma |
| **SRSF1** | ['p.P89S', 'p.?', nan] | Breast; Carcinoma; Ductal Carcinoma |
| **SRSF10** | ['p.?', nan] | Liver; Carcinoma; Hepatocellular carcinoma |
| **SRSF11** | ['p.S281C', 'p.S221C', nan] | Breast; Carcinoma; Luminal NS Carcinoma |
| **SRSF12** | [nan] | Central nervous system; Primitive neuroectodermal tumour-medulloblastoma; Classic |
| **SRSF2** | ['p.R94=', 'p.F62C', 'p.P95H', 'p.V87L', 'p.K211Sfs*21', 'p.Y92H', 'p.D97Gfs*20', 'p.P201F', 'p.P95_H99del', 'p.S179F', 'p.R103_G114del', 'p.A78_G82del', 'p.E69Q', 'p.G4D', 'p.M1?', 'p.P96L', 'p.S169T', 'p.Y115Tfs*7', 'p.S132R', 'p.G93delinsDR', 'p.P95T', 'p.R108C', 'p.S98*', 'p.S132_R143del', 'p.P96dup', 'p.S171F', 'p.A166S', 'p.?', 'p.G93dup', 'p.P95Xfs*2', 'p.H63Qfs*169', 'p.S142F', 'p.S54F', 'p.G93_H100del', 'p.H99L', 'p.P107S', 'p.S169Vfs*44', 'p.R125H', 'p.P95_R117del', 'p.R94_P95del', 'p.E53K', 'p.P95S', 'p.R186_S191del', 'p.D9N', 'p.T25S', 'p.R66C', 'p.D67N', 'p.P95A', 'p.S183F', | Haematopoietic and Lymphoid Tissue; Haematopoietic Neoplasm; Acute Myeloid Leukaemia |

| | | |
|---|---|---|
| | 'p.S173F', 'p.P96=', 'p.E214Kfs*18', 'p.P209S', 'p.D19G', 'p.R129L', 'p.H63Qfs*157', 'p.R155Q', 'p.A154S', 'p.E214Kfs*30', 'p.R94_S101del8', 'p.P107H', 'p.P95_D97del', 'p.Y44H', 'p.S118F', 'p.K199Sfs*21', 'p.S130F', 'p.R94_P106del', 'p.D97Gfs*8', 'p.R103Pfs*149', 'p.P95L', 'p.S101Rfs*9', 'p.G77E', 'p.L16F', 'p.P197S', 'p.E215K', 'p.P95R', 'p.R113H', 'p.V60Afs*183', 'p.S169F', 'p.R103Pfs*125', 'p.S196F', 'p.R127C', 'p.D19Y', 'p.Q88E', 'p.H63Qfs*181', 'p.K211Sfs*33', 'p.R167L', 'p.S157F', 'p.E203K', 'p.S202F', 'p.K197N', 'p.S161F', 'p.V87Gfs*35', 'p.S151P', 'p.S147W', 'p.Y92_H99del', 'p.S204L', 'p.V87Gfs*23', 'p.G104Dfs*140', 'p.S157Vfs*44', 'p.F57Y', 'p.V18L', 'p.R155L', 'p.S183Y', 'p.G104Dfs*116', 'p.P96S', 'p.E69K', 'p.V60Afs*171', 'p.G12D', 'p.0', 'p.G93_R94insRVQMARYG', 'p.S184F', 'p.E202Kfs*18', 'p.P96Rfs*148', 'p.G104Dfs*128', 'p.R66Afs*4', 'p.P95_R102del', 'p.R167Q', 'p.S120_R131del', 'p.S220F', 'p.S194L', 'p.R155Sfs*62', 'p.P8S', 'p.S192L', 'p.S135W', 'p.R94dup', 'p.P96Rfs*136', 'p.S208F', 'p.P95delinsRX', 'p.R167Sfs*62', 'p.R94_P95insR', 'p.R103Pfs*137', 'p.S101Rfs*21', 'p.P213F', 'p.R94P', 'p.S171Y', 'p.S206L', 'p.K185N', 'p.V60Afs*159', 'p.S190F', 'p.R174_S179del', 'p.R115C', 'p.S139P', 'p.P96Rfs*124', 'p.R167Sfs*74', 'p.S167F', 'p.P95Xfs*?', 'p.S157T', 'p.S120R', nan] | |
| **SRSF3** | ['p.?', nan] | Liver; Carcinoma; Hepatocellular carcinoma |
| **SRSF4** | ['p.G338A', 'p.S366N', 'p.G356S', 'p.E253D', 'p.V173A', nan] | Soft tissue; Gastrointestinal stromal tumour |
| **SRSF5** | ['p.R81Q', 'p.R84Q', nan] | Large Intestine; Carcinoma; Adenocarcinoma |
| **SRSF6** | ['p.R145Q', 'p.?', nan] | Large Intestine; Carcinoma; Adenocarcinoma |
| **SRSF7** | ['p.?', nan] | Liver; Carcinoma; Hepatocellular carcinoma |
| **SRSF8** | ['p.S222P', 'p.H189D'] | Large Intestine; Carcinoma; Adenocarcinoma |
| **SUGP1** | ['p.R290H', 'p.?', 'p.A121S', 'p.P389=', nan] | Soft tissue; Gastrointestinal stromal tumour |
| **TFIP11** | ['p.?', nan] | Liver; Carcinoma; |

| | | |
|---|---|---|
| | | Hepatocellular carcinoma |
| **THOC1** | ['p.Q584R', 'p.?', 'p.D370Afs*16', nan] | Soft tissue; Gastrointestinal stromal tumour |
| **THOC2** | ['p.L601=', 'p.K1336_A1337delinsN', 'p.L486=', 'p.E1335_K1336insRP', 'p.K1221_A1222delinsN', 'p.H55D', 'p.I389del', 'p.I1453del', 'p.K154Gfs*5', 'p.I1568del', 'p.?', 'p.K157_A158delinsN', 'p.V286=', 'p.V171=', 'p.E1220_K1221insRP', 'p.K1333Gfs*5', 'p.K1218Gfs*5', 'p.E156_K157insRP', nan] | Lung; Carcinoma; Adenocarcinoma |
| **THOC3** | [nan] | Central nervous System; Primitive neuroectodermal tumour-medulloblastoma; Large Cell |
| **THOC5** | ['p.?', 'p.G499S', 'p.V515A', 'p.T380K', 'p.L216V', 'p.V525I', 'p.W443*', nan] | Liver; Carcinoma; Hepatocellular carcinoma |
| **THOC6** | ['p.V55I', 'p.V79I', 'p.L219=', 'p.L243=', nan] | Central Nervous System; glioma; Astrocytoma Grade IV |
| **THOC7** | ['p.?', 'p.L23F', 'p.L75F', nan] | Liver; Carcinoma; Hepatocellular carcinoma |
| **THRAP3** | ['p.P905S', 'p.F73=', 'p.P555S', 'p.S783F', 'p.R522*', 'p.K558T', 'p.R837Q', 'p.P273=', 'p.P269=', 'p.K131=', 'p.T564S', 'p.S746=', 'p.P254=', 'p.S32=', 'p.P496Q', 'p.A201V', 'p.?', 'p.Q620*', 'p.W869*', 'p.K711E', 'p.P95S', 'p.D577Y', 'p.S28L', 'p.R608C', 'p.R245W', 'p.S562F', 'p.R89C', 'p.P275L', 'p.G473D', 'p.R23C', 'p.R774=', 'p.Q301*', 'p.S164F', 'p.R718C', 'p.R12C', 'p.S684R', 'p.I679R', 'p.R677K', 'p.S560F', 'p.P395I', nan] | Skin; Carcinoma; Basal Cell Carcinoma |
| **TRA2A** | [nan] | Central nervous system; Primitive neuroectodermal tumour-medulloblastoma;classic |
| **TRA2B** | [nan] | Central nervous system; Primitive neuroectodermal tumour-medulloblastoma |
| **TXNL4A** | [nan] | Central nervous system; Primitive |

| | | |
|---|---|---|
| | | neuroectodermal tumour-medulloblastoma;Large Cell |
| **U2AF1** | ['p.F150I', 'p.R159T', 'p.Q84H', 'p.?', 'p.F77I', 'p.Q157R', 'p.R202C', 'p.R190H', 'p.R209H', 'p.P73L', 'p.D160E', 'p.E51G', 'p.Q84P', 'p.R166Q', 'p.R161C', 'p.E159K', 'p.G214D', 'p.R83H', 'p.L130S', 'p.D107H', 'p.R239*', 'p.P146L', 'p.L57S', 'p.G149_G150del', 'p.R203C', 'p.C154S', 'p.0', 'p.N170S', 'p.G167S', 'p.R165*', 'p.H120R', 'p.R188H', 'p.D233E', 'p.E184K', 'p.K15R', 'p.R35L', 'p.Q157H', 'p.A69=', 'p.R116C', 'p.V23=', 'p.E162Q', 'p.S199F', 'p.A126T', 'p.R92*', 'p.G150*', 'p.Y158_E159dup', 'p.D34H', 'p.C163*', 'p.Y158_E159del', 'p.S34F', 'p.R232K', 'p.R35Q', 'p.Y85_E86del', 'p.R203H', 'p.R129H', 'p.*241Sext*10', 'p.R194S', 'p.R118C', 'p.R46H', 'p.L65W', 'p.G215S', 'p.E91D', 'p.R121S', 'p.R130C', 'p.N170Y', 'p.C82=', 'p.R35G', 'p.E111K', 'p.Q157Xfs*?', 'p.E159Xfs*?', 'p.A47V', 'p.R117H', 'p.Q157G', 'p.R196T', 'p.M99I', 'p.E184Q', 'p.R189C', 'p.V69L', 'p.G212A', 'p.G223*', 'p.R28H', 'p.Q75R', 'p.R153C', 'p.S158L', 'p.F22S', 'p.R66C', 'p.R53C', 'p.L130F', 'p.C81R', 'p.R110W', 'p.K23R', 'p.C155=', 'p.Q84R', 'p.C154R', 'p.Q157P', 'p.R130H', 'p.D31H', 'p.R232T', 'p.G222_G223del', 'p.L57F', 'p.*168Sext*10', 'p.R239Q', 'p.F9Lfs*15', 'p.F82Lfs*15', 'p.Y85_E86dup', 'p.E10Q', 'p.N104=', 'p.R136H', 'p.S7=', 'p.G141D', 'p.R156Q', 'p.C81S', 'p.E184V', 'p.R125*', 'p.M26I', 'p.E124G', 'p.R202H', 'p.R109W', 'p.N31=', 'p.E111Q', 'p.I24T', 'p.G223R', 'p.R234C', 'p.S126F', 'p.E162Mfs*32', 'p.S34Y', 'p.A142=', 'p.Q2R', 'p.R156H', 'p.G150R', 'p.E159A', 'p.G94S', 'p.E18D', 'p.H193R', 'p.R129C', 'p.R159K', 'p.R123T', 'p.N97S', 'p.R45C', 'p.R92Q', 'p.R165Q', 'p.E73Q', 'p.E83Q', 'p.G215Xfs*?', 'p.G139A', 'p.R53H', 'p.G30E', 'p.R119H', 'p.S231L', 'p.R182W', 'p.E86K', 'p.R183W', 'p.G142S', 'p.A53T', 'p.R166*', 'p.E89Mfs*32', 'p.R226C', 'p.N97Y', 'p.R198*', 'p.V96=', 'p.E235Q', 'p.R115H', 'p.E159_M160insYE', 'p.K39*', nan] | Kidney; Carcinoma; Clear Cell Renal Cell Carcinoma |
| **U2AF2** | ['p.G154S', 'p.R284L', 'p.?', 'p.R227H', 'p.E225D', 'p.I191Efs*20', 'p.L187V', 'p.E389D', 'p.G154V', 'p.R452L', 'p.T145I', 'p.E393D', 'p.R63H', 'p.D45N', 'p.S349Efs*28', 'p.R448L', 'p.I27Efs*20', 'p.L23V', nan] | Haematopoietic and Lymphoid Tissue; Haematopoietic Neoplasm; Myelodysplastic Syndrome |
| **U2SURP** | ['p.?', nan] | Liver; Carcinoma; Hepatocellular |

| | | |
|---|---|---|
| | | carcinoma |
| **UPF1** | ['p.?', 'p.D371Mfs*46', 'p.D360Mfs*46', 'p.D108Y', 'p.G147R', 'p.I941=', 'p.I952=', 'p.D389E', 'p.T429M', 'p.D371E', 'p.D360E', 'p.D225H', 'p.L84=', 'p.K311T', 'p.S438C', 'p.K282E', 'p.T440M', 'p.D400E', 'p.S449C', nan] | Breast; Carcinoma; Ductal Carcinoma |
| **USP39** | ['p.K171=', 'p.F187L', 'p.G486R', 'p.T289M', 'p.T392M', 'p.G535R', 'p.G564R', 'p.G461R', 'p.F290L', 'p.?', 'p.K93=', 'p.K68=', 'p.T314M', 'p.F212L', nan] | Breast; Carcinoma; Ductal Carcinoma |
| **WBP11** | ['p.S364C', nan] | Lung; Carcinoma; Adenocarcinoma |
| **WBP4** | ['p.V40=', 'p.K113R', nan] | Kidney; Carcinoma; Granular Cell |
| **WDR83** | ['p.T114=', 'p.?', nan] | Breast; Carcinoma; Basal (triple-negative) Carcinoma |
| **XAB2** | ['p.?', 'p.R138W', 'p.I283=', nan] | Liver; Carcinoma; Hepatocellular carcinoma |
| **YBX1** | [nan] | Central nervous system; Primitive neuroectodermal tumour-medulloblastoma;classic |
| **YJU2** | [nan] | Central nervous system; Primitive neuroectodermal tumour-medulloblastoma |
| **ZMAT2** | [nan] | Central nervous system; Primitive neuroectodermal tumour-medulloblastoma |
| **ZNF830** | ['p.H99Q', 'p.N46D', nan] | Soft tissue; Gastrointestinal stromal tumour |

**Table S3. Cancer-associated Mutations located within IDRs.**

| **Protein** | **Residue** | **Class/ Family** | **Tumor Location** |
|---|---|---|---|
| RBM10 | 1;2;3;4;5;6;7;10;11;12;13;14;15;16;18;20;21;22;23;24;25;27;28;29;30;31;32;33;35;36;37;41;42;43;45;46;47;48;49;50;51;52;55;56;60;61;63;64;66;67;68;70;71;73;76;77;79;80;81 | Recruited at A comple | Thyroid; carcinoma |

| | | | |
|---|---|---|---|
| | ;82;83;84;85;86;87;88;90;91;92;94;97;98;99;100;101;102;103;104;107;108;109;110;111;112;113;115;116;117;118;119;120;121;123;124;125;126;420;495;496;497;498;499;503;504;506;507;508;509;512;513;514;515;516;627;628;629;630;631;632;633;634;635;636;637;638;640;641;642;643;644;645;646;647;648;649;650;661;662;667;668;669;672;679;683;684;685;686;687;688;689;690;692;696;736;739;740;741;742;743;744;745;746;747;748;818;820;821;822;823;824;827;828;829;830;831;832;835;836;837;838;840;841;842;844;846;847;848;849;850;851;852;853;854;855;856;857;871;872;873;874;875;879;880;881;882;883;888;889;896;897;900;901;902;903;905 | x | |
| SF3B1 | 0;3;6;7;8;9;10;11;12;13;14;15;19;21;22;23;27;41;52;53;64;65;67;68;73;75;80;83;87;90;124;129;130;132;141;144;145;147;157;161;176;182;184;188;191;196;212;220;224;229;232;238;243;253;263;264;271;277;282;283;284;286;291;293;297;299;300;302;304;305;306;312;315;318;327;330;338;347;350;355;360;363;364;367;370;372;380;385;390;393;397;399;400;409;415;419;421;425;444;446;451;453 | U2 snRNP | Prostate; carcinoma; adenocarcinoma |
| SRSF2 | 1;95;96;97;98;99;100;101;102;103;104;106;107;108;113;114;115;116;117;118;120;124;125;127;128;129;130;131;132;135;136;137;139;140;142;143;147;148;149;151;154;155;157;159;161;166;167;169;171;173;174;179;181;183;184;185;186;190;191;192;194;196;197;199;201;202;203;204;206;208;209;211;213;214;215;220 | SR related | Haematopoietic and lymphoid tissue; haematopoietic neoplasm; acute myeloid leukaemia |
| FUS | 10;150;151;152;169;170;178;179;212;213;216;221;222;224;225;228;229;230;231;243;244;266;267;277;278;311;312;313;376;377;378;393;394;395;427;428;429;430;431;432;437;438;439;440;472;473;474;485;486;487;488;489;490;491;495;496;497;498;499;500;503;504;505;520;521;522 | hnRNP | Skin; carcinoma; Merkel cell carcinoma |
| U2AF1 | 57;182;183;188;189;190;193;194;196;198;199;202;203;209;212;214;215;222;223;226;231;232;233;234;235;239 | SR protein | Kidney; carcinoma; clear cell renal cell carcinoma |
| THOC2 | 1218;1220;1221;1222;1333;1335;1336;1337;1453;1568 | TREX | Lung; carcinoma; adenocarcinoma |
| RALY | 214;215;230;231;232;235;248;251 | hnRNP | Soft tissue; gastrointestinal stromal tumour |
| PPIG | 181;309;324;430;445;684;699 | U2 | Soft tissue; |

| | | | |
|---|---|---|---|
| | | snRNP | gastrointestinal stromal tumour |
| SRPK1 | 28;44;281;297;325;341 | Linked to splicing | Breast; Carcinoma; ductal carcinoma |
| U2AF2 | 20;23;27;28;45;63 | U2 snRNP associated | Haematopoietic and lymphoid tissue; haematopoietic neoplasm; Myelodysplastic Syndrome |
| DDX23 | 8;33;39;95;247 | U5 snRNP | Breast; carcinoma; ductal carcinoma |
| PRCC | 81;136;159;232;233 | Recruited at Bact complex | Breast; carcinoma; ductal carcinoma |
| SF3A1 | 331;390;511;528;558 | hnRNP | Haematopoietic and lymphoid tissue; haematopoietic neoplasm; acute myeloid leukaemia |
| DDX42 | 24;143;769;888 | U2 snRNP associated | Lung; carcinoma; adenocarcinoma |
| RNPC3 | 4;8;358;359 | U11/ U12 snRNP | Ovary; carcinoma; mucinous carcinoma |
| SRSF4 | 253;338;356;366 | SR protein | Soft tissue; gastrointestinal stromal tumour |
| CPSF4 | 166;199;224 | CPSF/ | Liver; |

| | | | |
|---|---|---|---|
| | | CSTF | carcinoma; hepatocellular carcinoma |
| DDX20 | 536;636;642 | GEM | Upper aerodigestive tract; carcinoma; squamous cell carcinoma |
| GCFC2 | 108;124;211 | Non-specified | Central nervous system; glioma; astrocytoma Grade IV |
| NXF1 | 32;71;75 | EJC/ mRNP | Haematopoietic and lymphoid tissue; lymphoid neoplasm; chronic lymphocytic leukaemia-small lymphocytic lymphoma |
| RBM22 | 338;339;388 | Prp19 related | Kidney; carcinoma; papillary renal cell carcinoma |
| SF3B2 | 120;146;296 | Recruited at B complex | Skin; carcinoma; Merkel cell carcinoma |
| CHERP | 341;352 | U2 snRNP associated | Upper aerodigestive tract; carcinoma; squamous cell carcinoma |
| CWC22 | 742;773 | Recruited at Bact | Soft tissue; gastrointestinal stromal |

| | | | |
|---|---|---|---|
| | | complex | tumour |
| CWC27 | 256;335 | Step II factor | Oesophagus; carcinoma; squamous cell carcinoma |
| DDX41 | 25;133 | Recruited at C complex | Central nervous system; glioma; astrocytoma Grade IV |
| LSM2 | 85;86 | LSM | Liver; carcinoma; hepatocellular carcinoma |
| PQBP1 | 129;224 | Prp19 related | Large intestine; carcinoma; adenocarcinoma |
| RBM5 | 518;729 | Recruited at A complex | Soft tissue; fibrosarcoma |
| SRSF5 | 81;84 | SR protein | Large intestine; carcinoma; adenocarcinoma |
| SRSF8 | 189;222 | SR protein | Large intestine; carcinoma; adenocarcinoma |
| SUGP1 | 121;389 | Recruited at A complex | Soft tissue; gastrointestinal stromal tumour |
| BUD13 | 190 | RES complex | Soft tissue; gastrointestinal stromal tumour |
| CPSF1 | 561 | SR protein | Soft tissue; gastrointestinal stromal |

| | | | |
|---|---|---|---|
| | | | tumour |
| DHX8 | 448 | U5 snRNP | Haematopoietic and Lymphoid Tissue; lymphoid neoplasm; acute lymphoblastic T cell leukaemia |
| ESS2 | 121 | Recruited at C complex | Breast; carcinoma; luminal NS carcinoma |
| MFAP1 | 153 | Recruited at B complex | Liver; carcinoma; hepatocellular carcinoma |
| NONO | 36 | Linked to splicing | Lung; carcinoma; small cell carcinoma |
| PPWD1 | 484 | Recruited at C complex | Large intestine; carcinoma; adenocarcinoma |
| RBM25 | 643 | hnRNP | Haematopoietic and lymphoid tissue; lymphoid neoplasm; plasma cell myeloma |
| RBM28 | 715 | Recruited at Bact complex | Liver; Carcinoma; hepatocellular carcinoma |
| RBM8A | 41 | EJC/ mRNP | Soft tissue; gastrointestinal stromal tumour |
| SDE2 | 321 | U4/U6 snRNP | Liver; Carcinoma; |

| | | | |
|---|---|---|---|
| | | | hepatocellular carcinoma |
| SNRPA | 99 | U1 snRNP | Liver; Carcinoma; hepatocellular carcinoma |
| WBP11 | 364 | Prp19 related | Lung; Carcinoma; adenocarcinoma |
| WBP4 | 113 | Recruited at B complex | Kidney; carcinoma; granular cell |

**Table S5. Enrichment analysis of SPL proteins with PTM sites in IDRs.**

| Name | PTM position | PTM type | Enrichment term |
|---|---|---|---|
| DDX41 | [21, 23] | ['Phosphoserine'] | Abnormality of the skin, Neoplasm |
| FUS | [216, 216, 218, 218, 221, 242, 244, 248, 277, 377, 383, 388, 394, 473, 476, 481, 485, 487, 491, 495, 498, 503, 503] | ['Asymmetric dimethylarginine', 'Omega-N-methylarginine', 'Phosphoserine'] | Abnormality of the skin |
| PRCC | [157, 159] | ['Phosphoserine'] | Neoplasm |
| RBM10 | [61, 89, 736, 845, 902, 2, 30] | ['Phosphoserine', 'Omega-N-methylarginine', 'N-acetylserine'] | Abnormality of the skin |
| RBM8A | [42] | ['Phosphoserine'] | Abnormality of the skin, Neoplasm |
| SF3B1 | [125, 129, 141, 142, 157, 194, 207, 211, 214, 223, 227, 229, 235, 244, 248, 257, 261, 267, 273, 278, 287, 296, 299, 303, 313, 322, 326, 328, 332, 341, 344, 349, 350, 354, 400, 426] | ['Phosphothreonine', 'Phosphoserine', 'N6-acetyllysine', 'Citrulline'] | Abnormality of the skin, Neoplasm |
| SRSF2 | [189, 191, 204, 206, 208, 212, 220] | ['Phosphoserine'] | Abnormality of the skin, Neoplasm |
| THOC2 | [1222, 1450] | ['Phosphoserine'] | Abnormality of |

| | | | |
|---|---|---|---|
| | | | the skin |
| U2AF2 | [15] | ['5-hydroxylysine'] | Abnormality of the skin |
| WBP11 | [361, 364] | ['Phosphoserine'] | Abnormality of the skin |
